# Branched photoswitchable tethered ligands enable ultra-efficient optical control and detection of class C G protein-coupled receptors

**DOI:** 10.1101/563957

**Authors:** Amanda Acosta-Ruiz, Vanessa A. Gutzeit, Mary Jane Skelly, Samantha Meadows, Joon Lee, Anna G. Orr, Kristen Pleil, Johannes Broichhagen, Joshua Levitz

**Affiliations:** Biochemistry, Cell and Molecular Biology Graduate Program, Weill Cornell Medicine, New York, NY 10065; Neuroscience Graduate Program, Weill Cornell Medicine, New York, NY 10065; Department of Pharmacology, Weill Cornell Medicine, New York, NY 10065; Department of Biochemistry, Weill Cornell Medicine, New York, NY 10065; Brain and Mind Research Institute, Weill Cornell Medicine, New York, NY 10065; Appel Alzheimer’s Disease Research Institute, Weill Cornell Medicine, New York, NY 10065; Max Planck Institute for Medical Research, Department of Chemical Biology, Jahnstr. 29, 69120 Heidelberg, Germany; Tri-Institutional PhD Program in Chemical Biology, New York, NY 10065

## Abstract

The limitations of classical, soluble drugs in terms of subtype-specificity, spatiotemporal precision, and genetic targeting have spurred the development of advanced pharmacological techniques, including the use of covalently-tethered photoswitchable ligands. However, a major shortcoming of tethered photopharmacology is the inability to obtain optical control with a comparable efficacy to the native ligand. To overcome the limitations of photoisomerization efficiency and tethered ligand affinity, we have developed a family of branched photoswitchable compounds to target G protein-coupled metabotropic glutamate receptors (mGluRs). These compounds permit photo-agonism of G_i/o_-coupled group II mGluRs with near-complete efficiency relative to saturating glutamate when attached to receptors via a range of orthogonal, multiplexable modalities including SNAP-, CLIP-, and Halo-tags, as well as via receptor-targeting nanobodies. Through a chimeric approach, branched ligands also allow efficient optical control of G_q_-coupled mGluR5 with precise, dynamic subcellular targeting. Finally, branched ligands enabled the development of dual photoswitch-fluorophore compounds that allow simultaneous imaging and manipulation of receptors via the same attachment point. Together this work provides a new design framework for photoswitchable ligands and demonstrates a toolset suitable for quantitative, mechanistic study of neuromodulatory receptors at the molecular, cellular and circuit levels.

## Introduction

Pharmacological studies have provided a foundation for our understanding of the molecular basis of biological function, particularly with regard to membrane receptors^1^. However, shortcomings in the spatiotemporal precision at which drugs can be added and removed from a preparation, the paucity of drugs which unambiguously distinguish between molecular subtypes, and the inability to target drug action to genetically-defined cell types limits the mechanistic insight that can be gleaned from such studies. In recent years, photopharmacology has emerged as an alternative approach where light-dependent chemical cages, such as 4-methoxy-7-nitroindolinyl (MNI), or photoswitches, such as azobenzenes, are conjugated to established compounds to allow light to control ligand efficacy and thus afford the system with improved spatial and temporal precision^2^.

When such chemical photoswitches are covalently tethered to a protein target they facilitate the highest degree of specificity, spatial and temporal control and, through genetic targeting of the protein target itself, allow drug action to effectively be limited to defined cell types within physiological systems^3^. By tethering the ligand to a specific tag, the subtype specificity problem is solved at the level of attachment rather than at the ligand binding site, enabling native ligands to serve as the functional group without concerns about off-target pharmacological effects. Genetic encoding of the target protein can also, in principle, facilitate the incorporation of receptor variants or mutants to test their role in a physiological context. While earlier studies used native nucleophiles^4^ or engineered cysteines^5-7^ or reactive unnatural amino acids^8^ for tethering, we recently introduced photoswitchable, orthogonal, remotely-tethered ligands (“PORTLs”) which attach to a genetically-encoded self-labeling tag (i.e. SNAP), enabling efficient and orthogonal labeling of the protein target^9,10^.

G protein-coupled receptors (GPCRs), the largest family of membrane receptors in eukaryotes and the largest class of drug targets^11,12^, are particularly well-suited to tethered photopharmacology. Many receptor subtypes for the same ligand often exist and due to their highly conserved binding sites, developing specific agonists and antagonists remains a major challenge. Furthermore, the cellular and physiological complexity of GPCR signaling, especially in the nervous system where they signal in distinct spatially-delimited contexts such as the synapse, and in distinct cell types within neural circuits, highlights the limitations of traditional pharmacology. The metabotropic glutamate receptors (mGluRs) form a family of neuromodulatory GPCRs which respond to the excitatory neurotransmitter glutamate to control neuronal excitability and synaptic strength in many different brain regions^13^. These central roles in neuronal signaling have allowed the eight mGluRs to emerge as highly attractive drug targets for disorders ranging from psychiatric and neurological disease to cancer^14^, but deciphering their underlying mechanisms has proven to be challenging, limiting our understanding of both their fundamental signaling properties and the basis for their roles in disease. Azobenzene-glutamate PORTLs were initially developed for N-terminally SNAP-tagged mGluR2^9,10^ where they have been shown to effectively turn mGluR2 signaling on and off with sub-second precision for applications in cultured cells^15^ and in the retina of behaving mice^16^. While this system represents one of the most efficient tethered photoswitch systems characterized to date, many limits still exist including incomplete efficacy of the photoswitch relative to glutamate, the inability to visualize labeled receptors, and the lack of validation of the technique in intact brain tissue. In addition, extending the PORTL approach to other mGluR subtypes has been a major challenge that has so far limited this approach primarily to analysis of mGluR2.

Here we sought to develop strategies to enhance the efficiency and applicability of PORTLs, with a focus on mGluRs. Following mechanistic characterization of the determinants of PORTL efficiency where we discover the importance of flexibility of the self-labeling protein tag, we introduce and characterize a family of branched, tethered azobenzene-glutamate photoswitches that allow for photo-agonism of mGluR2 with near-complete efficacy relative to glutamate. We successfully design and characterize branched PORTLs for SNAP, CLIP, Halo and nanobody-based labeling strategies. The branching approach also allows for high-efficiency optical control of mGluR3, which previously showed minimal light responses with single-branch PORTLs^10^, and of mGluR5 through a chimera strategy. We use this approach for control and spatiotemporal analysis of mGluR5-induced calcium oscillations and demonstrate the use of this tool to control mGluR5 signaling in astrocytes, where we find evidence for confinement of calcium responses to processes. Finally, the branching framework allows for the development of a first generation of dual photoswitchable ligand/fluorophore compounds that permit simultaneous optical control and sensing of mGluR2 in cortical brain slices. Together this work introduces a diverse toolset of multiplexable, photoswitchable mGluRs and a generalizable strategy for enhancing the efficiency of tethered and photoswitchable ligands.

## Results

### SNAP-tag flexibility, not linker composition, is a major determinant of tethered photoswitch efficiency

We initially focused on a mechanistic analysis of the prototypical system of <u>b</u>enzyl<u>g</u>uanine-<u>a</u>zobenzene-<u>g</u>lutamate (“BGAG_n_” where n=number of PEG repeats between BG and azobenzene) conjugated to SNAP-mGluR2 (**Fig. 1a**). We previously analyzed the role of BGAG length and found that a similar 50-60% photoswitching efficiency was observed in compounds ranging from BGAG_0_ to BGAG_12_, but that beyond 12 PEG repeats the efficiency starts to decrease^10^. The lack of clear length-dependence over a wide range raised the questions of what other parameters determine photoswitch efficiency. We probed the role of the distance and orientation between PORTL attachment and ligand binding sites by analyzing the linker between the self-labelling SNAP-tag and the mGluR2 ligand binding domain. We added two, four or six flexible glycine-glycine-serine (“GGS”) repeats or removed the existing threonine-arginine (“TR”) linker along with the C-terminal glycine-leucine-glycine (“GLG”) motif of SNAP. For each construct we determined BGAG_12_ efficiency relative to glutamate by measuring G protein-coupled inward rectifier potassium (GIRK) channel activation in HEK 293T cells using whole cell patch clamp electrophysiology (**Fig. 1b; Fig S1a, b**). The system tolerated either the complete removal of the linker or the addition of up to 4 GGS repeats, but a decrease in efficiency was observed with 6 GGS repeats (**Fig. S1c**). We also tested SNAP-(GGS)_6_-mGluR2 with short (BGAG_0_) and long (BGAG_28_) variants and found that in all cases a clear reduction in efficiency was seen (**Fig. S1d, e**). The tolerance of the system for a wide range of linker lengths means that there likely is not a precise optimal linker or photoswitch length for fine-tuning photoswitch efficiency.

**Figure 1.**
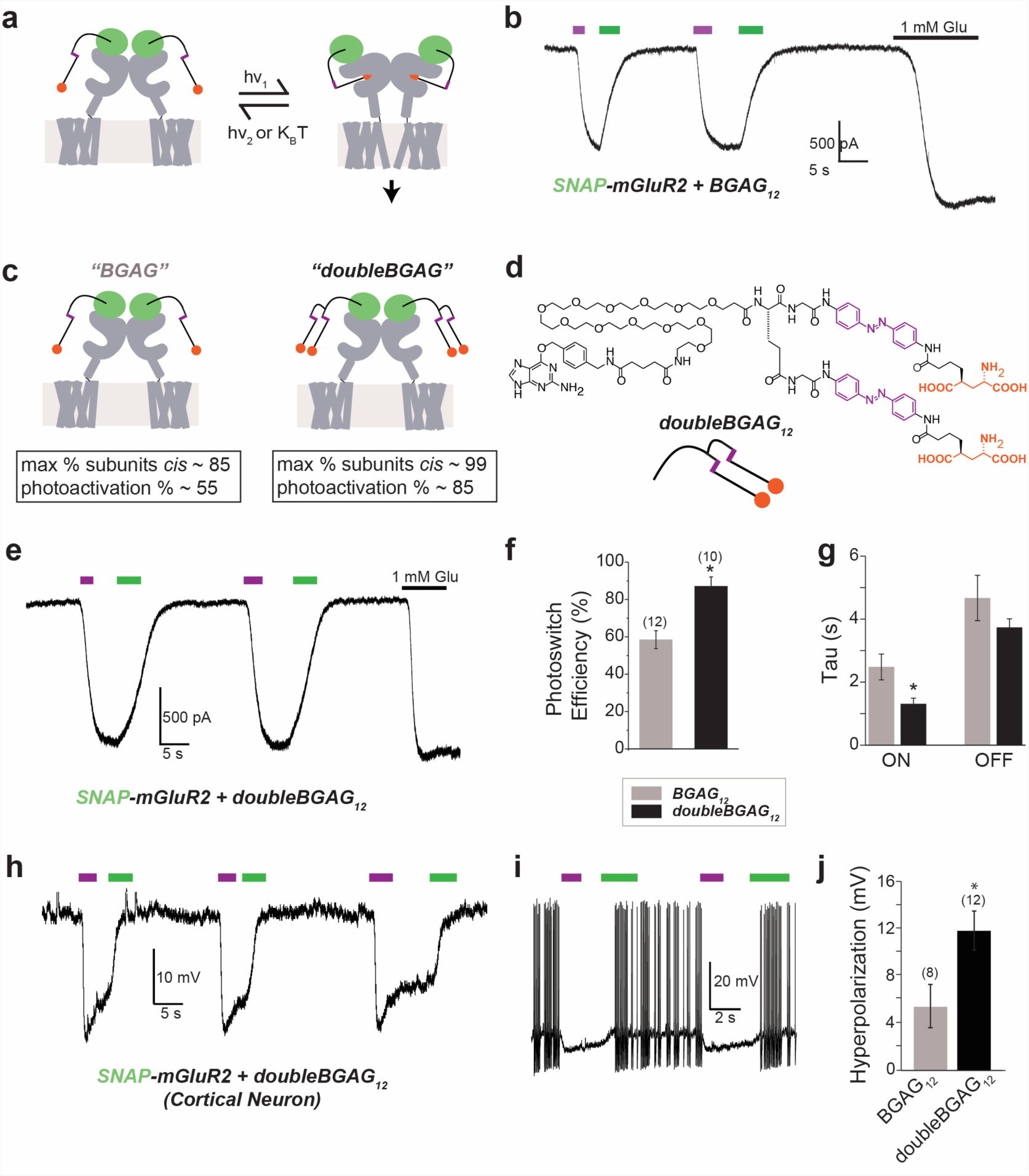
A branched photoswitchable ligand enables high efficiency, rapid photo-activation and de-activation of mGluR2 (**a**) Schematic showing photo-activation of SNAP-tagged mGluR2 with covalently-tethered “BGAG” photoswitches. Full-length wild-type mGluR2 is shown in grey and the genetically-encoded N-terminal SNAP-tag is shown in green. BGAG molecules contain an O^6^-benzylguanine moiety for SNAP labeling, a central azobenzene photoswitch (magenta) and a 4’ tethered L-glutamate (orange circle). (**b**) Representative HEK 293T whole cell patch clamp electrophysiology trace showing BGAG_12_- mediated photoactivation of mGluR2 by 385 nm illumination (magenta bars) and deactivation with 525 nm illumination (green bars) compared to application of saturating 1 mM glutamate. (**c**) Schematic showing branched BGAG concept. The percentage of subunits with an active, *cis*-BGAG is calculated based on azobenzene photostationary states at 385 nm and the photoactivation efficiency is estimated based on the cooperativity of mGluR2 where agonist binding in one subunit activates 20% relative to binding in both subunits (see Fig. S4). (**d**) Chemical structure of doubleBGAG_12_. The azobenzene moieties are shown in magenta and the glutamates are shown in orange. (**e-f**) Photoswitch efficiency is enhanced to near complete efficiency. Representative whole cell patch clamp recording (e) and summary bar graph (f) showing high efficiency photoactivation of SNAP-mGluR2 via doubleBGAG_12_. * indicates statistical significance (unpaired t-test, p=0.0004). (**g**) Summary of kinetics of photo-activation and photo-deactivation of mGluR2 with BGAG_12_ versus doubleBGAG_12_. * indicates statistical significance (unpaired t-test, p=0.03). (**h-j**) doubleBGAG12 enables high-efficiency photoactivation of native effectors via SNAP-mGluR2 in cultured cortical neurons. Representative traces show doubleBGAG_12_-mediated light-induced hyperpolarization (h) and action potential silencing (i) and summary bar graph (j) shows enhanced hyperpolarization for doubleBGAG_12_. * indicates statistical significance (unpaired t-test, p=0.02). The numbers of cells tested are shown in parentheses. Error bars show s.e.m.

We next turned to the SNAP-tag itself as a potential source of photoswitch modulation. SNAP was evolved from a human alkyltransferase and many variants exist with different biophysical and labeling properties^17-21^. All existing SNAP-mGluR photoswitching has been based on the SNAP26m variant which is referred to simply as SNAP in this study. We decided to test the “SNAPfast” (SNAP_f_) variant which is defined by an increased affinity for a broader substrate profile. Surprisingly, BGAG_12_ photoswitch efficiency was drastically reduced with SNAP_f_-mGluR2 (**Fig. S2a, b;** efficiency= 31.9 ± 5.0% for SNAP_f_; unpaired t-test versus SNAP, p=0.0007) and BGAG_0_ switching was nearly abolished (**Fig. S2c, d**). Based on its increased intramolecular interactions^22^, we hypothesized that SNAP_f_ provides a less flexible attachment point which constrains the conformational landscape of BGAG limiting its ability to access the glutamate binding site. We tested this with single molecule Förster resonance energy transfer (smFRET) measurements of detergent-solubilized, surface-immobilized SNAP-mGluR2 or SNAP_f_-mGluR2 dimers (**Fig. S3a-b)**^23^. In the presence or absence of 1 mM glutamate, histograms of FRET values measured from individual molecules showed sharper peaks for SNAP_f_-mGluR2 compared to SNAP-mGluR2 (**Fig. S3c-f**), suggesting that labelled SNAP_f_ is indeed more rigid than SNAP. Together these data point to flexibility of the labeling tag itself as a determinant of PORTL photoswitch efficiency. The flexibility of SNAP may also explain how BGAG_0_, a short ligand with a limited reach, is able to provide efficient photoactivation. Furthermore, this finding should also inform the application of SNAP tags for conformational studies or the development of biosensors.

### Branched BGAGs enable near-complete optical control of SNAP-mGluR2

Despite optimization of BGAG length, inter-domain linkers and SNAP variants, a photoswitch efficiency of ∼50-60% appeared to be an upper limit for the system (**Fig. 1b**). We sought a strategy to improve this efficiency. Following illumination, azobenzenes reach a photostationary state (PSS) with a certain proportion of molecules populating the high-energy *cis*-isomer. At optimal wavelengths, typically 80-90% of bis-acylated 4,4’-diaminoazobenzene scaffolds are able to populate the *cis*-state^24^. When factoring in mGluR2 cooperativity^15^, this 80-90% photoisomerization translates to ∼60-80% photoswitch efficiency, even with 100% labeling efficiency (**Fig. S4a**). Based on this, we reasoned that a molecule with two azobenzene groups and, thus, two independent chances of photoisomerization would increase the photoswitch efficiency. Since the azobenzene glutamate head group is the pharmacologically active part of BGAGs, we envisioned a branched molecule containing two azobenzene glutamates (**Fig. 1c**). Indeed, when we calculated the expected photoswitch efficiencies for hypothetical “doubleBGAG” molecules across a range of systems of different cooperativities we predicted clear improvements across all photostationary states (**Fig. S4a**) and over all labeling efficiencies (**Fig. S4b**). When we used our most likely scenario of mGluR2 cooperativity with a PSS of 85% and labeling efficiency of 90%, we estimated that “doubleBGAG” would substantially enhance photoswitching efficiency to >80% (**Fig. S4c, d**). It is worth noting that these calculations do not take into account any potential changes of the local, effective glutamate concentrations which, in principle, should double for such compounds.

The design of branched BGAGs is based on previously described BGAG compounds to maintain an efficient synthetic pipeline, and BG-COOH was chosen as the functionalizable bioconjugation motif for SNAP (**Scheme S1**). Next, the branching moiety and point had to be considered. Since the azobenzene is linked to the PEG chain by a glycine linker, we anticipated that an additional branching amino acid would not significantly prolong the attachment chain. Fmoc-glutamate thus appeared well-suited and was reacted as its doubly-activated NHS ester with two azobenzene glutamate precursors (**Scheme S2**). After Fmoc deprotection, the PEG_12_ chain was installed and finally conjugated to BG to obtain doubleBGAG_12_ (“dBGAG_12_”) (**Fig. 1d; Scheme S2**).

dBGAG_12_ showed similar photophysical properties compared to BGAG_12_ (**Fig. S5a, b**) and similarly efficient labeling of SNAP-mGluR2 (**Fig. S5c**). When conjugated to SNAP-mGluR2, dBGAG_12_ enabled optical control of mGluR2 with nearly complete efficiency relative to saturating glutamate (**Fig. 1e, f**). As predicted (**Fig. S4b**), even at decreased labeling efficiencies dBGAG_12_ enhanced photoswitch efficiency (**Fig. S6a, b**). Facile tuning of photocurrent amplitude was permitted by varying the illumination wavelength (**Fig. S6c**). In addition to enhancing overall efficiency, dBGAG_12_ also produced faster optical activation of mGluR2 (**Fig. 1g**) likely due to either a critical amount of active *cis*-conformers being reached more quickly or faster downstream signaling due to more complete population of the receptor’s active state.

To test if the enhanced efficiency of dBGAG_12_ would be maintained in a more complex, physiological environment with native effectors, we turned to cultures of cortical neurons. Following transfection of SNAP-mGluR2, photoactivation with either dBGAG_12_ or BGAG_12_ led to robust hyperpolarization and decreased neuronal firing, but the magnitude of the effect was larger for dBGAG_12_ (**Fig. 1h-j**). dBGAG_12_-mediated photo-activation was bistable (**Fig. S6d**) and substantially faster in neurons compared to HEK 293T cells (τ_on_ = 0.30 ± 0.10 s and τ_off_ = 0.48 ± 0.07 s in neurons, n= 12 cells). Confirming that the rapid light-induced hyperpolarization was indeed mediated by mGluR2 activation, the group II mGluR antagonist LY341495 was able to fully abolish light-induced effects on membrane potential (**Fig. S6e**). Together these experiments show that dBGAG_12_-mediated optical control of mGluR2 is well-suited to studies in the native neuronal context.

Given the utility of branched BGAGs, we decided to further characterize this new approach. We first asked if the presence of two azobenzene-glutamate moieties would lead to increased basal activation of mGluR2 by *trans*-dBGAG_12_ in the dark. Application of a saturating concentration of LY341495 had a marginal effect on baseline current which was identical for unlabeled, BGAG_12_- or dBGAG_12_-labeled SNAP-mGluR2, indicating that there is minimal basal receptor activation in all systems (**Fig. S7a-b**). We also photoisomerized dBGAG_12_ in the presence of glutamate and, as was reported for BGAG_12_, did not observe any current decrease (**Fig. S7c**), consistent with dBGAG_12_ serving as a full rather than a partial agonist. To probe the effective concentration of the glutamate groups of dBGAG_12_, we analyzed photoswitching in SNAP-mGluR2-R57A, a low affinity variant that we previously characterized with BGAG_0_ and BGAG_12_^10^. Whereas single chain BGAG variants showed no photoactivation in this mutant, small but detectable photoactivation was observed with dBGAG_12_ (**Fig. S7d, e**) confirming that the local concentration is increased with this PORTL. Based on the concentration-dependence of this mutant, we estimate that dBGAG_12_ mimics a local glutamate concentration of ∼100 μM. In addition to weak photoactivation in the absence of glutamate, dBGAG_12_ also showed large photocurrents in the presence of sub-saturating 1 mM glutamate (**Fig. S7d, e**), consistent with previous studies showing cooperativity where agonizing two subunits produces substantially larger activation than agonizing one^15^. These experiments show that dBGAG_12_ only weakly increases the local concentration of the tethered azobenzene-glutamate on mGluR2, supporting PSS-based enhancements (**Fig. S4**) as the main contributor to the improved photoswitching.

We next explored the specific composition of the branched PORTL by synthesizing the following branched BGAG variants: a short variant dBGAG_0_ (**Scheme S3**), dBGAG_12,v2_ (**Scheme S4**) where the branch point is placed just after the BG and, thus, two 12-repeat PEG chains are included in the compound, and qBGAG_12_ (**Scheme S5**) where four azobenzene-glutamates are added in parallel. dBGAG_12,v2_ permitted high-efficiency photoactivation of SNAP-mGluR2 to the same level as dBGAG_12_ (**Fig. S8a**), indicating that the branch location is not a critical parameter. Similarly, qBGAG_12_ also showed high efficiency photoactivation but did not allow any further enhancement (**Fig. S8b**), likely due to the fact that the proportion of subunits with a *cis*-azobenzene is already at saturation in dBGAGs (**Fig. S8c**). In contrast, and despite the robust photoswitching seen with BGAG_0_, dBGAG_0_ provided photoactivation of SNAP-mGluR2 with only ∼20% efficiency (**Fig. S8d**). This result suggests that branching may restrict the conformational freedom of BGAGs and, in the case of the short dBGAG_0_, this entropic cost reduces the efficiency of ligand binding.

Together these data provide a strategy for enhancing optical control of SNAP-mGluR2. To see if this branching-based strategy would also improve a less optimal PORTL system, we tested dBGAG_12_ on SNAP_f_-mGluR2 and observed a clear improvement in photoswitch efficiency from ∼30% to ∼50% (**Fig. S8e, f**). Given the robust improvement in photoswitch efficiency afforded by branched BGAGs on SNAP-based PORTLs, we next wondered if the strategy of branching could be generalized to other mGluR2-targeting PORTLs.

### Generalization of branched PORTLs to other labeling modes and spectral variants

A major advantage of the PORTL system is the ability to design and synthesize new PORTLs by flexible mix-and-matching of different chemical moieties (**Fig. 2a**). We hoped that this modularity would enable a toolset of photoswitchable mGluR2 variants optimized for different applications based on the desired attachment chemistry or spectral properties. To enable high efficiency optical control of mGluR2 tagged with CLIP, a variant with orthogonal labeling to SNAP^25^, we synthesized “doubleBCAG_12_” (dBCAG_12_) (**Scheme S6**). Similar to dBGAG_12_, dBCAG_12_ enabled near-complete optical control of CLIP-mGluR2 (**Fig. 2b-d**). Branching did not alter the specificity of dBCAG_12_, which showed no photocurrent when applied to cells expressing SNAP-mGluR2 (2.8 ± 0.3% relative to 1 mM glutamate, n=3 cells; labeled at 1 µM).

**Figure 2.**
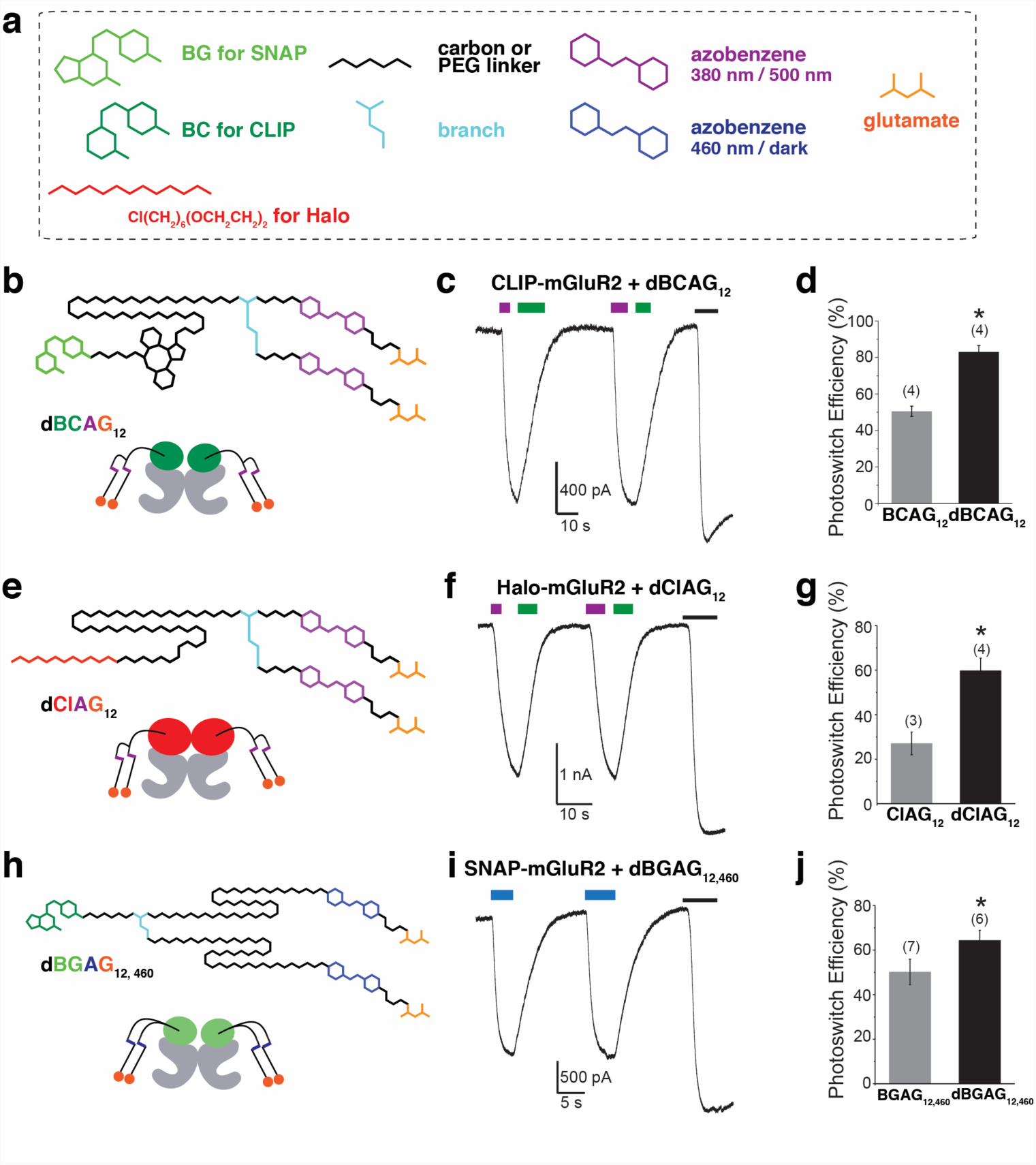
PORTL branching enhances mGluR2 photoactivation in a range of modalities. (**a**) Toolset of chemical moieties for mix-and-match design of PORTLs for SNAP, CLIP and Halo-tagged glutamate receptors. (**b-d**) doubleBCAG_12_ enhances efficiency of CLIP-mGluR2 photoactivation. * indicates statistical significance (unpaired t-test, p=0.04). (**e**-**g**) Photoactivation of Halo-mGluR2 by “ClAGs”. Branched doubleClAG_12_ (e) shows enhanced efficiency compared to ClAG_12_ (f-g). * indicates statistical significance (unpaired t-test, p=0.02). (**h-j**) doubleBGAG_12,460_ enhances efficiency of visible light-mediated (blue bar=460 nm) photoactivation of SNAP-mGluR2 relative to BGAG_12,460_. * indicates statistical significance (unpaired t-test; p=0.05). The numbers of cells tested are shown in parentheses. Error bars show s.e.m.

With the goal of further expanding the repertoire of PORTLs to a third self-labelling, orthogonal suicide enzyme, we aimed to implement the Halo-tag. The Halo-tag is an evolved dehalogenase that reacts specifically with alkyl chlorides to form a covalent bond^26^. Compared to BG and BC substrates, alkyl chlorides offer better permeability, a smaller and more physiological side product in chloride and higher acid stability, the latter which simplifies synthetic protocols. As such, we first synthesized the terminal alkyl chloride bearing ClAG_12_ (**Scheme S1, S7**) and cloned an N-terminally Halo-tagged mGluR2 construct. Based on the hypothesis that branching Halo-targeting PORTLs would enhance photoswitching, we also synthesized “doubleClAG_12_” (**Scheme S8; Fig. 2e**). Both PORTLs showed photoactivation of mGluR2 with identical spectral properties to BGAGs and BCAGs (**Fig. 2f**), but the efficiency of photoactivation of Halo-mGluR2 was boosted by branching with dClAG_12_ (**Fig. 2f, g**). This result introduces the Halo tag to the branched PORTL approach and expands the toolkit to three distinct, orthogonal protein tags.

A key advantage of azobenzene-based photoswitches is the ability to tune the photochemical properties of the compound. The previously reported red-shifted BGAG_12,460_ allows for visible-light induced, fast-relaxing photoactivation of mGluR2 which is advantageous in some settings, including for vision restoration applications^16^. To test if branching would also enhance PORTLs with a distinct azobenzene core, we synthesized “doubleBGAG_12,460_” and observed enhanced visible light photoactivation of SNAP-mGluR2 (**Scheme S6; Fig. 2h-j**).

Finally, a long-term goal of tethered photopharmacology is to incorporate the optical control afforded by such compounds into antibody-mediated targeting of proteins. Along these lines, we recently reported a new entity termed nanobody-photoswitch conjugates (NPCs) consisting of a SNAP-tagged nanobody labeled with a PORTL. NPCs containing an anti-GFP nanobody are able to photoactivate GFP-tagged mGluR2, albeit with limited efficiency^27^. We wondered if branched BGAGs would enhance the modest photoswitch efficiency observed with this system. Similar to all other systems tested, dBGAG_12_ doubled the photoswitch efficiency of NPC-mediated photoswitching of mGluR2 (**Fig. S9**). This result further confirms that branched PORTLs are an effective general strategy for improving photoswitch efficiency that should facilitate the further development and application of strategies for receptor targeting and optical control.

### Branched PORTLs drastically enhance optical control of mGluR3

Throughout the nervous system a variety of receptors work in parallel to control glutamatergic signaling^13^. Given this molecular diversity, it is desirable to obtain the optical control of multiple mGluRs with high efficiency for comparative or multiplexed studies. Toward this goal, we turned to mGluR3, the other member of the group II mGluR subfamily. Despite the challenges in deciphering the unique properties of mGluR2 and 3 due to the paucity of subtype-specific drugs, recent work has found distinct activation dynamics^23^ and subcellular localization^28^ between receptor subtypes and a high propensity for heterodimerization between mGluR2 and 3^15^, raising the motivation for PORTLs to dissect the contributions of each group II mGluR subtype to biological processes. Previous work with BGAG PORTLs has shown only weak ∼20% photoactivation of SNAP-mGluR3 with unbranched BGAGs^10^, precluding useful application of this tool. Unlike SNAP-mGluR2, SNAP-mGluR3 photoactivation shows steep length-dependence with the short variant, BGAG_0_, providing the highest efficiency. This suggests that mGluR3 photoactivation is limited, in part, by the local concentration of the azobenzene-glutamate moiety. We further characterized SNAP-mGluR3 photoswitching with unbranched BGAGs to see if SNAP-mGluR3 photoswitch efficiency could be enhanced by modifying the receptor construct. Neither introduction of SNAP_f_ to produce SNAP_f_-mGluR3 (**Fig. S10a, b**) nor addition of a flexible (GGS)_2_ linker between SNAP and mGluR3 enhanced photoswitching with BGAG_0_ or BGAG_12_ (**Fig. S10b**).

We next turned to branched BGAGs in the hope that the increased valence of the system along with the increase in the proportion of labeled subunits with a *cis*-azobenzene (**Fig. S4**) would enhance the enable efficient photoswitching. Indeed, dBGAG_12_ drastically improved photoactivation of SNAP-mGluR3 to levels comparable to SNAP-mGluR2 (**Fig. 3a-c**). To our surprise, neither dBGAG_0_ or dBGAG_12,v2_ were able to enhance photoswitch efficiency (**Fig. 3c**). It is difficult to account for the dramatically enhanced efficiency of branched PORTLs with either the increased proportion of subunits containing a *cis*-azobenzene or the modest, two-fold increase in local glutamate concentration. Furthermore, the dependence on specific branching pattern suggests that the branch point introduces secondary interactions with the mGluR3 LBD that provide some binding enthalpy to drive photo-agonism. Reducing the glutamate affinity of SNAP-mGluR3 by introducing an alanine at arginine 64, the homologous position to R57 in mGluR2, decreased but did not abolish dBGAG_12_ photoswitch efficiency (**Fig. S10c, d**). This result suggests that, similar to the case with mGluR2, the glutamate moiety is in an effective concentration range of hundreds of micromolar. We also tested dBCAG_12_ on CLIP-mGluR3 and observed drastically enhanced photoswitch efficiency compared to BCAG_12_ (**Fig. 3d-f**). Ultimately, the ability to optically control either mGluR2 or mGluR3 with either CLIP or SNAP-tags opens the door to dissecting their distinct functions in the nervous system.

**Figure 3.**
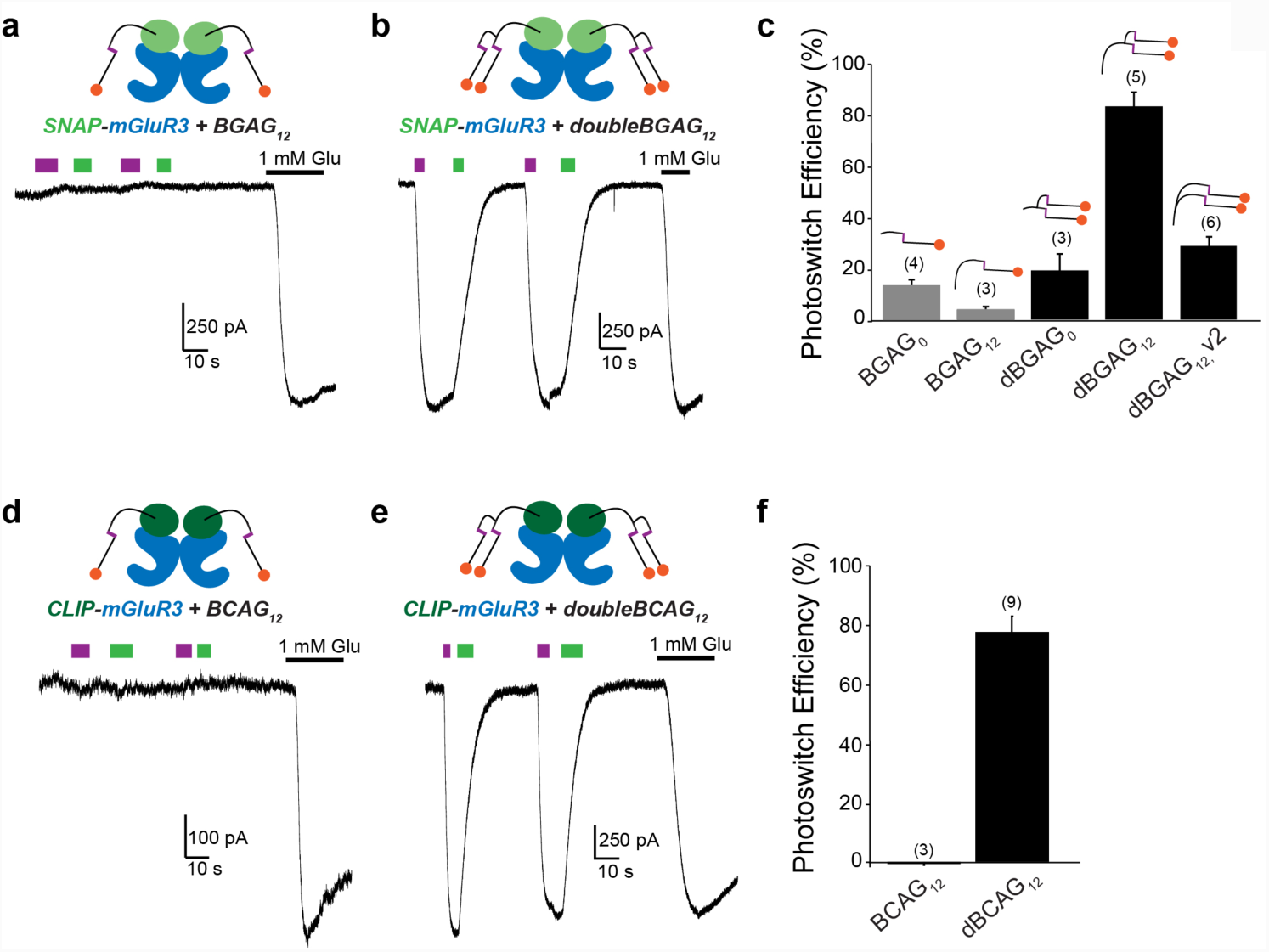
Branched PORTLs enable efficient optical control of mGluR3. (**a**-**b**) Representative traces showing photoactivation of SNAP-mGluR3 with BGAG_12_ (a) and dBGAG_12_ (b). (**c**) Summary bar graph showing optimal photoswitching of SNAP-mGluR3 with dBGAG_12_, but not dBGAG_12,v2_ or dBGAG_0_. (**d-e**) Representative traces showing photoactivation of CLIP-mGluR3 with BCAG_12_ (d) and dBCAG_12_ (e). (**f**) Summary bar graph showing optimal photoswitching of CLIP-mGluR3 with dBCAG_12_. The numbers of cells tested are shown in parentheses. Error bars show s.e.m.

### Efficient optical control of mGluR5 signaling with spatiotemporal precision

We next turned to mGluR5, a group I mGluR that is highly expressed throughout the central and peripheral nervous systems. A limited and debated toolset of specific, orthosteric compounds for group I mGluRs exists^29^ challenging the dissection of mGluR1 versus mGluR5 biology. Furthermore, mGluR5 is a prime target for PORTL-mediated control for a number of reasons. mGluR5 is expressed in a wide range of cell types within the brain, including excitatory principal cells, inhibitory interneurons and astrocytes, making it difficult to dissect the role of specific mGluR5 populations with global drug application or knock-out. While targeted knockouts have provided some insight^30,31^, the ability to photoactivate mGluR5 in defined cell types would allow one to approach such questions with high precision and reversibility within neural circuits. Furthermore, mGluR5 has myriad signaling partners, interacts with an extensive network of scaffold proteins via its large C-terminal domain (CTD) and couples to a variety of effectors and regulators ^13^. PORTL-based control of mGluR5 would allow for the testing of receptor mutants in native systems to probe the role of specific interactions or regulatory sites. For example, mGluR5 activation of G_q_ is known to induce a unique form of calcium oscillations that are thought to be due to reversible protein kinase C (PKC)-mediated phosphorylation of a residue on the membrane-proximal part of the CTD^32,33^. Finally, a deeper understanding of mGluR5 neurobiology is motivated by the roles of mGluR5 in long term synaptic depression and as a promising drug target for neurodevelopmental and neuropsychiatric disorders^34,35^.

To date, group I mGluRs have not been successfully photosensitized with PTLs or PORTLs, likely due to pharmacological incompatibility of the azobenzene-glutamate moiety. However, we and others have shown that extracellular domains of mGluRs are portable and can effectively gate the TMDs of other mGluR subtypes^15^. With this in mind, we reasoned that a chimera between mGluR2 and mGluR5 that contains the entire TMD and CTD of mGluR5 would maintain the unique properties of mGluR5 signaling, while allowing efficient PORTL-mediated optical control.

We thus engineered a SNAP-mGluR2-5 chimera (**Fig. 4a**) and observed similar glutamate-evoked oscillatory calcium responses to mGluR5 in HEK 293T cells (**Fig. S11a**). SNAP-mGluR2-5 showed strong surface expression and the expected glutamate dose-response (**Fig. S11a, b**). We next tested if conjugation with BGAGs would enable photoactivation of mGluR5 signaling and found that, indeed, subcellular 405 nm illumination produced BGAG-dependent, repeatable, robust calcium oscillations (**Fig. 4b; Fig S11c, d**) of a similar frequency to glutamate (1-3 oscillations/minute). In line with the enhanced efficiency of branched BGAGs on SNAP-mGluR2, light responses were more reliable, in terms of the percentage of cells that responded, for dBGAG_12_ versus BGAG_12_ (**Fig. 4c**). Light-induced calcium oscillations were blocked by MPEP, an mGluR5 negative allosteric modulator (**Fig. S11e**), or enhanced by VU 0360172, an mGluR5 positive allosteric modulator (**Fig. S11f**), showing that the mGluR5 TMD maintains its native allosteric binding site within this chimera.

**Figure 4.**
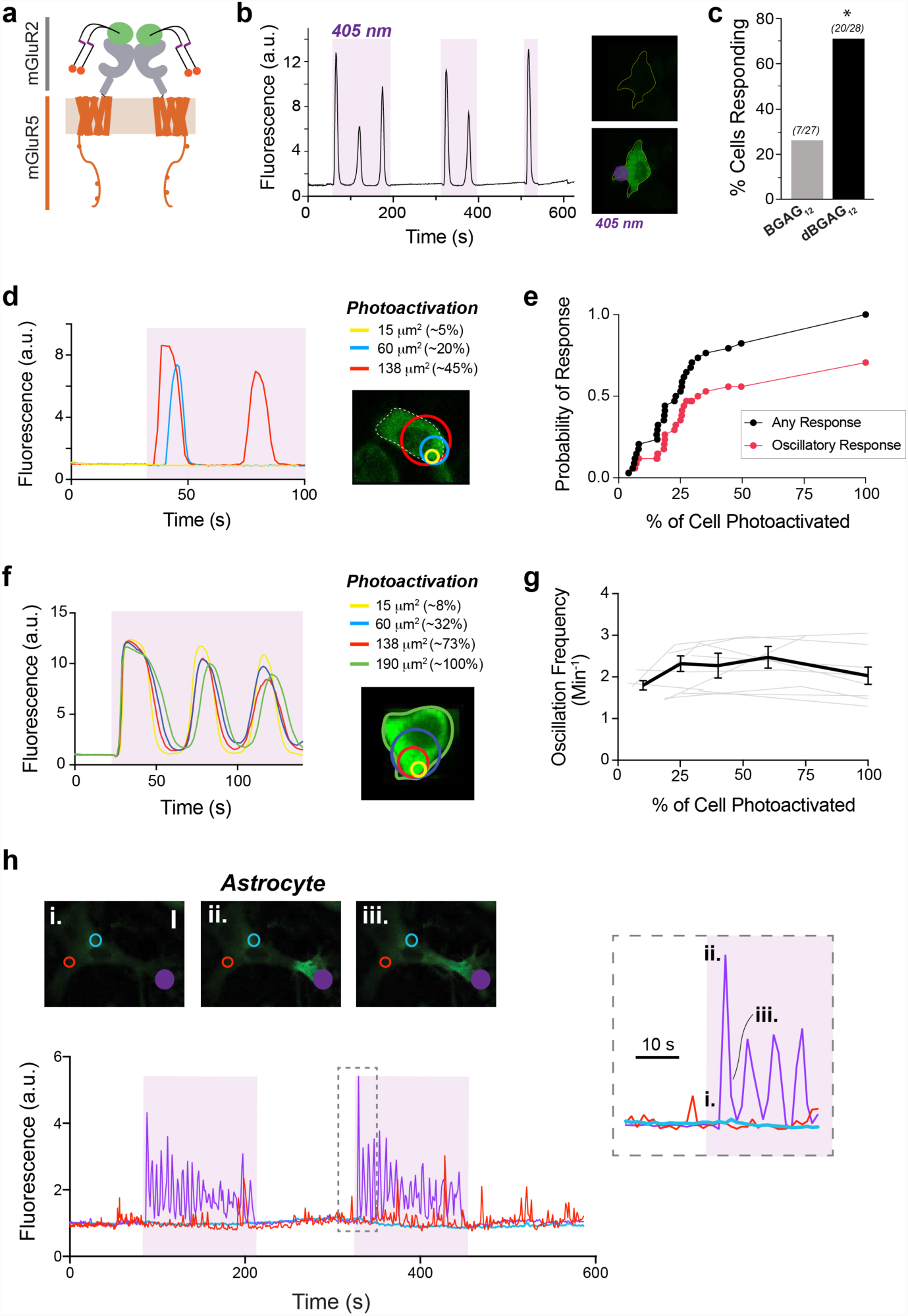
Optical control of mGluR5 signaling via branched PORTLs and a chimera-based approach. (**a**) Schematic showing design of chimera including an N-terminal SNAP tag, the extracellular domains of mGluR2 and the transmembrane and intracellular C-terminal domains of mGluR5. (**b**) GCaMP6 calcium imaging in HEK 293T cells reveals 405 nm-induced calcium oscillations mediated by SNAP-mGluR2-5 and dBGAG_12_. Right, corresponding images of cells before (top) and during (bottom) subcellular 405 nm illumination (purple dot). 488 nm imaging light is sufficient to rapidly de-activate dBGAG_12_ following 405 nm illumination. (**c**) Summary of efficiency of optical control of SNAP-mGluR2-5 for BGAG_12_ versus dBGAG_12_. * indicates statistical significance (Pearson’s chi-square test, p=0.002). (**d-e**) Increasing the subcellular area of photoactivation increases the probability of a single peak or oscillatory light response. Data from 38 cells were included in this analysis. (**f-g**) Increasing the subcellular area of photoactivation does not alter the frequency or amplitude of oscillatory light responses. The summary plot shows data for individual cells (grey lines; n=10 cells) and from an average of all tested cells (black). Error bars show s.e.m. (**h**) Subcellular photoactivation of SNAP-mGluR2-5 with dBGAG_12_ in cultured astrocytes produces reliable calcium oscillations and allows for the visualization of subcellular calcium waves. Photoactivation occurred only at the purple circle. Inset highlights that, in this representative cell, calcium oscillations occur in the soma (purple, green), but not in processes (red, blue).

An important application of PORTL-mediated control of mGluR5 is the ability to probe specific functional aspects of mGluR5 rather than to merely control generic G protein signaling cascades, as is typically done with DREADDs or opsin-based tools^36^. To further test if mGluR5 identity is maintained in our system, we probed the role of the mGluR5 PKC phosphorylation site, Ser839, in induction of light-induced calcium oscillations. Consistent with previous studies on mGluR5, mutation of Ser839 to alanine (S839A) nearly abolished the oscillatory response to both glutamate (**Fig. S12a, b**) and light (**Fig. S12c**) in our construct.

To test the power of spatiotemporal control for the quantitative study of mGluR5 signaling, we next used targeted photoactivation to small regions of the cell to probe the properties of evoked calcium oscillations. We first asked if the size of the illumination spot was a determinant of response probability and observed that, indeed, increasing the area of photoactivation increased the probability of observing a calcium response (**Fig. 4d, e**). **Fig. 4d** shows an example cell where 3 different-sized, overlapping photoactivation areas were targeted; the smallest ROI failed to produce an intracellular calcium increase, the intermediate size elicited a single peak and the largest ROI produced an oscillatory response. **Fig. 4e** summarizes the photoactivation size-dependence of calcium responses and shows that ∼25% of the cell was needed to have a 50% chance of a response. Consistent with previous work on mGluR5^37^, SNAP-mGluR2-5-induced calcium oscillation frequency was largely independent of the concentration of glutamate (**Fig. S13a, b**). Interestingly, in contrast to the weak effect of increasing glutamate concentration, it has been shown that increased receptor expression levels increase the frequency of calcium oscillations^37^. Given this, we asked if the area of photoactivation of a cell would lead to alterations in the calcium oscillation frequency. We varied the size of the photoactivation area and measured the subsequent calcium oscillation frequency and found that the frequency was largely insensitive to the proportion of the cell that was targeted (**Fig. 4f, g**). We also examined how calcium oscillations spread throughout the cell by comparing the calcium signal at the site of photoactivation to a distal part of the cell. Interestingly, both the temporal profile (**Fig. S13c**) and calcium response amplitude (100.6 ±13.5 % of the amplitude at the activation site, n= 6 cells) were maintained at the distal site. Furthermore, we were able to use the offset in signals to calculate a calcium wave velocity for each cell of 10-20 µm/s (**Fig. S13d**), which is comparable to prior measurements of spontaneous calcium waves in mammalian cells^38^.

One of the key mechanisms by which mGluR5 signaling is thought to modulate neural activity is through signaling within astrocytes, especially in the developing brain^39,40^. Astrocytic mGluR5 can respond to synaptic glutamate to control astrocyte calcium signaling dynamics, gliotransmission and morphology^41-43^, but the inability to isolate astrocytic mGluR5 signaling from neuronal mGluR5 signaling has hampered progress on the underlying mechanisms of this form of neuromodulation. As previously reported, cultured astrocytes from mouse hippocampus and cortex showed calcium elevations in responses to mGluR5 activation^44-46^, with a mix of single peaks and oscillations (**Fig. S14a-c**). We next asked if our photoswitchable mGluR5 system could allow us to mimic native mGluR5 responses and permit the use photoactivation to probe this system. Photoactivation of dBGAG_12_-labeled SNAP-mGluR2-5 in astrocytes initiated either single peaks or calcium oscillations (**Fig. 4h; Fig. S14d, e**) of higher frequency than in HEK cells (3-6 oscillations/minute). In the absence of the receptor construct, no light responses were observed (**Fig. S14f**). Targeted photoactivation to small areas revealed signal spreading heterogeneity between the soma and extended processes that was different from what was observed in the spatially-homogenous HEK 293T cells. When photoactivation was targeted to the soma, oscillations were observed in the soma in ∼55% of cells of those responding cells and oscillations were observed in at least one process in ∼50% of those cells (**Fig. S14g**). However, when photoactivation was targeted to a process, local oscillations were observed at the targeted process in ∼65% of cells but were only seen in the soma in 15% of those cells (**Fig. 4h; S14h**), consistent with previous observations of global or compartmentalized calcium responses in astrocytes^47^. In addition, oscillations were of a higher frequency in the processes compared to in the soma (6.9 ± 0.8 oscillations/minute in processes vs. 4.5 ± 0.7 oscillations/minute in soma; n= 4 cells where both sites were targeted). Together this work demonstrates the suitability of our approach for probing mGluR5 signaling dynamics in astrocytes and motivates future applications to dissect the contribution of neuronal versus astrocytic mGluR5 signaling to synaptic and circuit-level processes.

### Dual photo-activation and imaging with branched PORTL/fluorophore compounds

Optogenetic actuators, such as the PORTL-based control of mGluRs described here, are powerful tools for dissecting the molecular basis of biological processes. However, a complete understanding of the underlying molecular events also requires precise sensing of protein localization and/or conformation. Combining the ability to optically manipulate and sense the same receptor population would be a particularly powerful means of obtaining a full picture of receptor function. Chemical conjugation of expressed proteins with organic fluorophores produces brighter fluorescence compared to fluorescent proteins, allows one to maintain flexibility for using different spectral variants with the same protein target and permits surface versus intracellular targeting^48^. Combining attachment of fluorophores and photoswitch actuators to the same receptor population would enable the simultaneous study of receptor localization and/or conformational state while controlling its activity, a potentially very powerful technique.

Given our demonstration that SNAP-tagged receptor targets tolerate PORTL branching, we decided to test if incorporation of a fluorophore into a branched BGAG would enable dual optical manipulation and detection. As such, we chose the same branching point endowed with a fluorophore instead of a second azobenzene-glutamate. The fluorophore should be spectrally orthogonal to switching wavelengths, rendering Cy5 with its absorption/emission wavelengths in the far-red (650/670 nm) perfectly suited. Accordingly, Cy5 was installed on the ε-amine of lysine, and whose acid was coupled to the azobenzene-glutamate before linking to BG and a PEG_12_ chain to obtain “BGAG_12_-Cy5” (**Fig. 5a; Scheme S8**). BGAG_12_-Cy5 showed an expected absorption spectrum with peaks for both the azobenzene and Cy5 (**Fig. S15a**), and showed efficient labeling of SNAP-mGluR2 over the same concentration range as other BGAGs (**Fig. S15b**). Most importantly, BGAG_12_-Cy5 permits efficient optical control of SNAP-mGluR2, while also allowing imaging of the receptor on the plasma membrane of cells (**Fig. 5b; Fig. S15b**). The photoswitch efficiency and kinetics were indistinguishable from BGAG_12_ alone (**Fig. S15c**), indicating that the incorporation of Cy5 did not alter the pharmacology of the azobenzene-glutamate.

**Figure 5.**
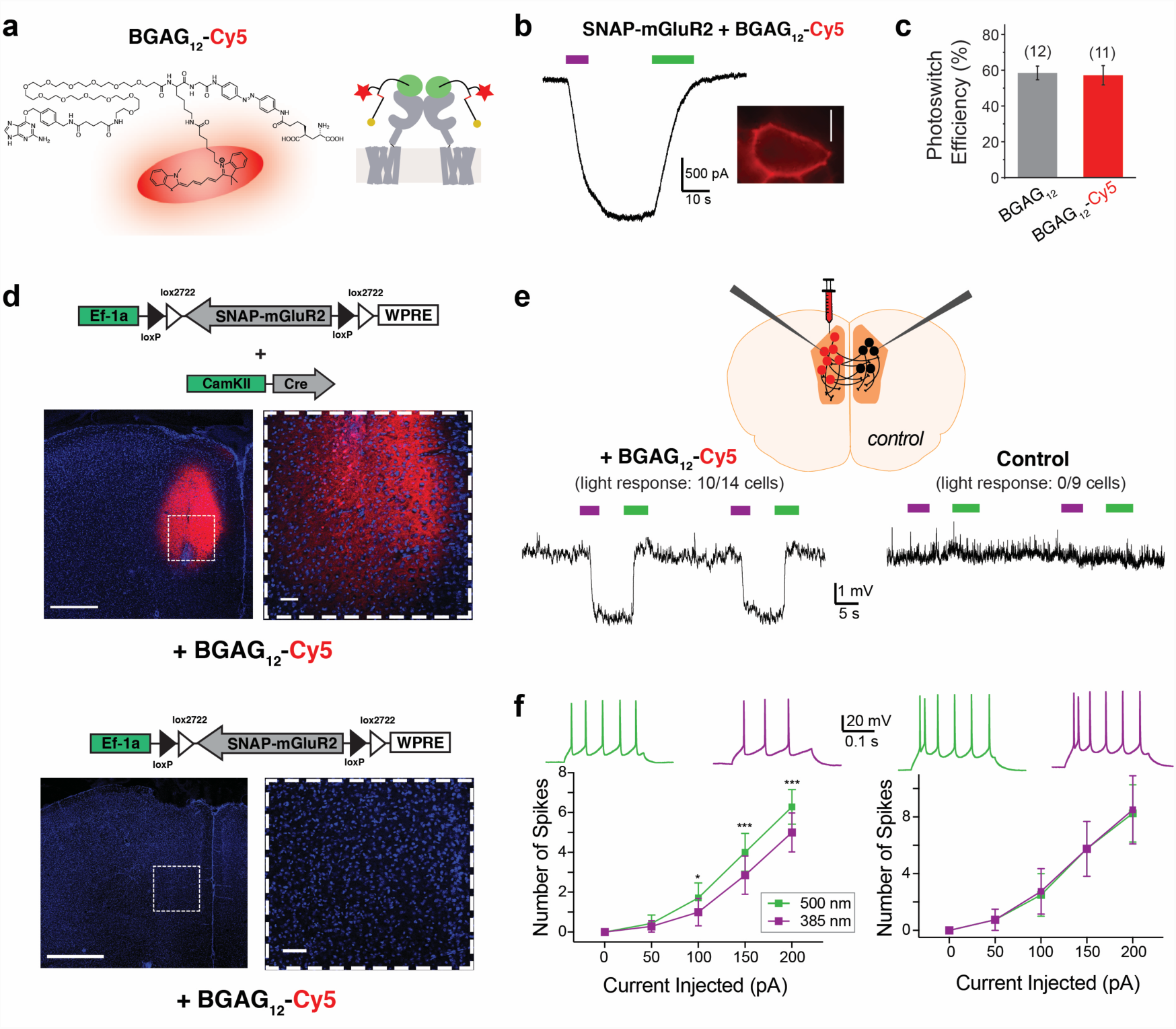
Branched fluorophore-containing BGAGs allow dual photoactivation and detection of mGluR2 in cultured cells and acute slices following *in vivo* labeling. (**a**) Chemical structure, left, and schematic, right, showing BGAG_12_-Cy5, a PORTL for dual optical manipulation and sensing of mGluR2. (**b**) Representative trace, left, and image, right, showing photoactivation and detection of SNAP-mGluR2 in a HEK 293T cell. Scale bars= 10 μm. (**c**) Summary bar graph showing comparable photoswitch efficiency relative to saturating glutamate for BGAG_12_-Cy5 and BGAG_12_. (**d**) Top, images showing SNAP-mGluR2 labeled with BGAG_12_-Cy5 following viral expression and *in vivo* PORTL injections. Bottom, images showing control slices from mice injected with BGAG_12_-Cy5 but not expressing SNAP-mGluR2. Scale bars= 500 μ (left) or 50 (right) μm. (**e**) Top, schematic showing experiment where viral delivery and BGAG12-Cy5 injection is only done in one hemisphere of the medial prefrontal cortex of a mouse. Bottom, representative current clamp traces showing light-induced hyperpolarization only in the fluorescent hemisphere and not in the non-fluorescent “control” hemisphere. (**f**) Input-output curves showing current induced firing for cells in the fluorescent experimental hemisphere (left) and the non-fluorescent control hemisphere (right). * indicate statistical significance (paired t-test, p= 0.05 for 100 pA injection, p=0.003 for 150 pA injection, p=0.003 for 200 pA injection). Inset shows representative spike firing traces following 525 nm (green) or 385 nm (purple) illumination. Error bars show s.e.m.

Labeling with BGAG_12_-Cy5 should provide a real time view of which cells or tissue regions have labeled, *photoswitchable* receptors. This would both enhance the efficiency of identifying expressing, labeled cells as well as defining precise cellular or subcellular targeting experiments. To test if BGAG_12_- Cy5 can be used for facile identification of the region of expression and labeling, we injected viruses (Cre-dependent FLEX-SNAP-mGluR2 and Cre recombinase under the CaMKII promoter) and BGAG_12_- Cy5 into the medial prefrontal cortex (mPFC) of mice in only one hemisphere. The mPFC was targeted because this is an area of native mGluR2 expression^49^ and because of its involvement with many of the psychiatric disorders for which mGluRs are implicated. Confocal images demonstrate clear targeting of SNAP-mGluR2 by BGAG_12_-Cy5 with no background labeling in mice infected with FLEX-SNAP-mGluR2 but not the CaMKII: Cre virus (**Fig. 5d**). This demonstrates the suitability of branched PORTLs for *in viv*o labeling of SNAP-tagged receptors without background. We then performed patch clamp recordings from slices prepared from injected mice. The fluorescence from BGAG_12_-Cy5 allowed for the easy identification of the site of receptor expression and labeling. We recorded from fluorescent cells in the injected hemisphere and non-fluorescent cells in the control hemisphere of the medial PFC and observed a light-induced hyperpolarization only in the fluorescent hemisphere (**Fig. 5d**). In addition, photoactivation of mGluR2 produced a reversible, repeatable decrease in spike firing in response to current injection (**Fig. 5e**). The modest 2-5 mV hyperpolarization and decreased excitability observed was similar to what is typically seen for endogenous or heterologously-expressed G_i/o_-coupled receptors in cortical pyramidal cells^50,51^. Importantly, no light responses were observed in slices from wild type mice injected with BGAG_12_-Cy5 (**Fig. S15d**) and expression and labeling of SNAP-mGluR2 did not alter the resting membrane potential of neurons (**Fig. S15e**). Notably, previous drug application-based slice studies of GPCRs have provided minimal information on the kinetics and repeatability of receptor effects, likely due to long bath exchange and tissue penetration times on the seconds to minutes time scales. Here, photo-activation of SNAP-mGluR2 revealed rapid on and off kinetics on the hundreds of milliseconds time scale (τ_ON_ = 330 ± 41 ms, τ_OFF_= 687 ± 64 ms; n= 13 cells) and clear reproducibility over many cycles (**Fig. S15f**), showing the utility of photoswitchable ligands for dissecting the precise timing of neuromodulatory processes in intact tissue. Together this demonstrates the utility of the bi-functional fluorophore/PORTL approach and provides a framework for using branching to further expand the world of PORTLs and PORTL-based applications.

## Discussion

### Branched PORTLs: A new strategy for high efficiency optical control and dual control and detection

Despite their emergence as useful tools for studying membrane receptor signaling, the applicability and generalizability of tethered photoswitches has remained limited by both a lack of understanding of the underlying mechanisms and a limited number of strategies for enhancing optical control of a given target. Here we report a family of branched, tethered photoswitchable ligands that enhance the efficiency of existing photoswitchable receptors (i.e. mGluR2) and enable extension of the approach to other, related targets (i.e. mGluR3, mGluR5). We show that a primary effect of branching is to increase the number of azobenzene moieties that reach the active *cis*-state. This effect overcomes the limitations of the incomplete *cis* occupancy achieved at all photostationary states and should be useful for a wide range of probes, including soluble photochromic ligands^2^. Furthermore, this re-design highlights the fact that cooperative, multimeric receptors which show non-linear ligand-occupancy dependent activation pose major obstacles to achieving high efficiency optical control. As a secondary effect, branching enhances the local concentration of the functional group which likely also contributes to the enhanced efficiency of the pharmacophore. This effect indicates that branching should be useful for all tethered ligand approaches, even those that do not incorporate photosensitivity^52-54^.

As a complement to our implementation of branched PORTLs, we also provide new insight into the mechanisms of tethered photoswitches. Along with previous studies^9,10^, we have found that in the case of SNAP-mGluR2 there is a tolerance for a wide range of PORTL lengths and lengths of the linker between the SNAP-tag and the receptor. In contrast, we find that tag flexibility itself is a key parameter needed when assessing self-labeling protein tags and their ability to be employed for PORTLs. Together these observations indicate that the efficiency of photoswitching is determined by the relative energetic contributions of the enthalpy change associated with binding of the active form of the functional group and the associated entropy loss of the protein tag and PORTL linker, similar to what has previously been described for a tethered enzyme inhibitor^55^. Further work focused on the pharmacology of the tethered ligand itself is needed to understand why different subtypes photoswitch to different degrees or with different directionality and should provide a further means of optimization.

The PORTL approach incorporates the spatiotemporal precision of light, genetic targeting, and receptor subtype-specificity. Furthermore, the use of full length receptors gives the ability to incorporate mutations to alter activation mechanism, effector coupling or regulation. Here we report high efficiency photoactivation of mGluR2 via branched PORTLs which target SNAP, CLIP or Halo tags and have different spectral properties. We also report that dBGAG_12_ and dBCAG_12_ allow near-complete photoactivation of SNAP-mGluR3 and CLIP-mGluR3, respectively. This demonstrates the generalizability of the branched PORTL approach and provides a toolset that allows the user to tailor the tool to their particular application. Indeed, the advantage of orthogonal labeling and distinct activation wavelengths of this toolset should enable multiplexing optical control of receptor populations. We previously demonstrated the ability to sequentially photoactivate SNAP and CLIP-tagged receptors with normal azobenzene (i.e. BGAG_12_) and red-shifted azobenzene cores (i.e. BGAG_12,460_)^10^. Based on this, one can design applications where different receptor populations in distinct cell types or subcellular regions are activated simultaneously or sequentially to probe their functional interactions. The greatly expanded toolset reported here should enable myriad studies of this type. Finally, we show that the branched PORTL approach also enhances the efficiency of photoactivation when tethered to nanobodies. The ability to enhance the initially modest optical control reported with this configuration^27^ should spur the development of this approach which has the potential to allow gene-free optical control of native receptors.

We also introduce branched PORTLs that incorporate both a photoswitchable ligand (azobenzene-glutamate) and an organic fluorophore (Cy5) to enable dual manipulation and detection of the same receptor population. Both PORTLs and dye conjugation-based detection techniques^48^ are useful on their own but incomplete labeling efficiency limits the ability to use dual conjugation of chemical ligands and sensors together if they target distinct tags on the same protein. Furthermore, non-fluorescent PORTLs are hard to localize in complex tissue where identifying expression sites is non-trivial. Here we show that BGAG_12_-Cy5 enables simultaneous imaging and activation of SNAP-mGluR2. In acute cortical slices, this approach permits easy identification of fluorescently-labeled cells which showed consistent light-induced neuronal inhibition. The use of PORTLs allowed us to measure activation and de-activation kinetics in the 100-700 ms time scale in this system, something that has not been accessible in pharmacological studies of GPCRs in brain slice. Overall, this demonstrates the design possibilities afforded by PORTL branching and should open the door to optical studies that link receptor activation to localization, mobility, and conformation with high temporal precision.

### Optical dissection of mGluR5-induced calcium signaling dynamics

We also report optical control of mGluR5 using a chimera-based design where the SNAP-tagged extracellular domain of mGluR2 is combined with the transmembrane domain and intracellular C-terminal domain of mGluR5. When conjugated to branched PORTLs this construct enables efficient photoactivation while maintaining the entire TMD and CTD of mGluR5, which are the main determinants of receptor localization, effector coupling, regulation and allosteric pharmacology. In contrast, other chimera-based optogenetic approaches^36,56^ employ the 7-TM core of rhodopsin along with the intracellular loops and CTDs of a given receptor which is unlikely to fully recapitulate the complex conformational and signaling dynamics of the target receptor. With this in mind, rhodopsin-based chimeras are ill-suited to class C GPCRs which contain large extracellular domains and constitutively dimerize^15^.

We used the spatiotemporal precision of this system to probe the nature of mGluR5-induced calcium oscillations. Unlike typical GPCR-induced calcium oscillations which are thought to be largely driven by cycles of IP_3_ receptor desensitization and re-sensitization^57^, mGluR5-induced calcium oscillations are dependent on reversible phosphorylation of a PKC site (S839)^32,33^, which is maintained in our construct. Here we find that neither glutamate concentration, as previously reported^37^, nor the proportion of the cell that is photoactivated alters the frequency of calcium oscillations. This suggests that beyond a response threshold, which we calculate as either ∼1 µM glutamate or activation of ∼25% of the cell, calcium oscillations occur at an intrinsic frequency for any given cell. This essentially turns mGluR5 into a binary switch rather than a dial, as receptor responses are normally modeled. What determines this intrinsic frequency? One possibility is that it is merely determined by the relative densities of receptors, G proteins, protein kinase C, phosphatases and other factors that shape G protein turnover and calcium handling. Altering the proportion of the cell that is activated or the glutamate concentration does not effectively change these ratios, only altering the density of each component does. This is consistent with a previous report that increasing the expression of receptors can increase the frequency of calcium oscillations^37^. Future work is needed to understand the mechanisms that determine calcium oscillation properties and how these oscillations are decoded by downstream process, such as transcription.

We also demonstrate mGluR5 photoactivation in astrocytes, where we find similar oscillatory responses following receptor activation, but clear evidence that different sub-domains of the cell appear to confine calcium oscillations. Most strikingly, we find that activation of mGluR5 in astrocytic processes often produced local calcium oscillations that did not travel to the soma. This suggests that synaptic glutamate may induce local calcium responses that do not spread globally throughout the cell, perhaps to maintain input specificity. Consistent with this notion, imaging of spontaneous signals in astrocytes has led to numerous examples of calcium waves which remain confined either within processes or smaller microdomains^47^. In the case of mGluR5 activation, determining the mechanism of signal confinement will require the identification and study of the relevant effectors, calcium sources and scaffolding proteins. Notably, Arizono et al^58^ found that astrocytes mGluR5 translocates more rapidly in processes than in the soma and that a diffusion barrier prevents exchange of receptors from the processes back to the soma. Together, these experiments demonstrate the power of PORTL-based optical control for a quantitative dissection of receptor-induced cell signaling dynamics with a precision not afforded by soluble ligands.

## Methods

### Chemical Synthesis

See Supporting Information for details on chemical synthesis and characterization.

### Single Molecule FRET

Single-molecule Förster resonance energy transfer (smFRET) experiments were performed on an inverted microscope (Olympus IX73) in total internal reflection (TIR) mode using a 100x objective (NA=1.49) and a 561 nm laser diode. Movies were recorded simultaneously with two sCMOS ORCA-Flash4 v3.0 cameras (Hamamatsu) separated by a 635 nm long-pass dichroic mirror and with appropriate emission filters for donor (595/50) and acceptor (655LP). A microflow chamber was assembled using a passivated glass coverslip prepared with mPEG-SVA and biotinylated PEG (MW=5000, 50:1 molar ratio, Laysan Bio) as previously described^23^. The microflow channels were first incubated with 0.2 mg/ml of NeutrAvidin (ThermoFisher) for 2 min, followed by 10 nM of biotinylated secondary donkey anti-rabbit, antibody (ThermoFisher, A16039) for 30 min, and 10 nM of anti-Metabotropic Glutamate Receptor 2+3 antibody (abcam, ab6438) for 30 min. Microchannels were washed 5 times between each conjugation step with T50 buffer (10 mM Tris, 50 mM NaCl, pH 8).

Following 48 hr expression of either HA-SNAP-mGluR2 or HA-SNAP_f_-mGluR2, HEK293T cells were labeled for 1 hour at 37 °C using a 1:3 molar ratio of BG-LD555 (1 µM) and BG-LD655 (3 µM) dyes (Lumidyne Technologies) dissolved in EX buffer containing (in mM): 10 HEPES, 135 NaCl, 5.4 KCl, 2 CaCl_2_, 1 MgCl_2_. After labeling, cells were washed 2 times with EX buffer, harvested by incubating the cells in PBS (Ca^2+^ and Mg^2+^ free) for 20 min, and lysed for 1 hr at 4 °C using lysis buffer containing (in mM): 150 NaCl, 1 EDTA, protease inhibitor cocktail (Thermo Scientific) and 1.2% IGEPAL detergent (Sigma). Finally, cell lysates were spun down at 16,000 × g for 20 min at 4 °C, supernatant was collected, and maintained on ice prior to imaging. Protein samples were diluted using a dilution buffer (EX containing 0.1% IGEPAL) and added to the microflow chamber until the density reached ∼400 molecules per 2,000 µm^2^. Any unbound proteins were washed using the dilution buffer. Single molecule fluorescence movies were recorded at one frame per 100 ms in the presence of an oxygen scavenging system (1 mg/ml glucose oxidase, 0.04 mg/ml catalase, 0.8% w/v D-glucose) and photostabilizing agents (5 mM cyclooctatetraene). smFRET data analysis was performed using SPARTAN^59^. FRET histograms (averaged from 3 separate movies per condition) were then plotted and fitted using a single Gaussian function to get a peak position and FWHM.

### Molecular Cloning, Cell Culture, and Gene Expression

Experiments were performed using previously described N-terminally SNAP-or CLIP-tagged rat or human mGluRs^60^. Mutations, insertions, and deletions were introduced using standard PCR-based techniques. SNAPfast or Halo were introduced into mGluR2 at the same position as SNAP using Gibson assembly. The chimeric SNAP-mGluR2-mGluR5 was made, also via Gibson assembly, using the extracellular domain of SNAP-tagged mGluR2 (K23 to E559) and rat mGluR5 transmembrane domain and C-terminal domain (Y571 to STOP). For adeno-associated virus production, SNAP-mGluR2 was cloned into a previously described FLEX construct (addgene ID: 26966).

HEK 293T cells (authenticated by Bio-Synthesis, Inc. and tested negative for mycoplasma using a kit from Molecular Probes) were plated at low density on poly-L-lysine coated coverslips (18 mm) and transfected the following day using either Lipofectamine 2000 or 3000 (Thermo Fisher). 24-48 hrs after transfection, cells were used for electrophysiology or imaging experiments. As previously described^10^, for electrophysiology experiments each coverslip received 0.7 µg each of the receptor of interest and GIRK1-F137S and 0.1 ug of tdTomato as a transfection marker. The GIRK1-F137S homo-tetramerization mutant ^61^ was used to reduce the number of constructs to simplify transfection and expression. For calcium imaging experiments, each coverslip received 0.7 µg of the receptor of interest plus 0.3 µg of GCaMP6f.

Cortical neurons were isolated from P1-2 mice and plated at 50,000-75,000 cells per coverslip on poly-ornithine-coated coverslips (12 mm). Neurons were plated in media containing DMEM supplemented with 5 g/L glucose, 100 mg/L Transferrin (Millipore), 10% FBS, 2% B-27 (Thermo Fisher), 1% Glutamax (Thermo Fisher) and 0.25 g/L insulin. At DIV 3-4, 50% of the plating media was removed and exchanged for feeding media containing media supplemented with 4 µM cytosine β-d-arabinofuranoside (Ara-C). Neurons were transfected at DIV 7-9 using the calcium phosphate method. Each coverslip received 1.6 µg of SNAP-mGluR2 and 0.2 µg of tdTomato, both under a CMV promoter. Neurons were identified based on morphology and confirmed by their ability to fire action potentials.

Mixed cultures of cortical and hippocampal astrocytes were prepared from P1–3 mice and plated on poly-D-lysine-coated coverslips (18 mm) in media containing DMEM supplemented with 20% FBS, 25 mM glucose, 2 mM Glutamax, and 1 mM sodium pyruvate. At DIV 3–4, cells were washed to remove debris and media was changed every 3–4 days. Cells were transfected at DIV 7-9 using the calcium phosphate method with 2 µg DNA/well for GCaMP6f and/or SNAP-mGluR2-mGluR5, both under a CMV promoter. 3–5 days after transfection, a subset of coverslips were fixed in 4% paraformaldehyde for immunohistochemical analyses to probe the specificity of GCaMP6f expression for astrocytes relative to other cell types. This was determined by quantification of GCaMP6f colocalization with immunohistochemical markers for astrocytes (glutamine synthetase). Neu-N and Iba1 staining was performed on a subset of coverslips, but no positively labeled cells were observed. >90% of GCaMP6 positive cells showed co-labeling with glutamine synthetase, confirming their identity as astrocytes.

### HEK Cell and Cultured Neuron Electrophysiology

Whole cell patch clamp recordings from HEK 293T cells were performed 24-48 hours after transfection as previously described^27^. Briefly, voltage clamp recordings at −60 mV were performed in a high potassium (120 mM) solution to enable large inward currents upon receptor activation.

Whole cell patch clamp recordings of cultured cortical neurons were performed 4-6 days after transfection (11-15 DIV) in an extracellular solution containing (in mM): 138 NaCl, 1.5 KCl, 1.2 MgCl_2_, 2.5 CaCl_2_, 10 glucose and 5 HEPES, pH 7.4. Intracellular solution contained (in mM): 140 potassium gluconate, 10 NaCl, 5 EGTA, 2 MgCl_2_, 1 CaCl_2_, 10 HEPES, 2 MgATP and 0.3 Na_2_GTP, pH 7.2. For hyperpolarization measurements, cells were adjusted to −60 mV with current injection prior to photoswitching. Only cells with a resting potential ≤ −40 mV were analyzed.

Unless otherwise noted, cells were incubated with 1-10 µM of PORTL for 45-60 min at 37 °C in the appropriate extracellular recording solution. Labeling efficiency was determined as previously described ^10^. Illumination was applied to the entire field of view using a CoolLED pE-4000 through a 40× objective. Light intensity in the sample plane was 1-2 mW/mm^2^. pClamp software was used for both data acquisition and control of illumination.

### Calcium Imaging

HEK 293T cells were imaged at 30 °C in extracellular solution containing (in mM): 135 NaCl, 5.4 KCl, 10 HEPES, 2 CaCl_2_, 1 MgCl_2_, pH = 7.4 with continuous perfusion on a Zeiss LSM880 scanning confocal microscope using ZEN Black software with a 63x objective, a 488 nm laser for imaging and a 405 nm laser for photoactivation. For photoactivation experiments, a brief 10-100 µM glutamate application was used to identify healthy cells, based on calcium responses, which would be suitable for photoactivation experiments. Photoactivation was performed by scanning a 405 nm laser at 100% power in a defined region for 40 iterations between each imaging frame. For drug application experiments, imaging was performed at room temperature on an Olympus IX-73 microscope using a 60x 1.49 NA APO N TIRFM objective (Olympus) and snapshots were taken with a sCMOS ORCA-Flash4 v3.0 camera (Hamamatsu). GFP was excited with a 488 nm laser diode and glutamate was added by a gravity-driven perfusion system.

Astrocytes were imaged at 37 °C with the same protocol described above in an extracellular solution containing (in mM): 138 NaCl, 1.5 KCl, 1.2 MgCl_2_, 2.5 CaCl_2_, 10 glucose and 5 HEPES, pH 7.4. Process were defined as subcellular areas at least 15 µm from the nucleus where the thickness of decreased from that of the soma. ROIs for both analysis and photoactivation were maintained at ∼60 µm^2^. ROIs for photoactivation in processes were placed at least 15-20 µm from the soma. Oscillations were defined as more than one increase in intensity during a single application of UV or drug (at least one minute). DHPG was purchased from Tocris and applied using a gravity-driven perfusion system

Image analysis was performed using ZEN (Zeiss) and ImageJ (Fiji). Intensities were normalized to baseline prior to photoactivation or glutamate application. Oscillation frequency was measured as the number of events per unit time. Ca^2+^ wave velocity was calculated by first determining the time at 50% of the peak for both the photoactivation region of interest (ROI) and a secondary ROI of the same size (5 µm^2^) on the other side of the cell (typically 15-30 µm away from the photoactivation ROI). The wave velocity was then calculated as the distance between the ROIs divided by the difference in time between 50% of the peaks.

### Viral Expression and *in vivo* PORTL labeling

Male C57BL/6J mice were injected at p60 with either 1 µL of a 1:1 viral cocktail of AAV9-EF1a-FLEX-SNAP-mGluR2-WPRE-hGH (Penn Vector Core) and pENN-AAV9-CamKII 0.4-Cre-SV40 (Addgene) or, as a control, only AAV9-EF1a-FLEX-SNAPmGluR2-WPRE-hGH at 500 nL per site targeted to medial prefrontal cortex (AP +1.85, ML +/-0.35, DV −2.0, −1.5) using a Kopf stereotaxic and World Precision Instruments microinjection syringe pump with a 10 µL syringe and 33g blunt needle. For slice recordings, mice received unilateral infusions (ML +0.35). All imaging and slice experiments were performed at least 6 weeks after viral injection. For *in vivo* labeling, mice received infusion of 500 nL of 10 µM BGAG_12_-Cy5 targeted to the same site as viral injection. For imaging of slices, 4 hours later mice underwent transcardial perfusion and were fresh fixed with 4% paraformaldehyde. Brains were extracted and bathed in 4% paraformaldehyde for 24 hours followed by 72 hours in 30% sucrose PBS solution. Brains were mounted and frozen at −20 °C in OCT block and medial prefrontal cortex was sliced at 40 µm thick on a cryostat at −22 °C. Slices were wet mounted to glass slides and secured with coverslip using VECTASHIELD HardSet Antifade Mounting Medium with DAPI (Vector Laboratories). Glass slides were imaged using an Olympus Confocal FV3000 and images were processed in ImageJ.

### Brain Slice Electrophysiology

14–18 hours following BGAG_12_-Cy5 injection, coronal slices of the medial prefrontal cortex were prepared at room temperature in an NMDG-HEPES aCSF containing (in mM): 93 NMDG, 2.5 KCl, 1.2 NaH_2_PO_4_, 30 NaHCO_3_, 20 HEPES, 25 Glucose, 5 sodium ascorbate, 2 thiourea, 3 sodium pyruvate, 10 MgSO_4_, 0.5 CaCl_2_. Slices were maintained in this solution for ∼10 min at 34 °C, and then allowed to recover for at least 45 minutes at room temperature in a modified aCSF containing (in mM): 92 NaCl, 2.5 KCl, 1.2 NaH_2_PO_4_, 30 NaHCO_3_, 20 HEPES, 25 Glucose, 5 sodium ascorbate, 2 thiourea, 3 sodium pyruvate, 2 MgSO_4_, 2 CaCl_2_. Following recovery, slices were transferred to a modified superfusion chamber (Warner Instruments) and mounted on the stage of an Olympus U-TLUIR microscope. Recording were performed at 30 °C in a standard oxygenated aCSF that continually circulated and bubbled with 95% O_2_/5% CO_2_ containing: 124 NaCl, 2.5 KCl, 1.2 NaH_2_PO_4_, 24 NaHCO_3_, 5 HEPES, 12.5 Glucose, 2 MgSO_4_, 2 CaCl_2_

Whole-cell patch-clamp recordings were performed on a computer-controlled amplifier (MultiClamp 700B Axon Instruments, Foster City, CA) and acquired with an Axoscope 1550B (Axon Instruments) at a sampling rate of 10 kHz in current clamp and 500 Hz for current-evoked firing. Pipettes of 2-5 MΩ resistance were filled with a solution containing (in mM): 135 K-Gluconate, 5 NaCl, 2 MgCl_2_, 10 HEPES, 0.6 EGTA, 4 Na2ATP, 0.4 Na2GTP, pH 7.35, 280-290 mOsm. Medial prefrontal cortex pyramidal neurons (primarily in layer V) were identified under visual guidance using infrared-differential interference contrast (IR-DIC) video microscopy through a QImaging optiMOS Camera with a 40× water immersion objective. Illumination for both visualizing BGAG_12_-Cy5 fluorescence and photoswitching was from a CoolLED pE-4000 coupled to the microscope and filtered through an Olympus U-MF2 filter cube with a 635 BrightLine Beamsplitter and 692/40 filter. Light intensity for photoswitching at the sample was 1-2 mW/mm^2^. Neurons were recorded at their resting potential following a minimum 2-minute baseline. For current-induced firing, 250 ms depolarizing current steps were injected with 50 pA increments. All data was analyzed in Clampfit and Prism (Graphpad Software, Inc.).

### Animals

All animal use procedures were performed in accordance with Weill Cornell Medicine Institution Animal Care & Use Committee (IACUC) guidelines under approved protocol (2017-0023). Wild-type mice were of strain C57BL/6J provided by Jackson Laboratory.

## Acknowledgements

The authors thank Andreas Reiner for helpful discussion, Bettina Mathes and Fabio Raith for assistance with chemical synthesis, Zhu Fu and Konstantinos Vlachos for assistance with cloning. The authors acknowledge Scott Blanchard and Roger Altman for providing LD fluorophores for smFRET. JL is supported by an R35 grant (1 R35 GM124731) from NIGMS and the Rohr Family Research Scholar Award. MS is funded by an F32 grant (F32AA025530) from NIAAA. AGO is supported by an R00 grant (R00 AG048222) from the NIA, an Alzheimer’s Association Research Grant, the Leon Levy Fellowship in Neuroscience, and the Kellen Foundation Junior Faculty Fellowship.

## Author Contributions

AAR and Joshua Levitz designed, performed and analyzed all cultured cell electrophysiology experiments. Joon Lee designed, performed and analyzed smFRET experiments. AAR designed, performed and analyzed all cultured cell imaging experiments. SM designed and performed astrocyte experiments. AGO supervised and designed astrocyte experiments. VG designed, performed and analyzed *in vivo* fluorophore labeling. MS and VG designed, performed, and analyzed brain slice electrophysiology experiments. KP designed and supervised brain slice experiments. JB designed and synthesized all chemical compounds. Joshua Levitz contributed to design and analysis of all experiments. JB and Joshua Levitz conceived and oversaw the project and wrote the manuscript with input from all authors.

## Competing Interests

The authors declare no competing interests.

## Supplementary Figure Legends

**Figure S1.**
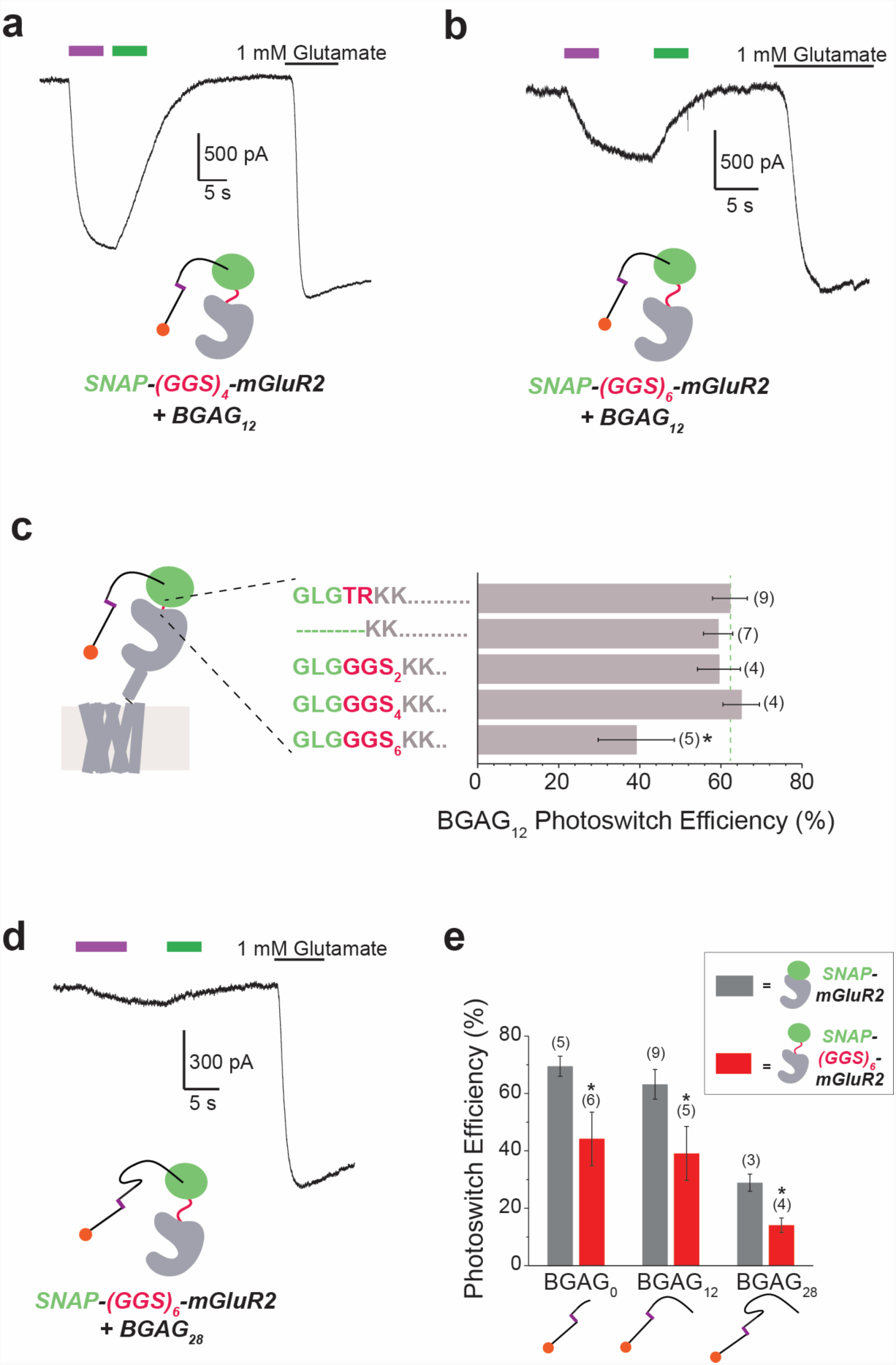
The role of SNAP-mGluR2 linker length in BGAG photoswitching. (**a-c**) SNAP-mGluR2 photoactivation with BGAG_12_ is weakly-sensitive to the composition of the linker-between SNAP and mGluR2. Summary bar graph (c) shows photoswitch efficiency relative to glutamate for various constructs. * indicates statistical significance (unpaired t-test between SNAP-mGluR2 and (GGS)_6_ construct, p=0.01). (**d**) Representative trace showing very weak photoactivation of SNAP-(GGS)_6_- mGluR2 by BGAG_28_. (**e**) Bar graph showing the effect of introducing a flexible (GGS)_6_ linker on SNAP- mGluR2 photoswitching across BGAG lengths. * indicates statistical significance (unpaired t-test between SNAP-mGluR2 and (GGS)_6_ construct, p=0.05 for BGAG_0_ and p= 0.03 for BGAG_28_). The numbers of cells tested are shown in parentheses. Error bars show s.e.m.

**Figure S2.**
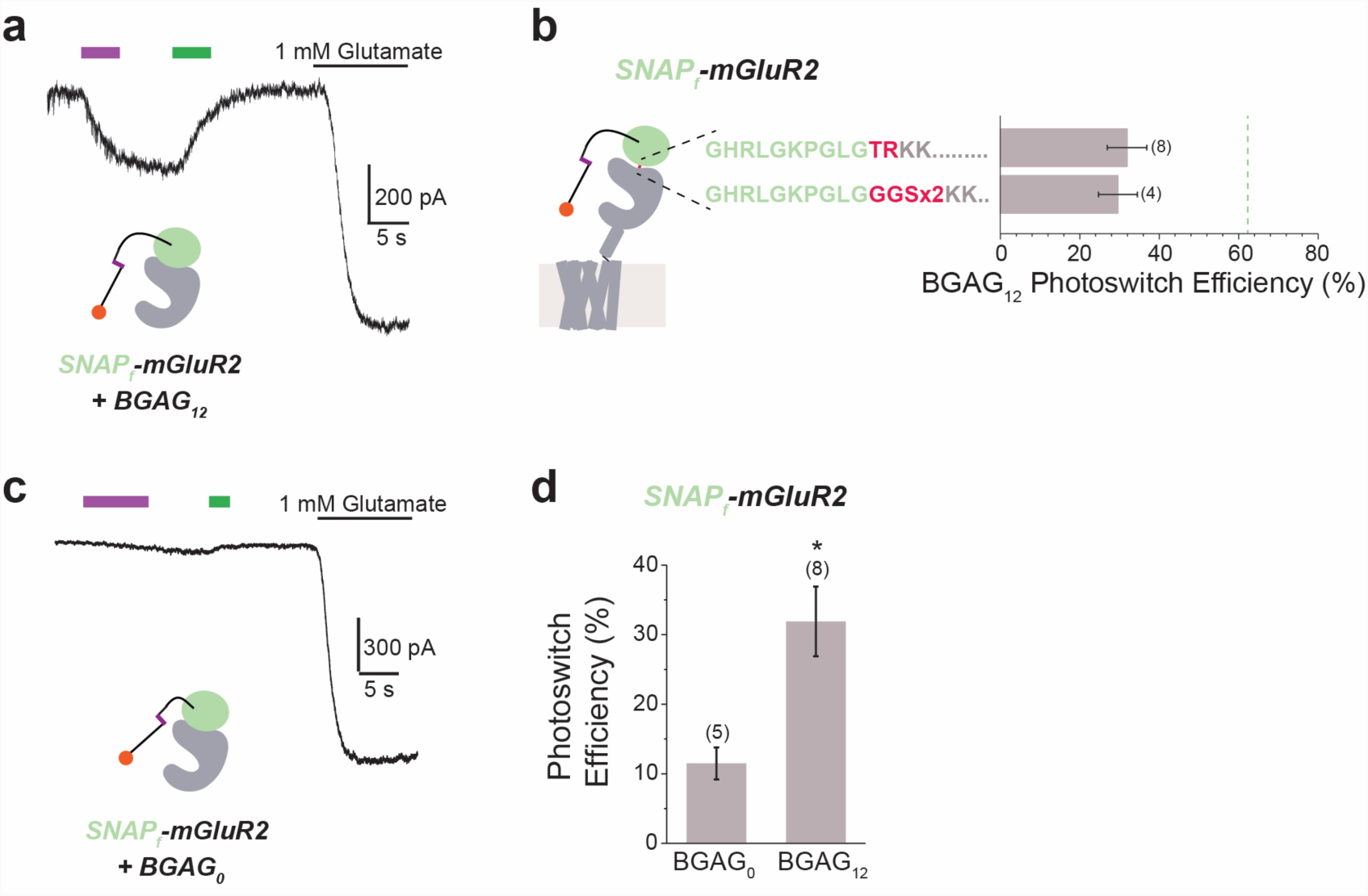
BGAG photoactivation efficiency is substantially decreased with SNAP_fast_-mGluR2 (**a-b**) SNAP-mGluR2 photoactivation by BGAG_12_ is reduced with introduction of the “SNAP_fast_” variant. A summary bar graph (b) shows the photoswitch efficiency for SNAP_f_-mGluR2 with and without introduction of a flexible (GGS)_2_ linker. The efficiency measured with standard SNAP-mGluR2 is shown as a grey dotted line. (**c-d**) The short BGAG_0_ variant is particularly sensitive to incorporation of SNAPfast. * indicates statistical significance (unpaired t-test, p=0.007). The numbers of cells tested are shown in parentheses. Error bars show s.e.m.

**Figure S3.**
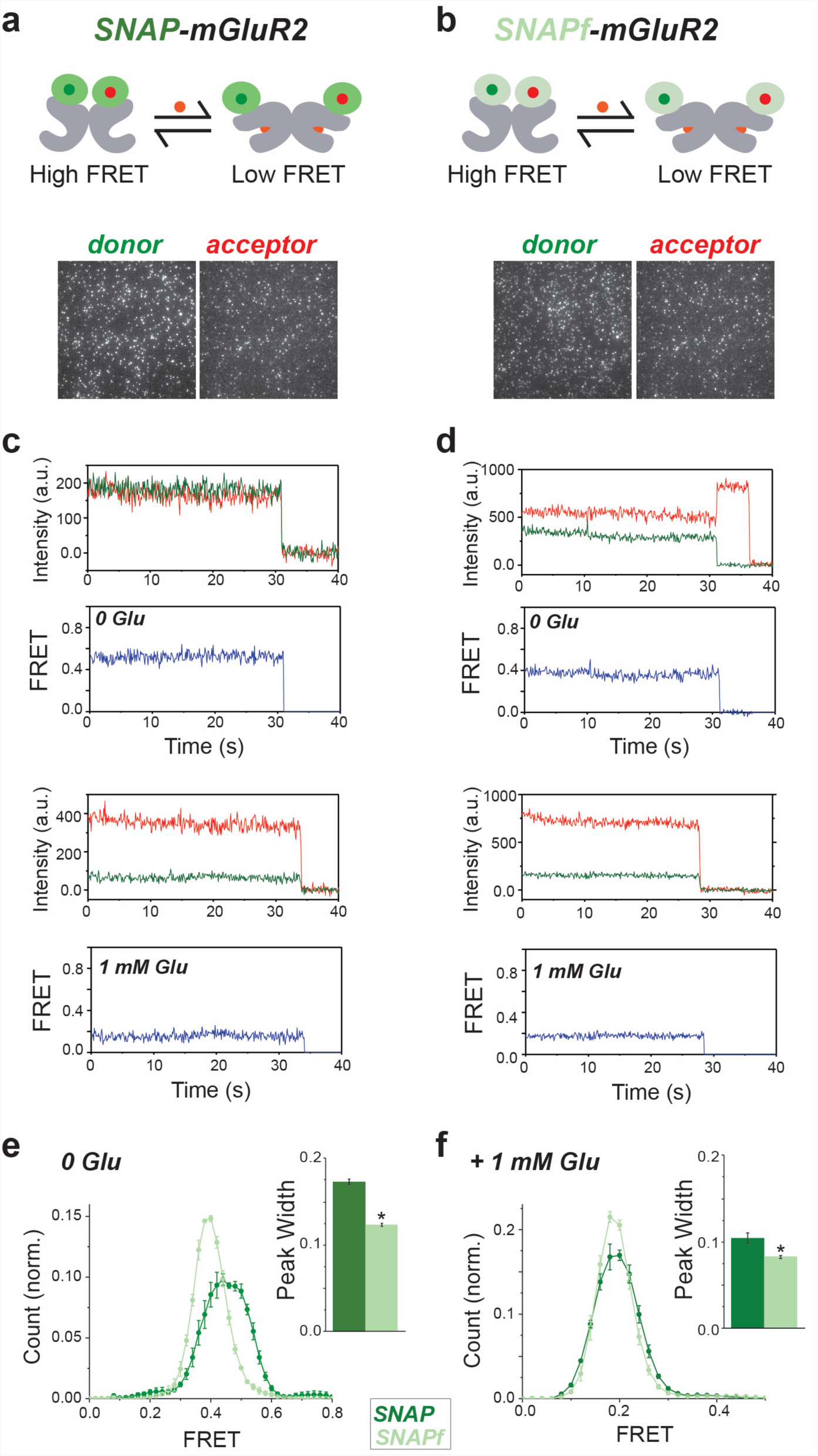
Single molecule FRET analysis reveals decreased flexibility of SNAPf compared to SNAP. (**a-b**) Schematics (a, b; top) and representative images (c, d; bottom) for single molecule imaging of donor and acceptors labeled SNAP-mGluR2 and SNAP_f_-mGluR2. (**c-d**) Representative smFRET traces for individual molecules in the absence (top) or presence of saturating glutamate (bottom). Similar FRET values are seen for SNAP-mGluR2 (e) and SNAP_f_-mGluR2 (f), but the noise thickness is reduced for SNAP_f_. (**e-f**) smFRET histograms in the absence (g) or presence of glutamate (h) show a narrower distribution of FRET values for SNAP_f_, indicative of decreasesd tag flexibility. Inset bar graphs show the peak width, as determined from Gaussian fits, for SNAP versus SNAP_f_. * indicates statistical significance (unpaired t-tests, p= 0.0006 for 0 glutamate and p=0.04 for 1 mM glutamate). Error bars show s.e.m.

**Figure S4.**
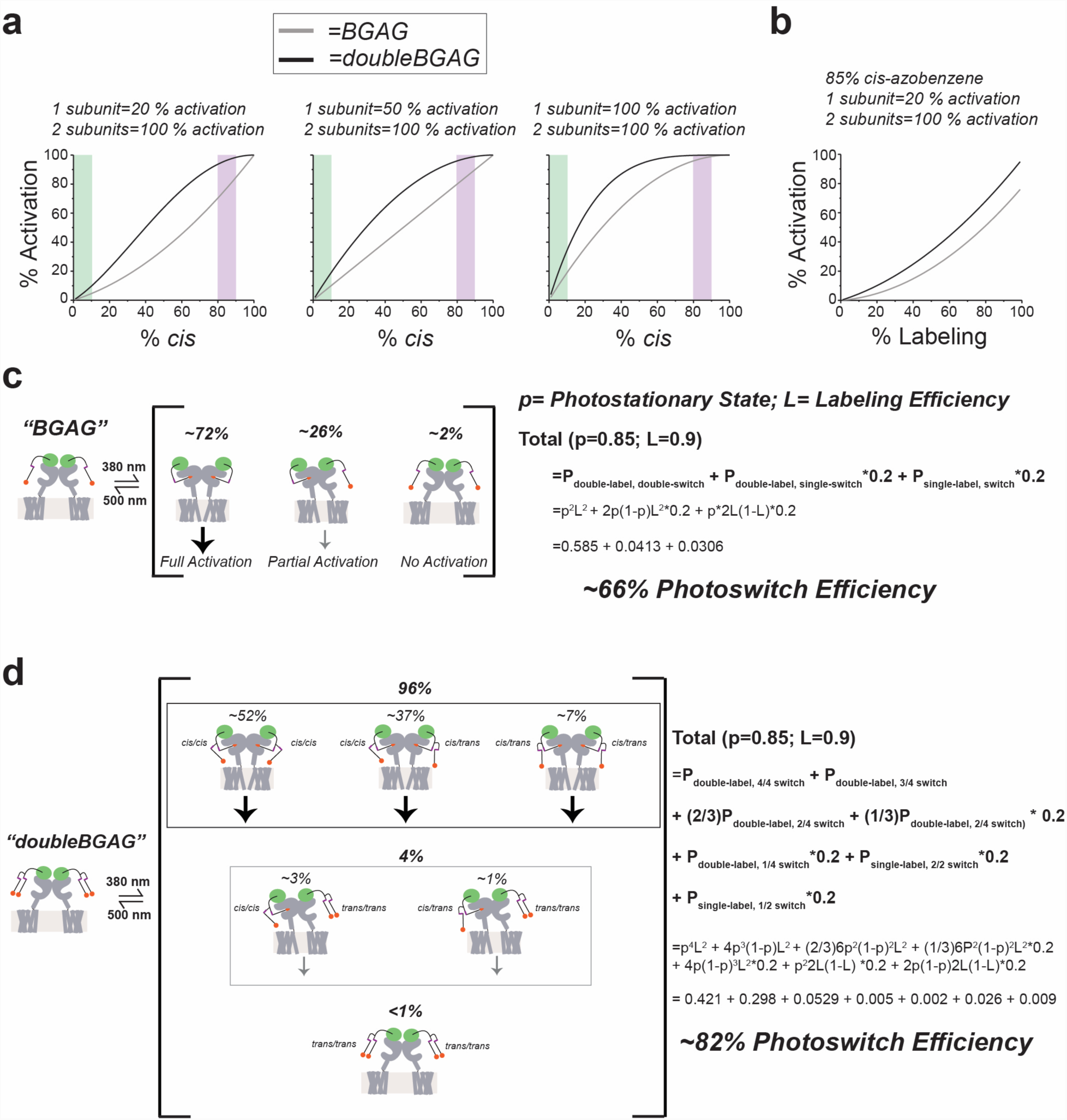
Theoretical rationale for the use of branched ligands to enhance photoswitch efficiency. (**a**) Calculated photoswitch efficiencies for a given proportion of *cis*-azobenzene for dimers of variable cooperativity profiles. Purple and green denote the expected population with 380 nm and >500 nm illumination, respectively. (**b**) Calculated photoswitch efficiency at 380 nm for a dimer with 20% activation with single subunit agonism at different labeling efficiencies. (**c-d**) Expected photoswitch efficiency for BGAG (c) and doubleBGAG (d) with a labeling efficiency of 90% and photostationary state of 85% of azobenzenes in *cis*. Schematic shows the relative population of different combinations of labeled subunits and *cis* versus *trans*-azobenzenes. doubleBGAG is predicted to produce a considerably higher efficiency of activation relative to saturating glutamate.

**Figure S5.**
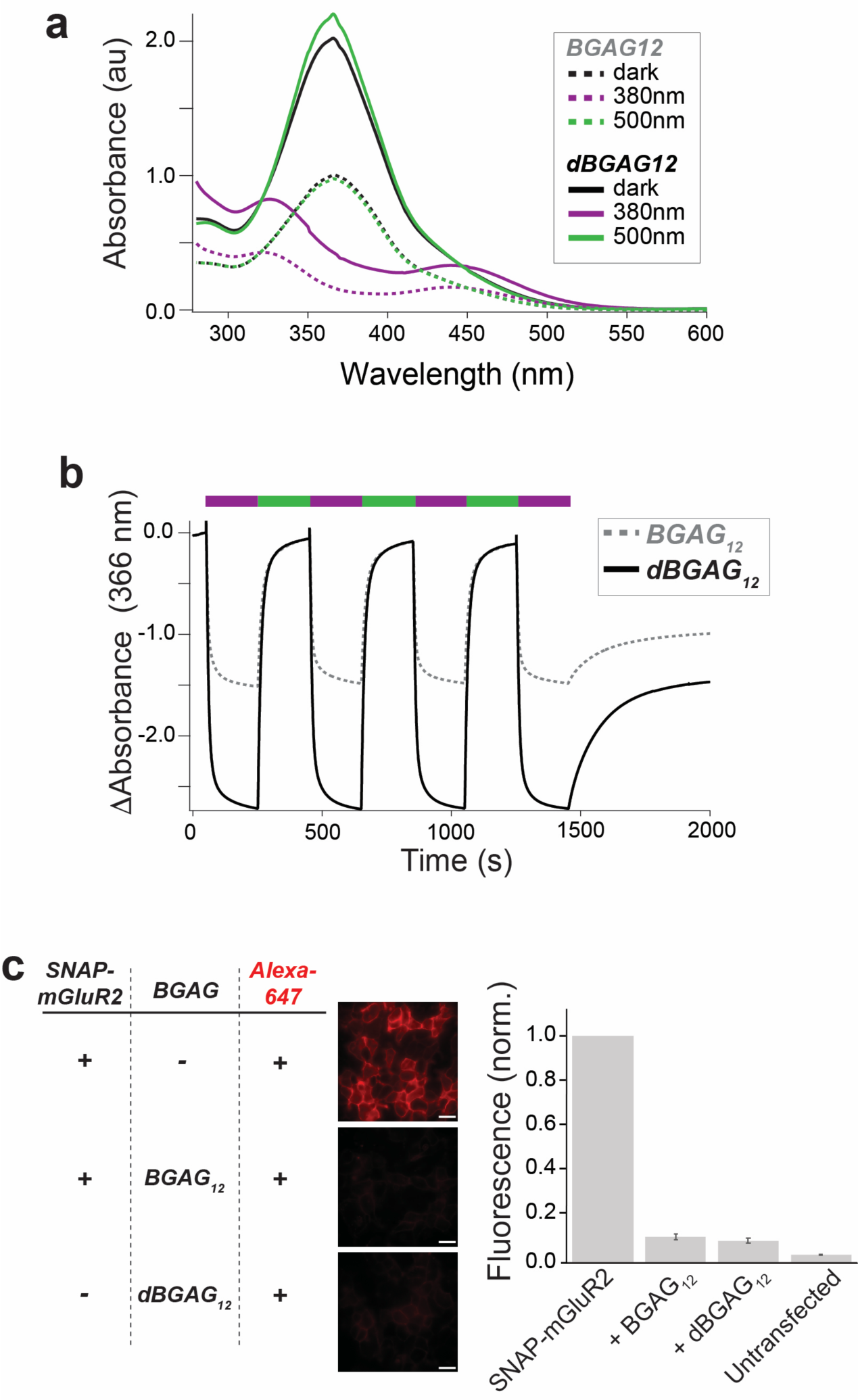
Characterization of branched doubleBGAG_12_: Photophysical properties and labeling efficiency. (**a**) Absorbance spectra determined by UV-Vis spectroscopy for BGAG_12_ and dBGAG_12_ in PBS. (**b**) *in vitro* photoswitching of BGAG_12_ and dBGAG_12_ in PBS. (**c**) Competition-based labeling assay reveals similar ∼90% labeling efficiencies for both BGAG_12_ and dBGAG_12_ at 1 μM.

**Figure S6.**
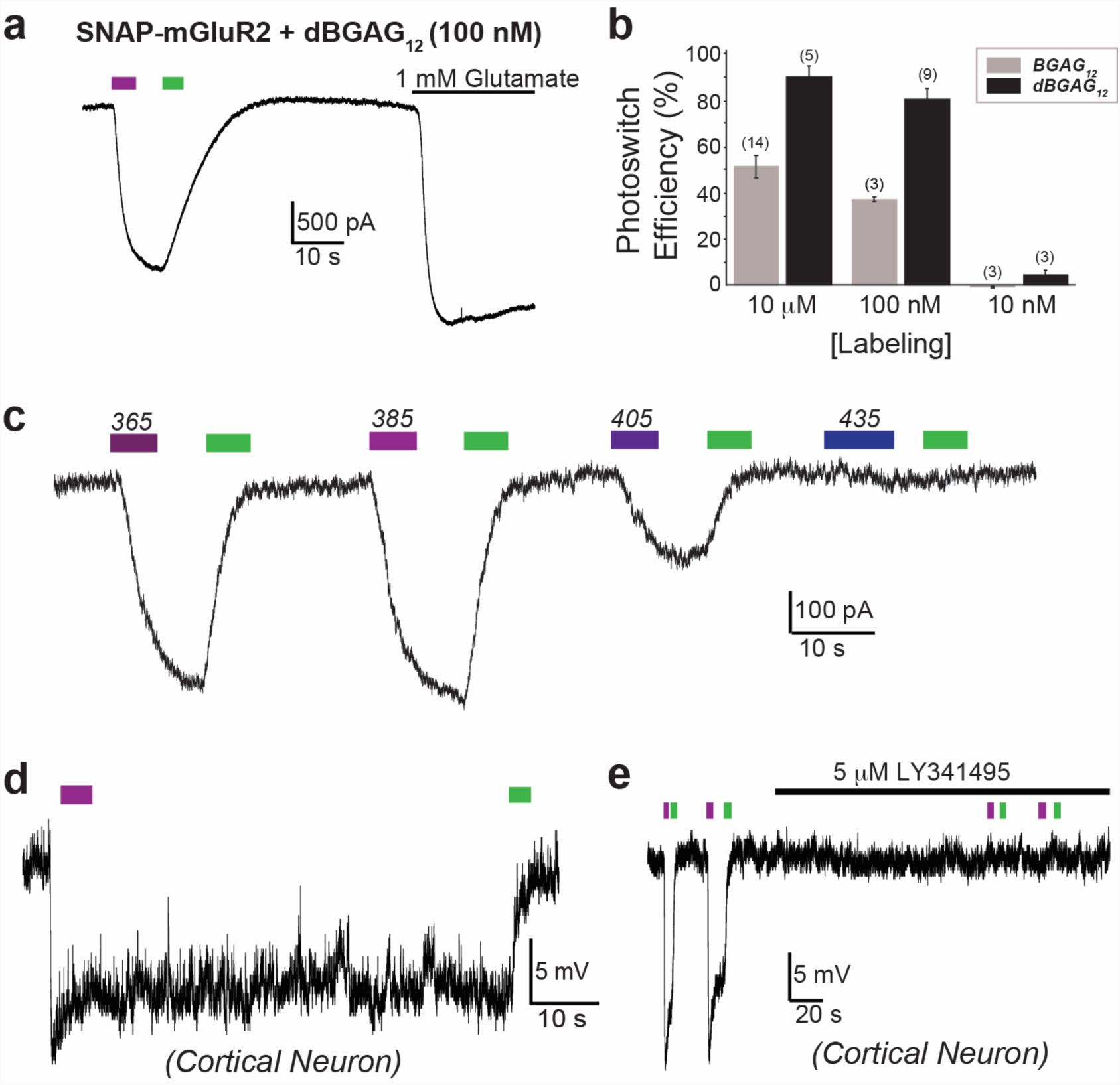
Further analysis of doubleBGAG_12_ photoswitching of SNAP-mGluR2. (**a-b**) Labeling concentration-dependence shows enhanced photoswitching for dBGAG12 versus BGAG12 across a range of concentrations. A representative photoswitching trace (a) for dBGAG_12_ on SNAP-mGluR2 following labeling at 100 nM shows high efficiency. A summary bar graph (b) shows enhanced photoswitching for dBGAG_12_ at all labeling concentrations tested. The numbers of cells tested are shown in parentheses. Error bars show s.e.m. (**c**) Whole cell patch trace showing spectral sensitivity of dBGAG12 on SNAP-mGluR2. Optimal photoactivation is seen with 385 nm illumination, but clear light responses are seen with 365 nm and 405 nm illumination. (**d-e**) Photoactivation of dBGAG_12_ in cultured neurons is bistable over the minute time scale (d) and blocked by the competitive mGluR2 antagonist, LY341495 (e).

**Figure S7.**
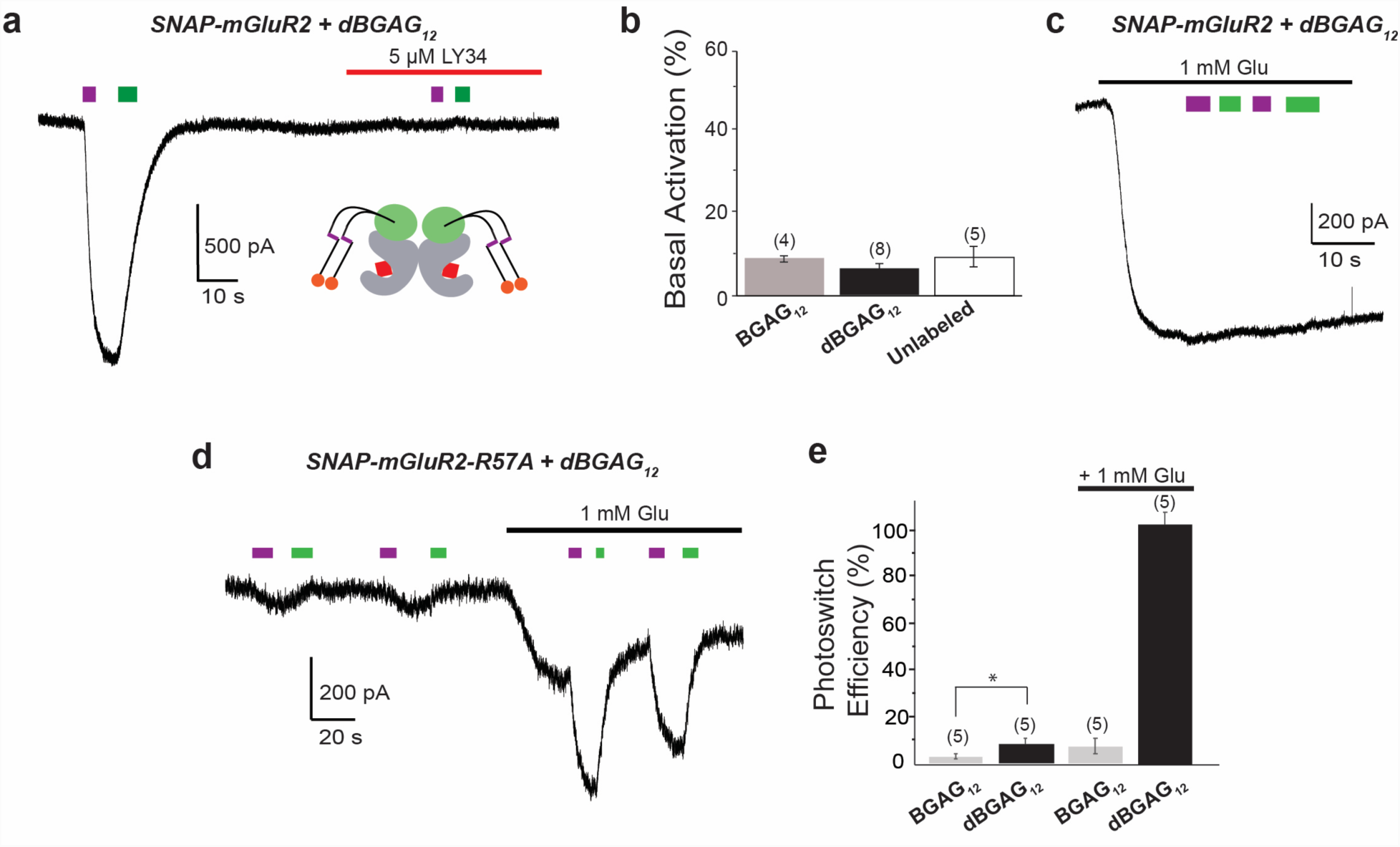
Mechanistic characterization of doubleBGAG photoswitching of SNAP-mGluR2: ligand binding properties. (**a-b**) *trans*-dBGAG12 does not increase basal activation of mGluR2. Representative trace shows that application of saturating LY341495 blocks photoactivation without substantially altering the baseline current level (a) and summary bar graph shows a similarly small effect in the absence or presence of BGAG_12_ or dBGAG_12_. (**c**) Photoswitching dBGAG_12_ in the presence of saturating glutamate does not lead to antagonism, consistent with azobenzene-glutamates functioning as saturating, not partial, agonists. (**d-e**) Photoswitching low affinity SNAP-mGluR2-R57A reveals enhanced effective concentration of dBGAG_12_ relative to BGAG_12_. In the absence or presence of glutamate, photocurrents are observed for dBGAG_12_ but not BGAG_12_. * indicates statistical significance (unpaired t-test, p= 0.01). The numbers of cells tested are shown in parentheses. Error bars show s.e.m.

**Figure S8.**
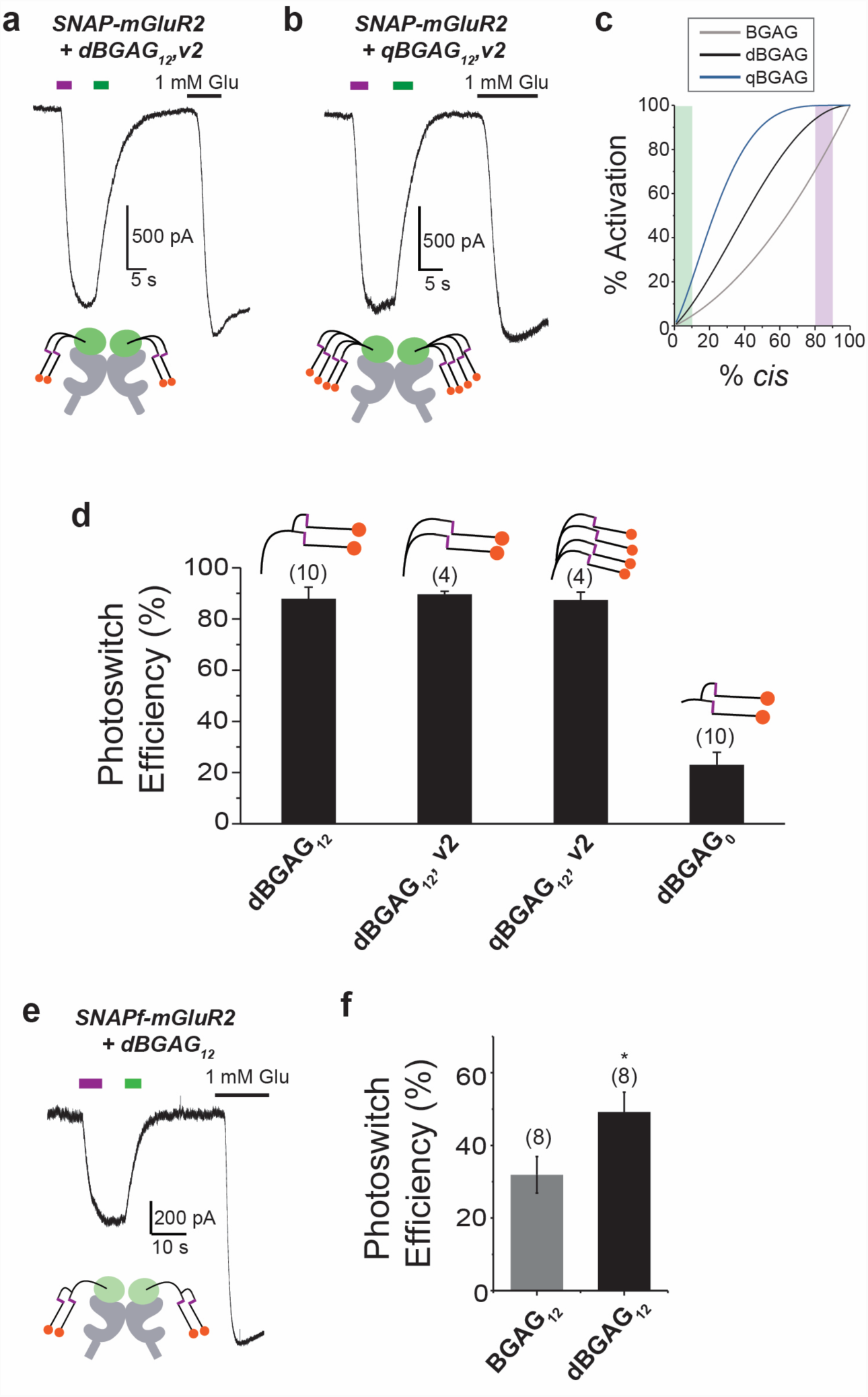
Mechanistic characterization of doubleBGAG photoswitching of SNAP-mGluR2: branching properties and SNAP variants. (**a-d**) Photoswitching of SNAP-mGluR2 with both dBGAG_12,v2_ (a) and qBGAG_12_ (b) shows high-efficiency photoactivation, consistent with theoretical model showing proportion of *cis*-azobenzene and resulting expected photoactivation (c). Summary bar graphs shows photoswitch efficiency for SNAP-mGluR2 with dBGAG_12,v2_, qBGAG_12_, and dBGAG_0_ (d). (**e-f**) SNAPf-mGluR2 shows improved photoswitching efficiency with branched dBGAG_12_ photoswitch (e) compared to equivalent single chain BGAG variant (f). The numbers of cells tested are shown in parentheses. Error bars show s.e.m. * indicates statistical significance (unpaired t-test, p= 0.04).

**Figure S9.**
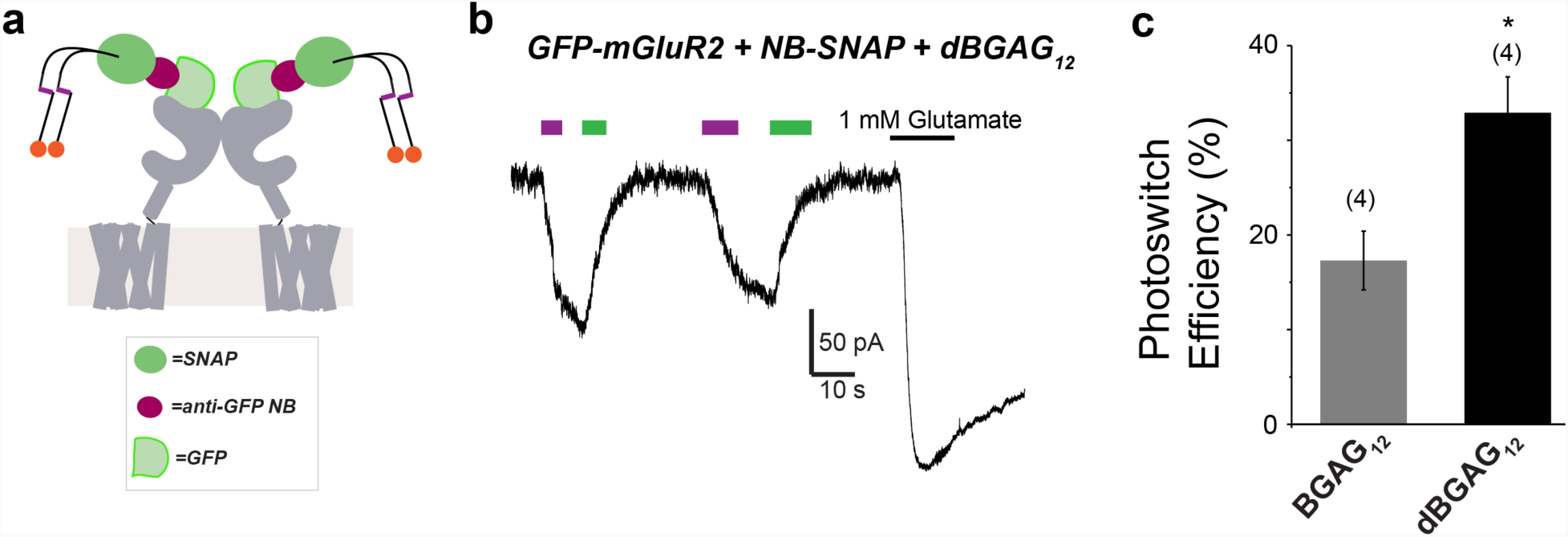
BGAG branching enhances nanobody-mediated mGluR2 photoactivation. (a) Schematic showing GFP-mGluR2 bound with anti-GFP SNAP-tagged nanobody labeled with dBGAG_12_. (b) Representative trace shows significant improvement in photoswitching efficiency (b) compared to equivalent single chain BGAG variant (c). * indicates statistical significance (unpaired t-test, p=0.04). The numbers of cells tested are shown in parentheses. Error bars show s.e.m.

**Figure S10.**
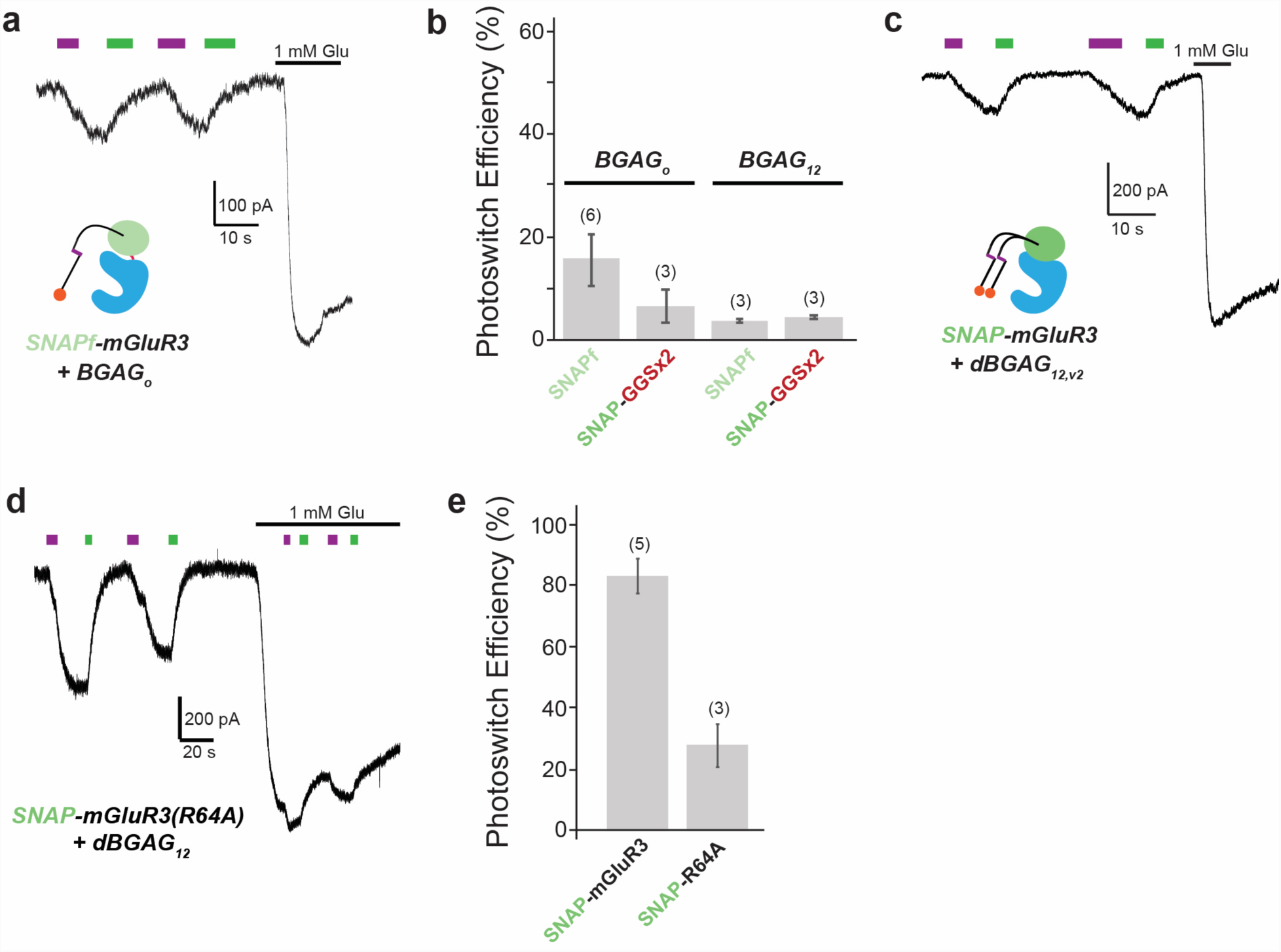
Branched BGAGs enable efficient optical control of mGluR3: further analysis. (**a-b**) Single chain variants do not improve mGluR3 photoactivation when modifications to SNAP flexibility or linker size are introduced. Neither SNAP_f_-mGluR3 (a) or SNAP-(GGS)_2_-mGluR3 showed improved photoswitching with BGAG_0_ or BGAG_12_ compared to SNAP-mGluR3 (b). (**c**) Branch point controls doubleBGAG efficiency on SNAP-mGluR3. Representative trace showing that SNAP-mGluR3 labeled with dBGAG_12,v2_ shows decreases photoswitching efficiency relative to dBGAG_12_. (c). (**d-e**) SNAP-mGluR3 (R64A) labeled with dBGAG_12_ (d) shows decreased photoactivation compared to SNAP-mGluR3 (e), demonstrating the requirement of an intact, high-affinity glutamate binding site. The numbers of cells tested are shown in parentheses. Error bars show s.e.m.

**Figure S11.**
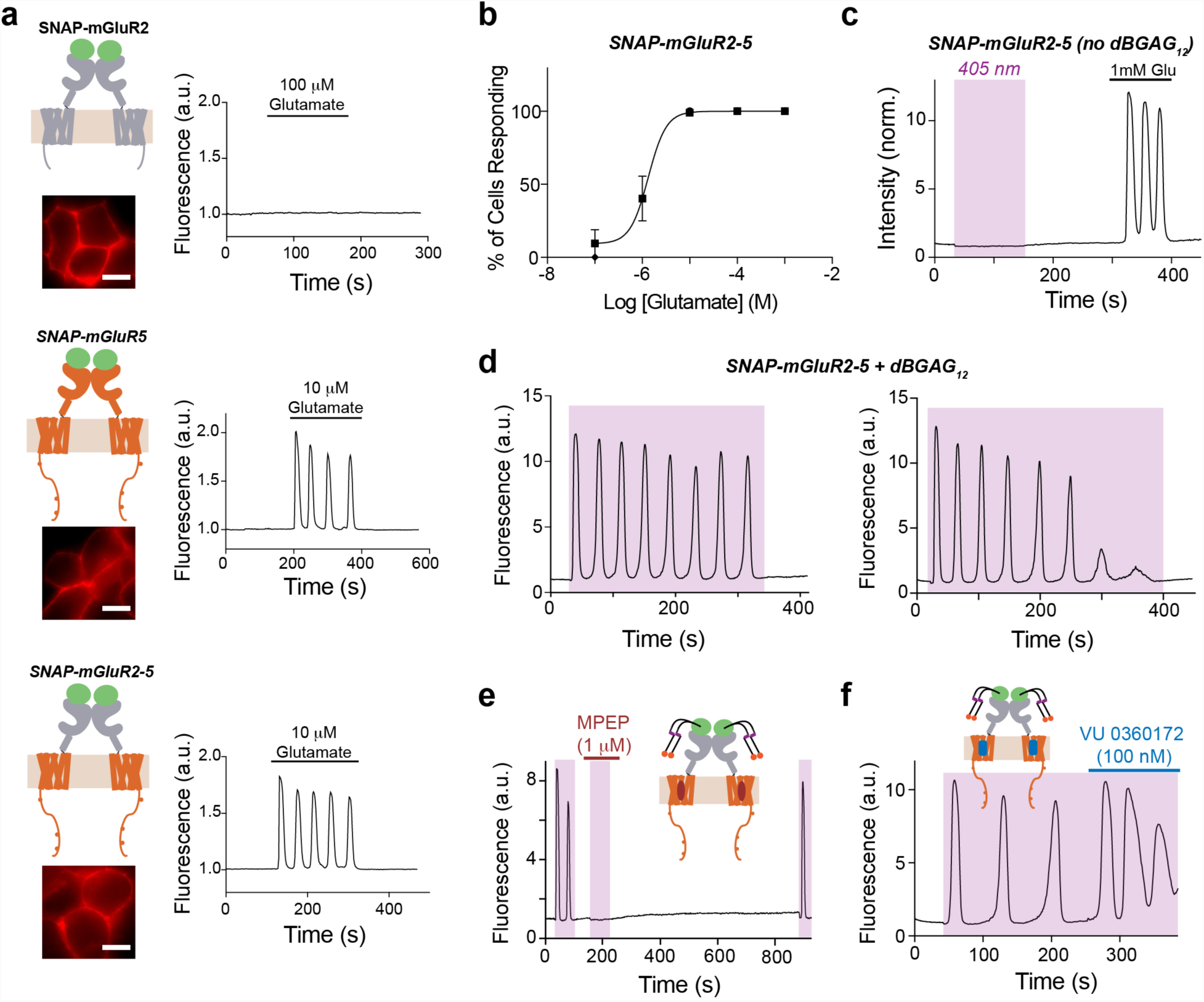
Further analysis of optical control of mGluR5 signaling with branched BGAGs. (**a**) Schematics and representative images, left, for SNAP-mGluR2, SNAP-mGluR5, and SNAP-mGluR2-5. Representative GCaMP6f calcium imaging traces for individual cells showing the response to glutamate application. Scale bars= 10 μm. (**b**) Glutamate dose-response curve for SNAP-mGluR2-5. For a given glutamate application, the percentage of cells responding was calculated. EC_50_= 1.3 ± 0.25 μM. Data comes from n=119 cells over 2 separate experiments. (**c**) Representative calcium imaging trace showing that no response to 405 nm (violet) is seen in cells expressing SNAP-mGluR2-5 but not labeled with dBGAG_12_. (**d**) Representative calcium imaging traces showing responses to extended application of 405 nm illumination. In some cells (21/37) calcium oscillations are sustained with minimal decay (left) and in others (16/37) attenuation is observed (right). (**e-f**) dBGAG_12_-mediated light responses of SNAP-mGluR2-5 are sensitive to an mGluR5 NAM (e) or PAM (f).

**Figure S12.**
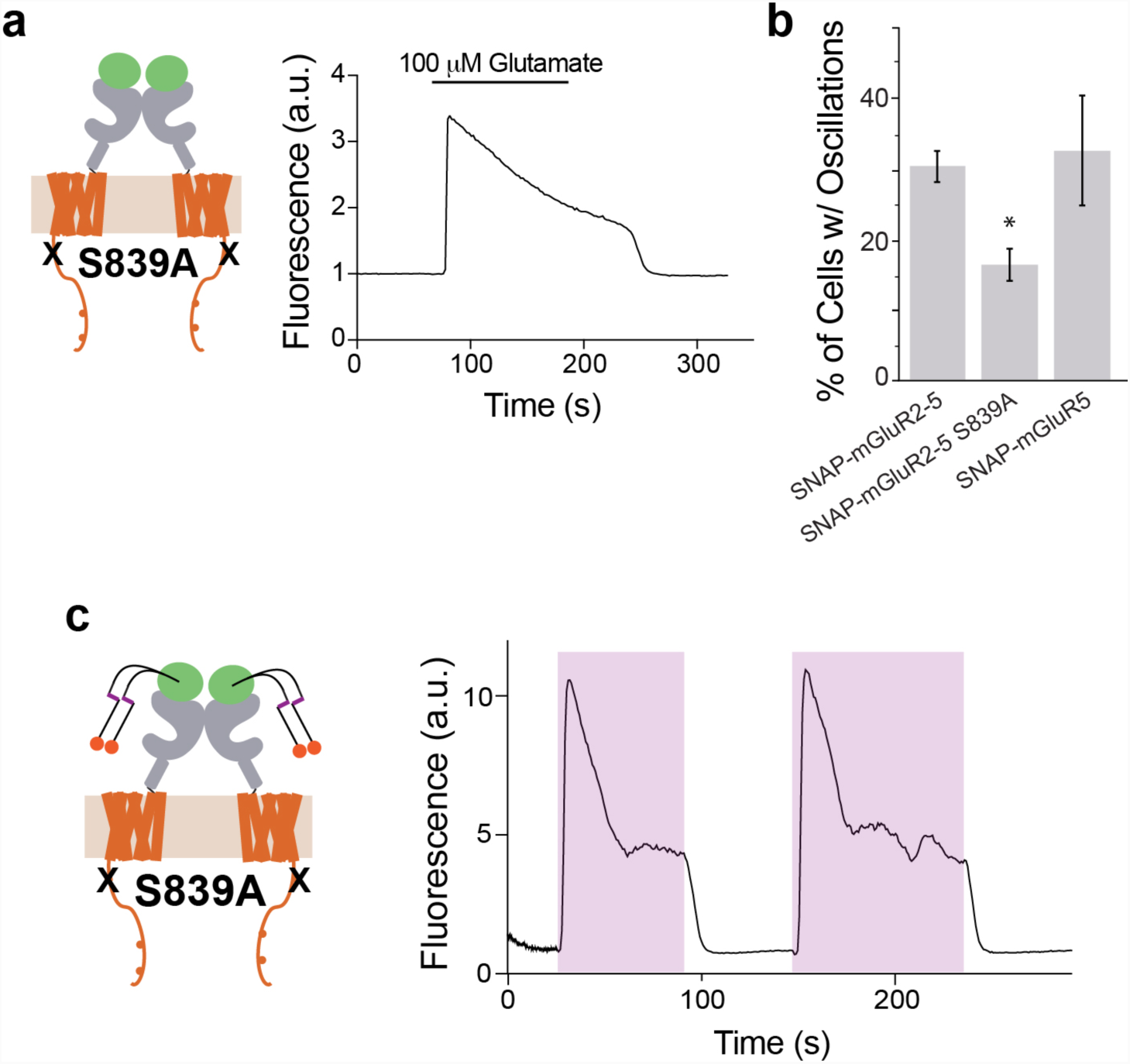
Analysis of a protein kinase C phosphorylation site mutant in the CTD of mGluR5. (**a-b**) Introduction of a mutation at a protein kinase C site (S839A) in the membrane-proximal C-terminus of mGluR5 significantly reduces the % of cells responding to glutamate with calcium oscillations. Data comes from n>100 cells per condition from 3-4 separate experiments. * indicates statistical significance (unpaired t-test for S839A versus SNAP-mGluR2-5, p=0.02). Note that the percentage of cells showing calcium oscillations is similar between SNAP-mGluR5 and SNAP-mGluR2-5 and that the S839A does not completely abolish oscillatory responses. (c) Representative trace showing that following introduction of S839A, SNAP-mGluR2-5 mediated light-induced oscillations are converted into slowly-desensitizing calcium responses typical of most G_q_-coupled receptors.

**Figure S13.**
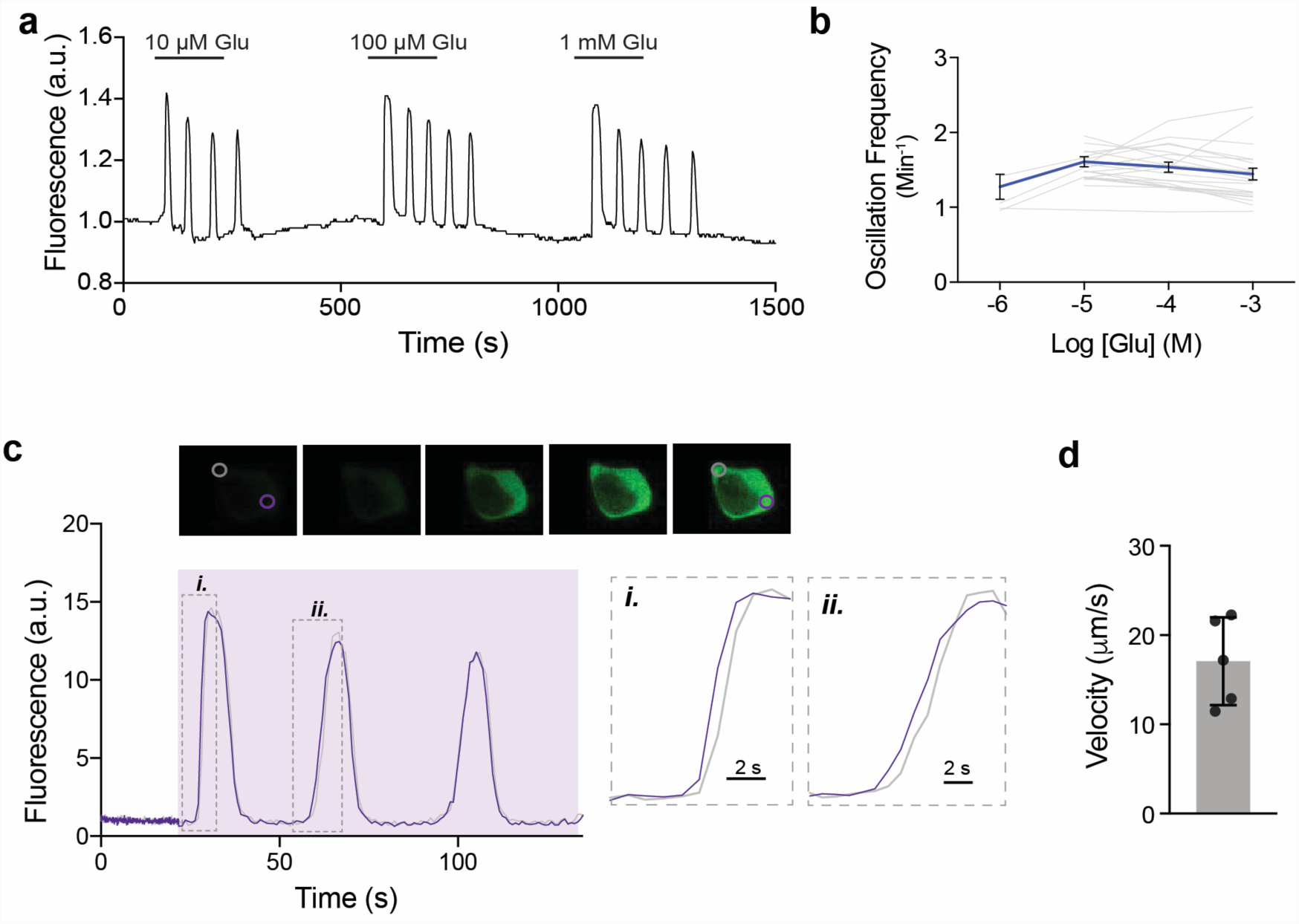
Further analysis of mGluR5-induced calcium oscillations. (**a-b**) The frequency of calcium oscillations induced by SNAP-mGluR2-5 is independent of agonist concentration. Representative trace (a) and dose response plot (b) show consistent oscillations across a range of glutamate concentrations. (**c-d**) Targeted photoactivation allows calculation of intracellular Ca^2+^ wave velocity. Representative trace shows photoactivation of SNAP-mGluR2-5 with dBGAG_12_ demonstrating an offset in Ca^2+^ response timing at two distinct ROIs, the photoactivation site in purple and a distal ROI in gray (c). Bar graph shows quantification of Ca^2+^ wave velocity (d), where n=5 cells and error bars shows s.e.m.

**Figure S14.**
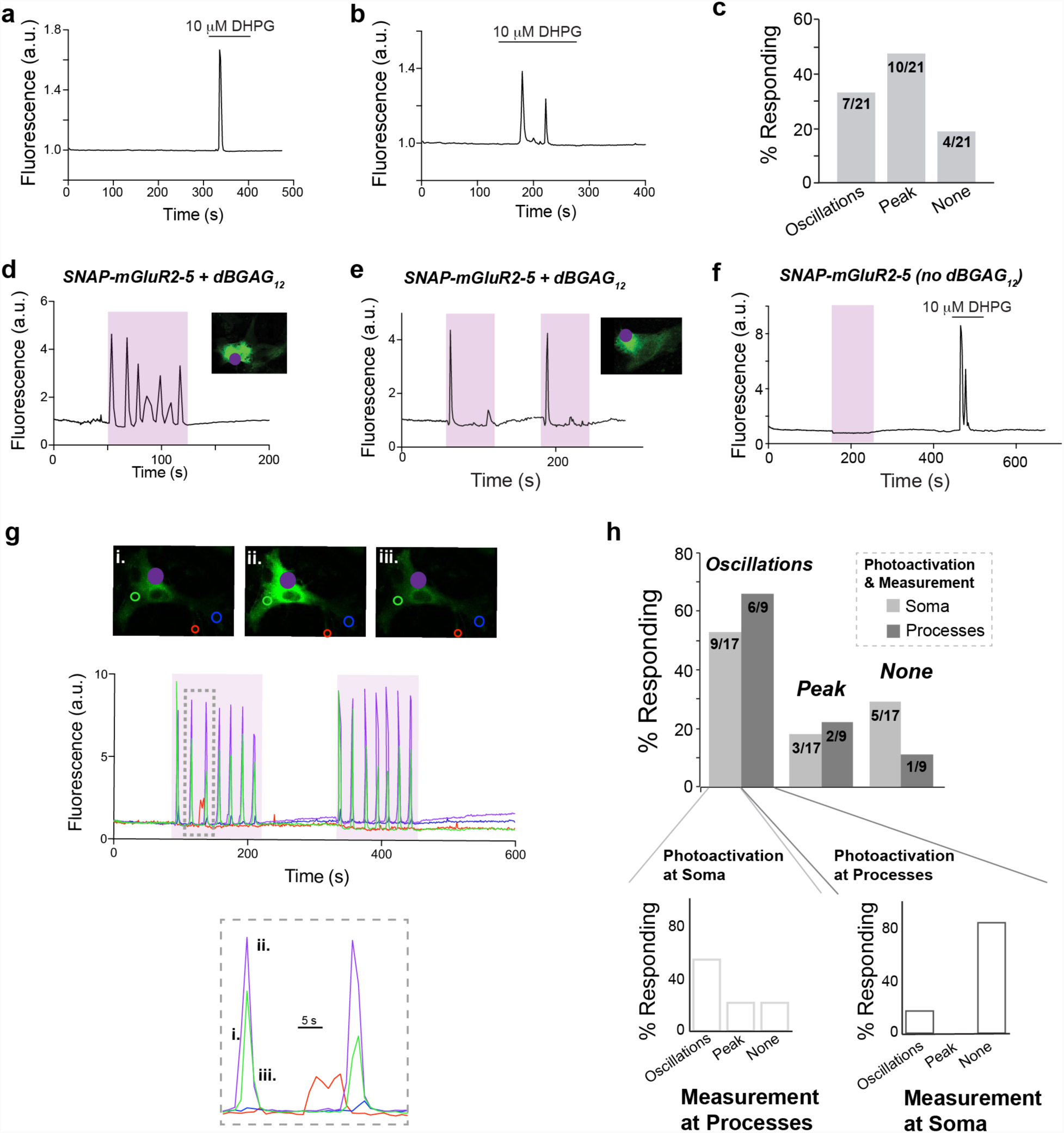
Further analysis of optical control of mGluR5 signaling in cultured astrocyte. (**a-c**) Shows the variability of responses in cultured astrocytes to DHPG, an mGluR5 agonist, representing the native mGluR5 activation response. Native mGluR5 activation results in either a single peak (a) or as calcium oscillations (b). A summary bar graph (c) shows the relative proportion of cells responding with either type of response. (**d-e**) Photoactivation of SNAP-mGluR2-5 with dBGAG_12_ in astrocytes shows either oscillatory responses (d) or single peak responses (e). (**f**) Light-induced calcium responses via SNAP-mGluR2-5 are dependent on labeling with dBGAG_12_ (e). (**g**) Spatial targeting afforded by dBGAG12 allows analysis of subcellular heterogeneity of calcium responses. Photoactivation at an astrocytic process (purple dot), drives oscillations at the photoactivation area, but Ca^2+^ does not spread to somatic subcellular areas (red and blue circles), shown in consecutive images (top) and representative trace (middle). Inset (bottom) shows oscillations, at the processes but not soma, during the second UV application. (**h**) Bar graphs shows the range of calcium responses, organized by site of photoactivation and measurement. The number of cells measured are shown inscribed in each bar.

**Figure S15.**
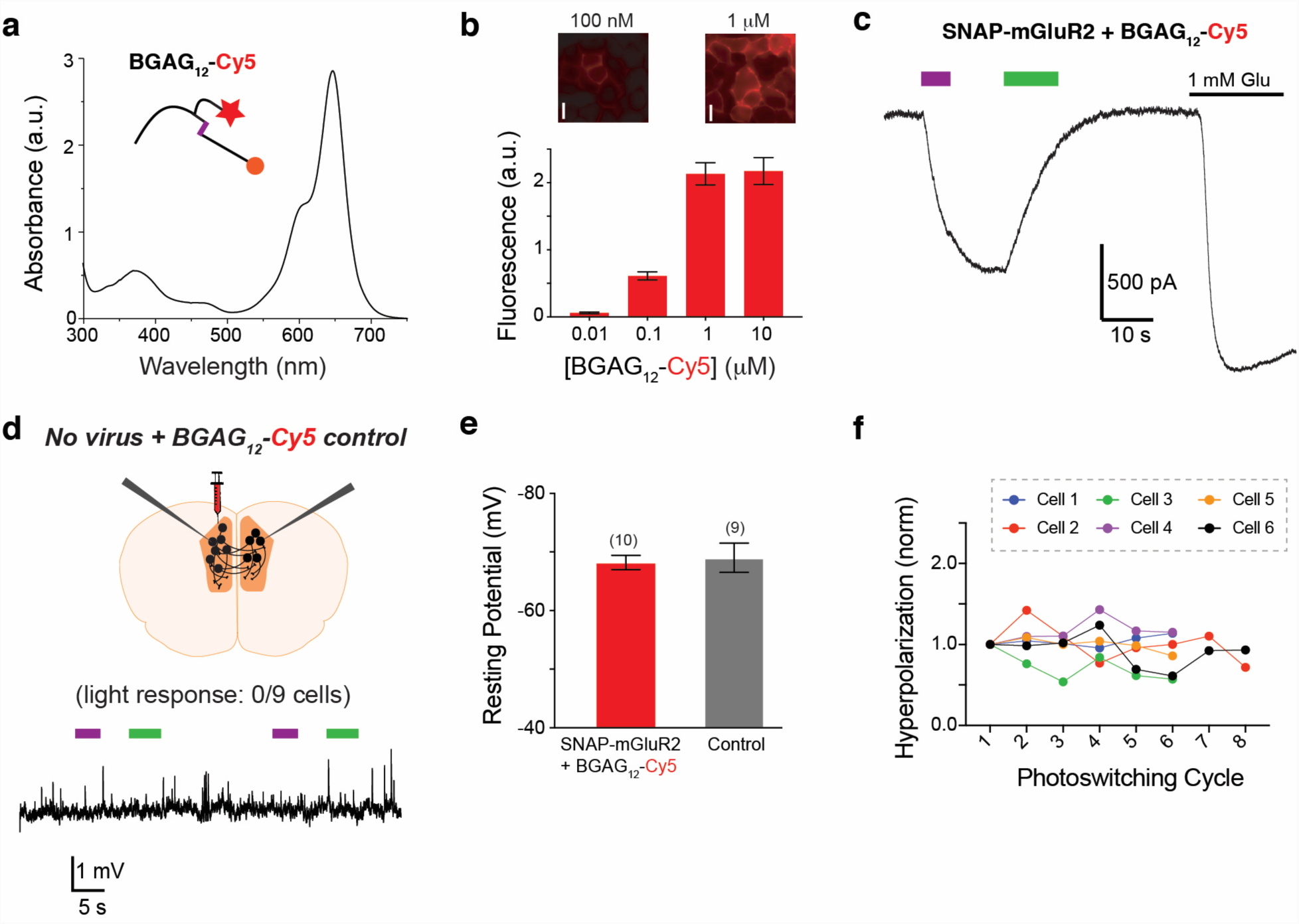
Further analysis of the branched, fluorophore-containing BGAG_12_-Cy5. (**a**) Absorbance spectrum for BGAG_12_-Cy5 showing clear absorption peaks for azobenzene (∼365 nm) and for Cy5 (∼640 nm). (**b**) Labeling of SNAP-mGluR2 with BGAG_12_-Cy5 shows a similar concentration-dependence to previously-described BGAGs, as assessed by Cy5 fluorescence. 1 µM labeling at 37° C for 45 minutes is sufficient for maximal labeling and fluorescence is confined to the cell surface, confirming that BGAG_12_-Cy5 is membrane impermeable. (**c**) Representative trace showing large photocurrents from HEK 293T cells labeled with BGAG_12_-Cy5. (**d**) Schematic and representative trace showing that no light-induced effect on membrane potential is observed in slices from wild-type mice injected with BGAG_12_-Cy5. (**e**) Bar graph summarizing the resting potential of fluorescent cells which express SNAP-mGluR2 and are labeled with BGAG12-Cy5 compared to control cells. The numbers of cells tested are shown in parentheses. Error bars show s.e.m. (**f**) Plots showing the repeatability of SNAP-mGluR2 light-induced hyperpolarization over multiple cycles of activation for individual cells.

