## Supplemental Information for "Branched photoswitchable tethered ligands enable ultra-efficient optical control and detection of class C G protein-coupled receptors"

### 1. *Chemical Synthesis*

#### 1.1. General

Solvents for chromatography and reactions were purchased dry over molecular sieves or in HPLC grade. Unless otherwise stated, all other reagents were used without further purification from commercial sources. LC-MS was performed on a Shimadzu MS2020 connected to a Nexera UHPLC system equipped with a Waters ACQUITY UPLC BEH C18 (1.7 µm, 50 × 2.1 mm). Buffer A: 0.1% FA in $H_2O$ Buffer B: acetonitrile. The typical gradient was from 10% B for 0.5 min → gradient to 90% B over 4.5 min → 90% B for 0.5 min → gradient to 99% B over 0.5 min with 1 mL/min flow. Retention times ($t_R$) are given in minutes (min).

High resolution mass spectrometry was performed using a Bruker maXis II ETD hyphenated with a Shimadzu Nexera system. The instruments were controlled *via* Brukers otofControl 4.1 and Hystar 4.1 SR2 (4.1.31.1) software. The acquisition rate was set to 3 Hz and the following source parameters were used for positive mode electrospray ionization: End plate offset = 500 V; capillary voltage = 3800 V; nebulizer gas pressure = 45 psi; dry gas flow = 10 L/min; dry temperature = 250 °C. Transfer, quadrupole and collision cell settings are mass range dependent and were fine-adjusted with consideration of the respective analyte's molecular weight. For internal calibration sodium format clusters were used. Samples were desalted *via* fast liquid chromatography. A Supelco Titan™ C18 UHPLC Column, 1.9 µm, 80 Å pore size, 20 × 2.1 mm and a 2 min gradient from 10 to 98% aqueous MeCN with 0.1% FA ($H_2O$: Carl Roth GmbH + Co. KG ROTISOLV® Ultra LC-MS; MeCN: Merck KGaA LiChrosolv® Acetonitrile hypergrade for LC-MS; FA - Merck KGaA LiChropur® Formic acid 98%- 100% for LC-MS) was used for separation. Sample dilution in 10% aqueous ACN (hyper grade) and injection volumes were chosen dependent of the analyte's ionization efficiency. Hence, on-column loadings resulted betwee 0.25–5.0 ng. Automated internal re-calibration and data analysis of the recorded spectra were performed with Bruker's DataAnalysis 4.4 SR1 software.

Preparative RP-HPLC was performed on a Waters e2695 system equipped with a 2998 PDA detector for product collection (at 220, 280, 360 or 460 nm) on a Supelco Ascentis® C18 HPLC Column (5 µm, 250 × 21.2 mm). Buffer A: 0.1% TFA in $H_2O$ Buffer B: acetonitrile. The typical gradient was from 10% B for 5 min → gradient to 90% B over 45 min → 90% B for 5 min → gradient to 99% B over 5 min with 8 mL/min flow.

Compounds **1**, **2** and **BG-COOH**, **BC-DBCO** were previously described in Broichhagen *et al.*, *ACS Cent. Sci.* **2015**, *1*, 383–393 and Levitz et al., *PNAS* **2017**, *114*, E3546–E3554, respectively.

#### 1.2. Abbreviations

DIPEA: *N*,*N*-di*iso*propylethylamine; DBU: 1,8-diazabicyclo[5.4.0]undec-7-ene; DMF: *N*,*N*-dimethylformamide; DMSO: dimethylsulfoxide; FA: formic acid; Su: succinimidyl; TFA: trifluoroacetic acid; TSTU: *O*-(*N*-succinimidyl)-*N*,*N*,*N*′,*N*′-tetramethyluronium tetrafluoroborate.

#### 1.3. Notes and observations

NHS ester stability: TSTU was the coupling reagent of choice used for the synthesis of BGAGs, converting an acid to its respective NHS-ester usually within minutes. While most NHS esters were used *in situ* without purification, they can be isolated by RP-HPLC (*cf.* compound **4**) and immediate lyophilization. Aliquoting and storage at –20 °C is recommended to avoid repeated freeze-thaw cycles that lead to decomposition.

Fmoc deprotection: Fmoc is a standard amine protecting group extensively used in solid phase peptide synthesis, where amide couplings (with activating agents in DMF) and subsequent deprotection (with piperidine in DMF) is performed iteratively in high yields. Inspired by this and with the aim to reduce labor and purification steps, peptide couplings were performed in DMF with TSTU as an activating agent, and after amide coupling was complete, 5 vol% of piperidine was added directly to the reaction mixture. This proved to work reliably in our hands with all blue-shifted azobenzene compounds based on structure **1**, but lead to complex reaction mixtures when using this method with red-shifted azobenzene compounds based on structure **2**. Why the reason for this was not further investigated, we chose to purify Fmoc-containing compounds by RP-HPLC mainly to remove DMF, DIPEA and urea side products from TSTU, and employed DBU in MeCN as a deprotection reagent. Indeed, this was tolerated very well by red-shifted compounds and is noted when used in the procedures below.

Stability of BG, BC and Halo-congeners towards acid: BGAGs need final deprotection of the NHBoc group to the free amine and TFA is the deprotecting agent of choice, however, the *O*-benzylated guanine and cytosine bases were shown to be labile towards such strong acids. As such, we investigated and found that TFA can be used with BG-containing compounds if kept on ice with pre-cooled TFA, and its removal is not done in a rotary evaporator but by applying a gentle stream of nitrogen in a well ventilated chemical hood. Unfortunately, BC-containing compounds to not survive this treatment unharmed, and this is the reason why the NHBoc group is deprotected beforehand and strain promoted alkyne azide click reaction is performed in another orthogonal way. The Halo-group, however, is inert towards neat TFA at r.t..

#### 1.4. General protocol to generate NHS esters

A 1 mL vial was charged with 1.0 equiv. of acid dissolved in DMF (1 mL / 10 mg) and 4.0 equiv. of DIPEA was added before 1.1 equiv. of TSTU in one portion (for amounts <1 mg of TSTU, stock solutions were prepared as it is critical to not overload TSTU). The active NHS ester was allowed to form for 15 min and used without further purification.

##### 1.5. General procedure for peptide couplings and *in situ* Fmoc deprotection

A 1 mL vial was charged with 1.0 equiv. amine dissolved in DMF (1 mL / 10 mg) and 4.0 equiv. DIPEA. The pre-formed NHS ester (section 1.3) was added drop-wise at and the reaction mixture was allowed to stir at r.t. Upon complete conversion according to LCMS (usually < 30 min), 5 vol% of piperdine was added to the reaction mixture and the reaction allowed to stir for additional 10 min, before it was quenched by addition of 5 vol% HOAc and 10 vol% water and subjected to RP-HPLC.

##### 1.6. General procedure for peptide couplings for branching

A 1 mL vial was charged with 3.0 equiv. amine dissolved in DMF (1 mL / 10 mg) and 8.0 equiv. DIPEA. The bis NHS ester **4** (1.0 equiv.) was dissolved in the same amount of DMF and added slowly and dropwise under vigorous stirring. The order and speed of addition is crucial to afford minimal amounts of side-products (*i.e.* imids, mono amides of succinates). Upon complete conversion according to LCMS, the reaction was directly deprotected or quenched and subjected to RP-HPLC (see below).

##### 1.7. General procedure for Boc deprotection

A 15 mL falcon tube was charged with Boc protected compound and put in an ice bath. Pre-cooled (4 °C) TFA was added neat. The reaction mixture was vortexed to ensure homogeneity and put back on ice for 15 min before all volatiles were removed under a gentle stream of nitrogen. The residue was taken up in DMF/water (9/1) and subjected to RP-HPLC.

NOTE: Azobenzene containing reaction mixtures turned deep red upon addition of TFA.

##### 1.8. 5-((2-(2-((6-Chlorohexyl)oxy)ethoxy)ethyl)amino)-5-oxopentanoic acid (Halo-COOH)

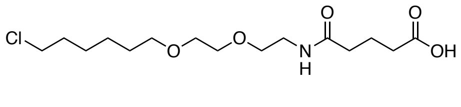

A 4 mL dram vial was charged with 100 mg (310 µmol, 1.0 equiv.) HaloNHBoc and 1 mL neat TFA was added. The solution was allowed to stand at r.t. for 5 min before all volatiles were removed under a gentle stream of nitrogen. 1 mL DMF and 160 µLDIPEA were added, before 35.3 mg (310 µmol, 1.0 equiv.) of glutaric anhydride was added in one portion. The reaction mixture was incubated o.n., before it was quenched with 160 µL HOAc, diluted with water and subjected to RP-HPLC to obtain 92 mg (274 µmol) of the desired product as a clear oil after lyophilization in 88% yield.

**HRMS** (ESI): calc. for $C_{15}H_{29}ClNO_5$ $[M+H]^+$: 338.1729, found: 338.1728.

##### 1.9. Bis(2,5-dioxopyrrolidin-1-yl) (((9*H*-fluoren-9-yl)methoxy)carbonyl)-*L*-glutamate (4)

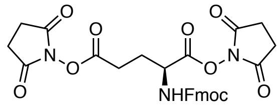

A 4 mL dram vial was charged with 300 mg (812 µmol, 1.0 equiv.) of (((9H-fluoren-9-yl)methoxy)carbonyl-*L*-glutamic acid (**3**) dissolved in 3 mL DMSO and 850 µL DIPEA before 978 mg (3.25 mmol, 4.0 equiv.) TSTU was added in one portion. The reaction mixture was stirred vigorously for 1 h before it was quenched by the addition of 850 µL HOAc and 200 µL water and subjected to preparative RP-HPLC. The desired product was obtained as a white powder after lyophilization in 35% yield (161 mg, 286 µmol).

NOTE: Immediate freeze-drying after elution from the HPLC system is highly recommended to suppress hydrolysis. The product was aliquoted to avoid multiple freeze-thaw cycles that also hydrolyzed the NHS esters.

**HRMS** (ESI): calc. for $C_{28}H_{26}N_3O_{10}$ $[M+H]^+$: 564.1613, found: 564.1614.

**1.10. (2*S*,2'*S*,4*S*,4'*S*)-4,4'-(((((1*E*,1'*E*)-(((2,2'-(((*S*)-2-Aminopentanedioyl)bis(azanediyl))bis(acetyl))bis(azanediyl))bis(4,1-phenylene))bis(diazene-2,1-diyl))bis(4,1-phenylene))bis(azanediyl))bis(4-oxobutane-4,1-diyl))bis(2-((*tert*-butoxycarbonyl)amino)pentanedioic acid) (5)**

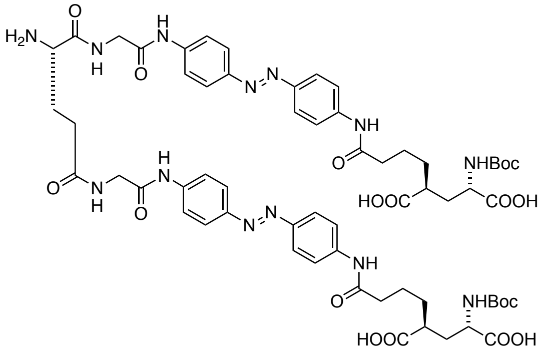

**5** was prepared according to general procedure 1.5 and was in situ deprotected by addition of 5 vol% of piperdine to the reaction mixture. The reaction allowed to stir for additional 10 min, before it was quenched by addition of 5 vol% HOAc and 10 vol% water and subjected to RP-HPLC.

**HRMS** (ESI): calc. for $C_{61}H_{79}N_{13}O_{18}$ $[M+2H]^{2+}$: 640.7828, found: 640.7835.

**1.11. (2*S*,4*S*)-2-(4-((4-((*E*)-(4-((*S*)-1-amino-41-((2-((4-((*E*)-(4-((5*S*,7*S*)-7-((*tert*-Butoxycarbonyl)amino)-5,7-dicarboxyheptanamido)phenyl)diazenyl)phenyl)amino)-2-oxoethyl)carbamoyl)-39,44-dioxo-3,6,9,12,15,18,21,24,27,30,33,36-dodecaoxa-40,45-diazaheptatetracontan-47-amido)phenyl)diazenyl)phenyl)amino)-4-oxobutyl)-4-((*tert*-butoxycarbonyl)amino)pentanedioic acid (6)**

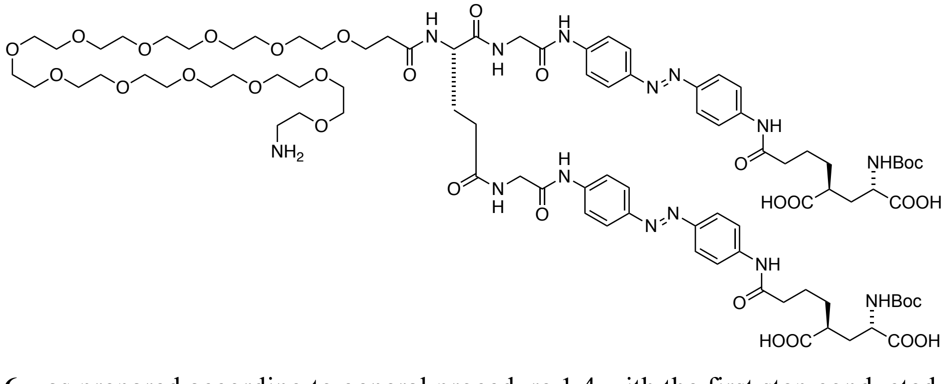

**6** was prepared according to general procedure 1.4 with the first step conducted at 50 °C.

**HRMS** (ESI): calc. for $C_{88}H_{132}N_{14}O_{31}$ $[M+2H]^{2+}$: 940.4586, found: 940.4582.

**1.12. (2*S*,4*S*)-2-(4-((4-((*E*)-(4-((*S*)-1-(4-(((2-Amino-9*H*-purin-6-yl)oxy)methyl)phenyl)-49-((2-((4-((*E*)-(4-((5*S*,7*S*)-7-((*tert*-butoxycarbonyl)amino)-5,7-dicarboxyheptanamido)phenyl)diazenyl)phenyl)amino)-2-oxoethyl)carbamoyl)-3,7,47,52-tetraoxo-11,14,17,20,23,26,29,32,35,38,41,44-dodecaoxa-2,8,48,53-tetraazapentapentacontan-55-amido)phenyl)diazenyl)phenyl)amino)-4-oxobutyl)-4-((*tert*-butoxycarbonyl)amino)pentanedioic acid (7)**

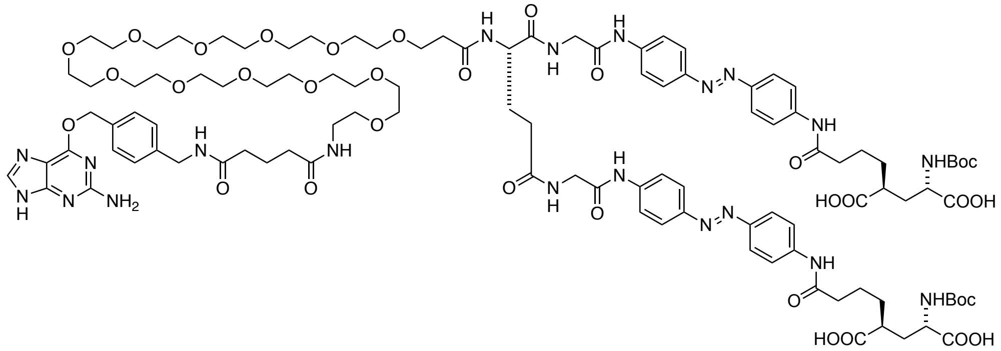

**7** was prepared according to general procedure 1.4, with **BG-COOSu** prepared according to general procedure 1.3, without the addition of piperidine.

**HRMS** (ESI): calc. for $C_{106}H_{150}N_{20}O_{34}$ $[M+2H]^{2+}$: 1124.0321, found: 1124.0327.

**1.13. (2*S*,4*S*)-2-Amino-4-(4-((4-((*E*)-(4-((*S*)-49-(3-((2-((4-((*E*)-(4-((5*S*,7*S*)-7-amino-5,7-dicarboxyheptanamido)phenyl)diazenyl)phenyl)amino)-2-oxoethyl)amino)-3-oxopropyl)-1-(4-(((2-amino-9*H*-purin-6-yl)oxy)methyl)phenyl)-3,7,47,50-tetraoxo-11,14,17,20,23,26,29,32,35,38,41,44-dodecaoxa-2,8,48,51-tetraazatripentacontan-53-amido)phenyl)diazenyl)phenyl)amino)-4-oxobutyl)pentanedioic acid ($dBGAG_{12}$)**

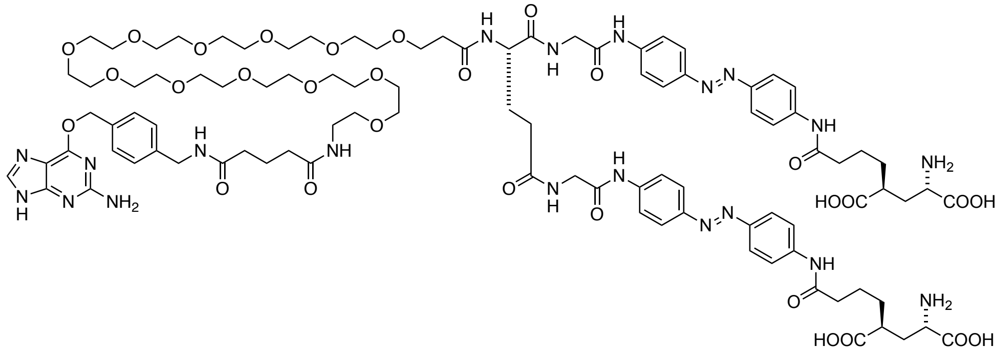

**$dBGAG_{12}$** was prepared according to general procedure 1.6.

**HRMS** (ESI): calc. for $C_{96}H_{134}N_{20}O_{30}$ $[M+2H]^{2+}$: 1032.9797, found: 1032.9787.

**1.14. (2*S*,2'*S*,4*S*,4'*S*)-4,4'-(((((1*E*,1'*E*)-(((2,2'-(((*S*)-2-(5-((4-(((2-Amino-9*H*-purin-6-yl)oxy)methyl)benzyl)amino)-5-oxopentanamido)pentanedioyl)bis(azanediyl))bis(acetyl))bis(azanediyl))bis(4,1-phenylene))bis(diazene-2,1-diyl))bis(4,1-phenylene))bis(azanediyl))bis(4-oxobutane-4,1-diyl))bis(2-((*tert*-butoxycarbonyl)amino)pentanedioic acid) (8)**

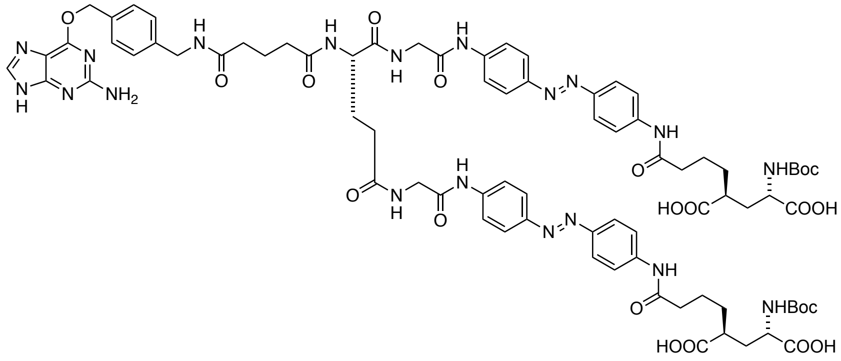

**8** was prepared according to general procedure 1.4, with **BG-COOSu** prepared according to general procedure 1.3, with the first step conducted at 50 °C and without the addition of piperidine.

**HRMS** (ESI): calc. for $C_{79}H_{95}N_{19}O_{21}$ $[M–H]^{2–}$: 821.8402, found: 821.8413.

**1.15. (2*S*,2'*S*,4*S*,4'*S*)-4,4'-(((((1*E*,1'*E*)-(((2,2'-(((*S*)-2-(5-((4-(((2-Amino-9*H*-purin-6-yl)oxy)methyl)benzyl)amino)-5-oxopentanamido)pentanedioyl)bis(azanediyl))bis(acetyl))bis(azanediyl))bis(4,1-phenylene))bis(diazene-2,1-diyl))bis(4,1-phenylene))bis(azanediyl))bis(4-oxobutane-4,1-diyl))bis(2-aminopentanedioic acid) (dBGAG$_0$)**

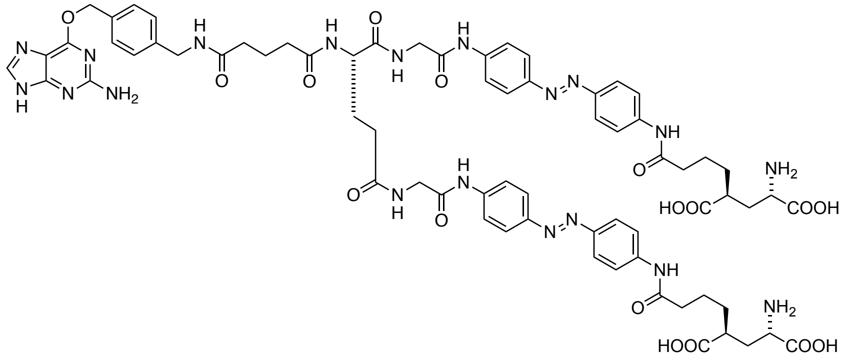

**dBGAG$_0$** was prepared according to general procedure 1.6.

**HRMS** (ESI): calc. for $C_{69}H_{79}N_{19}O_{17}$ $[M–H]^{2–}$: 721.7878, found: 721.7870.

**1.16. (2*S*,4*S*)-2-(4-((4-((*E*)-(4-(1-Amino-39-oxo-3,6,9,12,15,18,21,24,27,30,33,36-dodecaoxa-40-azadotetracontan-42-amido)phenyl)diazenyl)phenyl)amino)-4-oxobutyl)-4-((*tert*-butoxycarbonyl)amino)pentanedioic acid (9)**

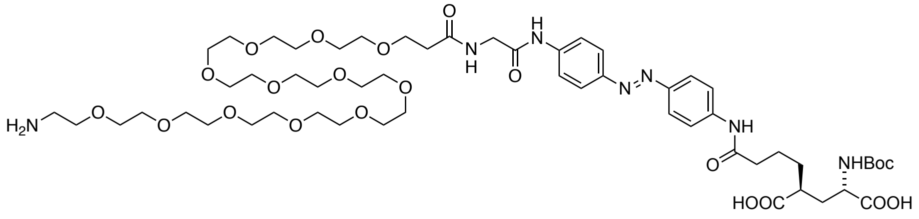

**9** was prepared according to general procedure 1.4.

**HRMS** (ESI): calc. for $C_{55}H_{90}N_7O_{21}$ $[M+H]^+$: 1184.6184, found: 1184.6176.

**1.17. (2*S*,2'*S*,4*S*,4'*S*)-4,4'-(((((1*E*,1'*E*)-((((*S*)-45-Amino-4,44,48,88-tetraoxo-7,10,13,16,19,22,25,28,31,34,37,40,52,55,58,61,64,67,70,73,76,79,82,85-tetracosaoxa-3,43,49,89-tetraazahennonacontanedioyl)bis(azanediyl))bis(4,1-phenylene))bis(diazene-2,1-diyl))bis(4,1-phenylene))bis(azanediyl))bis(4-oxobutane-4,1-diyl))bis(2-((*tert*-butoxycarbonyl)amino)pentanedioic acid) (10)**

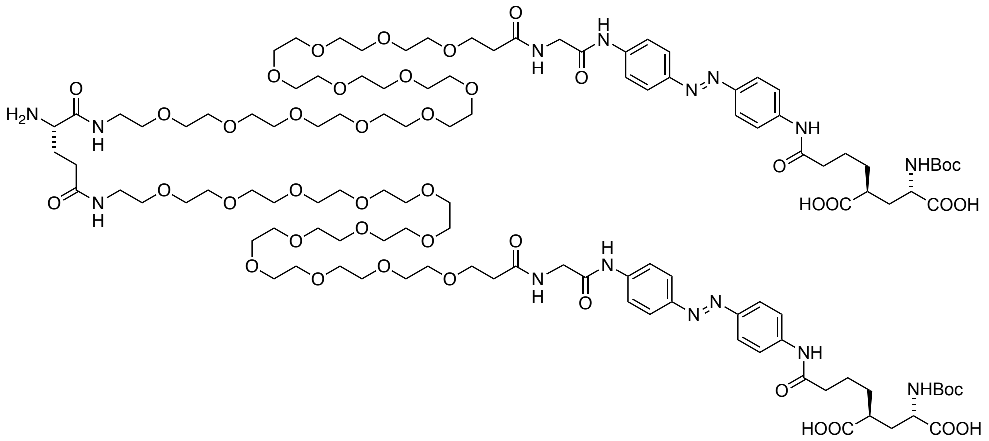

**10** was prepared according to general procedure 1.5 and was in situ deprotected by addition of 5 vol% of piperdine to the reaction mixture. The reaction allowed to stir for additional 10 min, before it was quenched by addition of 5 vol% HOAc and 10 vol% water and subjected to RP-HPLC.

**HRMS** (ESI): calc. for $C_{115}H_{186}N_{15}O_{44}$ $[M+3H]^{3+}$: 827.4264, found: 824.4261.

**1.18. (2*S*,2'*S*,4*S*,4'*S*)-4,4'-(((((1*E*,1'*E*)-((((*S*)-45-(5-((4-(((2-Amino-9*H*-purin-6-yl)oxy)methyl)benzyl)amino)-5-oxopentanamido)-4,44,48,88-tetraoxo-7,10,13,16,19,22,25,28,31,34,37,40,52,55,58,61,64,67,70,73,76,79,82,85-tetracosaoxa-3,43,49,89-tetraazahennonacontanedioyl)bis(azanediyl))bis(4,1-phenylene))bis(diazene-2,1-diyl))bis(4,1-phenylene))bis(azanediyl))bis(4-oxobutane-4,1-diyl))bis(2-((*tert*-butoxycarbonyl)amino)pentanedioic acid) (11)**

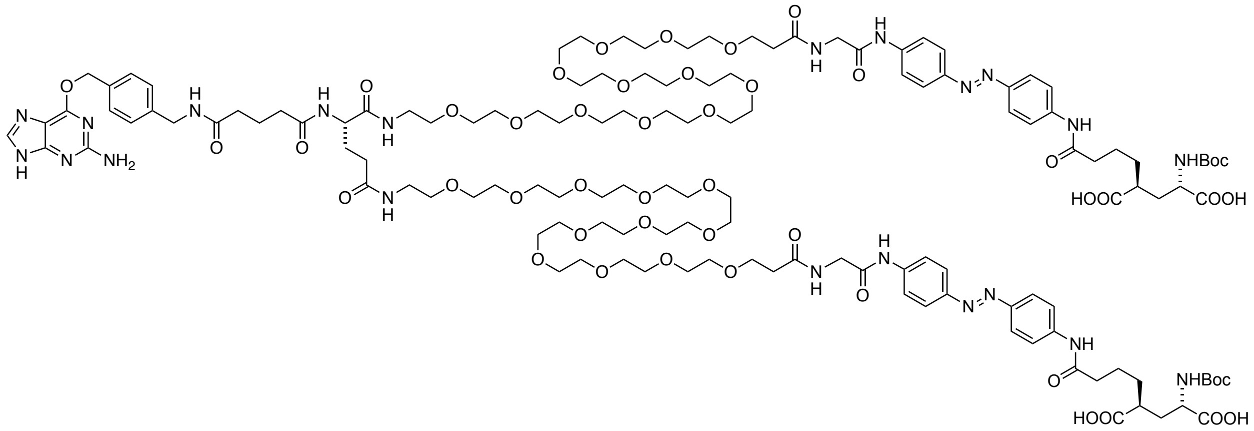

**11** was prepared according to general procedure 1.4, with **BG-COOSu** prepared according to general procedure 1.3, with the first step conducted at 50 °C and without the addition of piperidine.

**HRMS** (ESI): calc. for $C_{133}H_{204}N_{21}O_{47}$ $[M+3H]^{3+}$: 949.4744, found: 949.4747.

**1.19. (2*S*,2'*S*,4*S*,4'*S*)-4,4'-(((((1*E*,1'*E*)-((((*S*)-45-(5-((4-(((2-Amino-9*H*-purin-6-yl)oxy)methyl)benzyl)amino)-5-oxopentanamido)-4,44,48,88-tetraoxo-7,10,13,16,19,22,25,28,31,34,37,40,52,55,58,61,64,67,70,73,76,79,82,85-tetracosaoxa-3,43,49,89-tetraazahennonacontanedioyl)bis(azanediyl))bis(4,1-phenylene))bis(diazene-2,1-diyl))bis(4,1-phenylene))bis(azanediyl))bis(4-oxobutane-4,1-diyl))bis(2-aminopentanedioic acid) (dBGAG$_{12}$v2)**

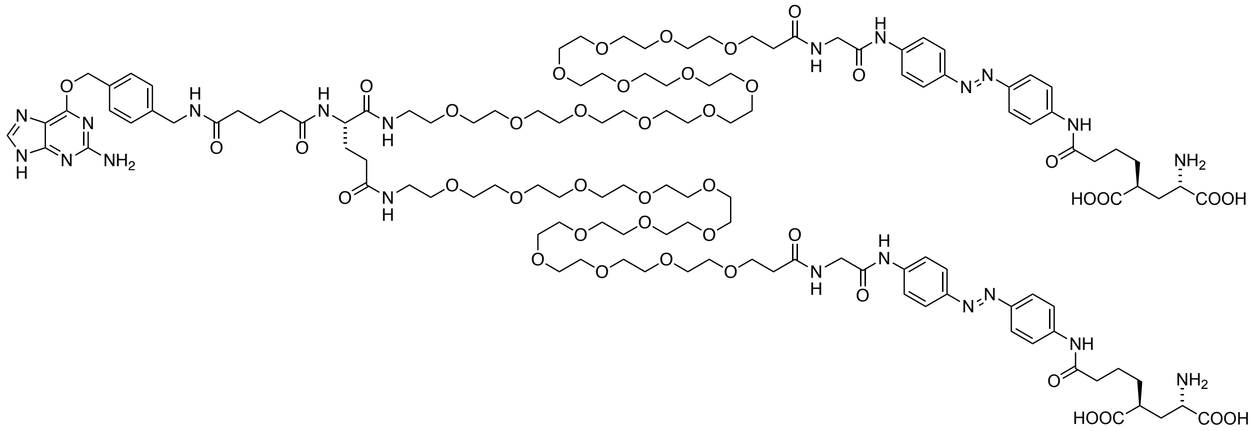

**dBGAG$_{12}$v2** was prepared according to general procedure 1.6.

**HRMS** (ESI): calc. for $C_{123}H_{188}N_{21}O_{43}$ $[M+3H]^{3+}$: 882.7728, found: 882.7723.

**1.20. (2*S*,2'*S*,4*S*,4'*S*)-4,4'-(((((1*E*,1'*E*)-((((*S*)-45-amino-4,44,48,88-tetraoxo-7,10,13,16,19,22,25,28,31,34,37,40,52,55,58,61,64,67,70,73,76,79,82,85-tetracosaoxa-3,43,49,89-tetraazahennonacontanedioyl)bis(azanediyl))bis(4,1-phenylene))bis(diazene-2,1-diyl))bis(4,1-phenylene))bis(azanediyl))bis(4-oxobutane-4,1-diyl))bis(2-((*tert*-butoxycarbonyl)amino)pentanedioic acid) (12)**

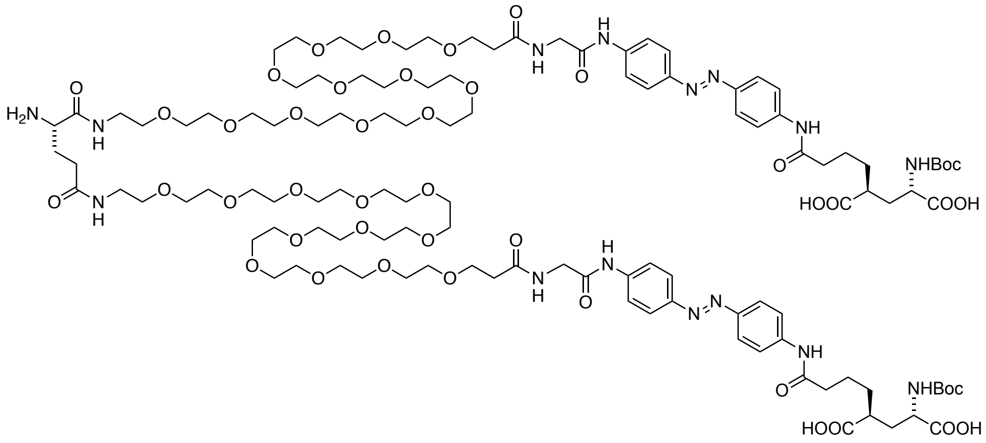

**12** was prepared according to general procedure 1.5 and was in situ deprotected by addition of 5 vol% of piperdine to the reaction mixture. The reaction allowed to stir for additional 10 min, before it was quenched by addition of 5 vol% HOAc and 10 vol% water and subjected to RP-HPLC.

**HRMS** (ESI): calc. for $C_{115}H_{186}N_{15}O_{44}$ $[M+3H]^{3+}$: 827.4264, found: 827.4255.

**1.21. (2*S*,4*S*)-2-[3-({4-[(1*E*)-2-[4-(2-{1-[(4*S*)-4-[(2*S*)-2-Amino-4-{[(1*S*)-1,3-bis[(38-{[({4-[(1*E*)-2-{4-[(5*S*,7*S*)-7-{[(*tert*-butoxy)carbonyl]amino}-5,7-dicarboxyheptanamido]phenyl}diazen-1-yl]phenyl}carbamoyl)methyl]carbamoyl}-3,6,9,12,15,18,21,24,27,30,33,36-dodecaoxaoctatriacontan-1-yl)carbamoyl]propyl]carbamoyl}butanamido]-4-[(38-{[({4-[(1*E*)-2-{4-[(5*S*,7*S*)-7-{[(*tert*-butoxy)carbonyl]amino}-5,7-dicarboxyheptanamido]phenyl}diazen-1-yl]phenyl}carbamoyl)methyl]carbamoyl}-3,6,9,12,15,18,21,24,27,30,33,36-dodecaoxaoctatriacontan-1-yl)carbamoyl]butanamido]-3,6,9,12,15,18,21,24,27,30,33,36-dodecaoxanonatriacontan-39-amido}acetamido)phenyl]diazen-1-yl]phenyl}carbamoyl)propyl]-4-{[(*tert*-butoxy)carbonyl]amino}pentanedioic acid (13)**

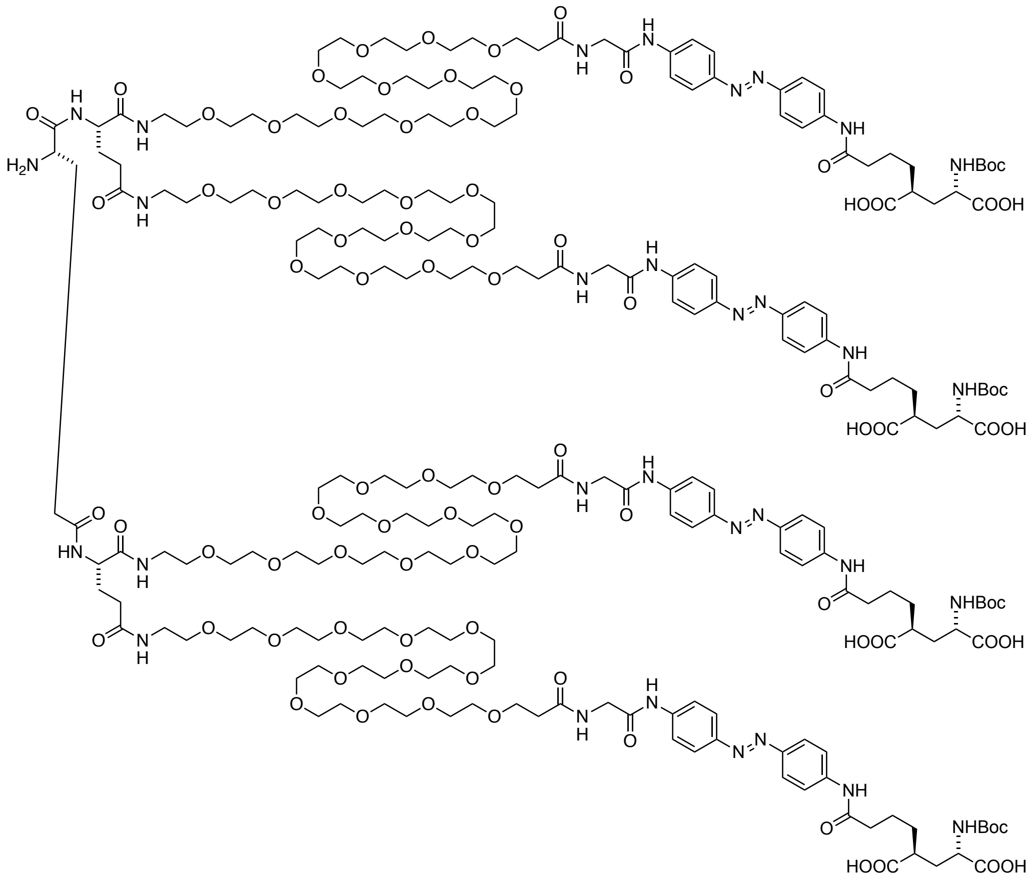

**13** was prepared according to general procedure 1.4 with the first step conducted at 50 °C.

**HRMS** (ESI): calc. for $C_{235}H_{375}N_{31}O_{90}$ $[M+4H]^{4+}$:1286.3940, found:1286.3925.

**1.22. (2*S*,4*S*)-2-[3-({4-[(1*E*)-2-[4-(2-{1-[(4*S*)-4-[(2*S*)-2-(4-{[(4-{[(2-Amino-9*H*-purin-6-yl)oxy]methyl}phenyl)methyl]carbamoyl}butanamido)-4-{[(1S)-1,3-bis[(38-{[({4-[(1*E*)-2-{4-[(5*S*,7*S*)-7-{[(*tert*-butoxy)carbonyl]amino}-5,7-dicarboxyheptanamido]phenyl}diazen-1-yl]phenyl}carbamoyl)methyl]carbamoyl}-3,6,9,12,15,18,21,24,27,30,33,36-dodecaoxaoctatriacontan-1-yl)carbamoyl]propyl]carbamoyl}butanamido]-4-[(38-{[({4-[(1*E*)-2-{4-[(5*S*,7*S*)-7-{[(tert-butoxy)carbonyl]amino}-5,7-dicarboxyheptanamido]phenyl}diazen-1-yl]phenyl}carbamoyl)methyl]carbamoyl}-3,6,9,12,15,18,21,24,27,30,33,36-dodecaoxaoctatriacontan-1-yl)carbamoyl]butanamido]-3,6,9,12,15,18,21,24,27,30,33,36-dodecaoxanonatriacontan-39-amido}acetamido)phenyl]diazen-1-yl]phenyl}carbamoyl)propyl]-4-{[(*tert*-butoxy)carbonyl]amino}pentanedioic acid (14)**

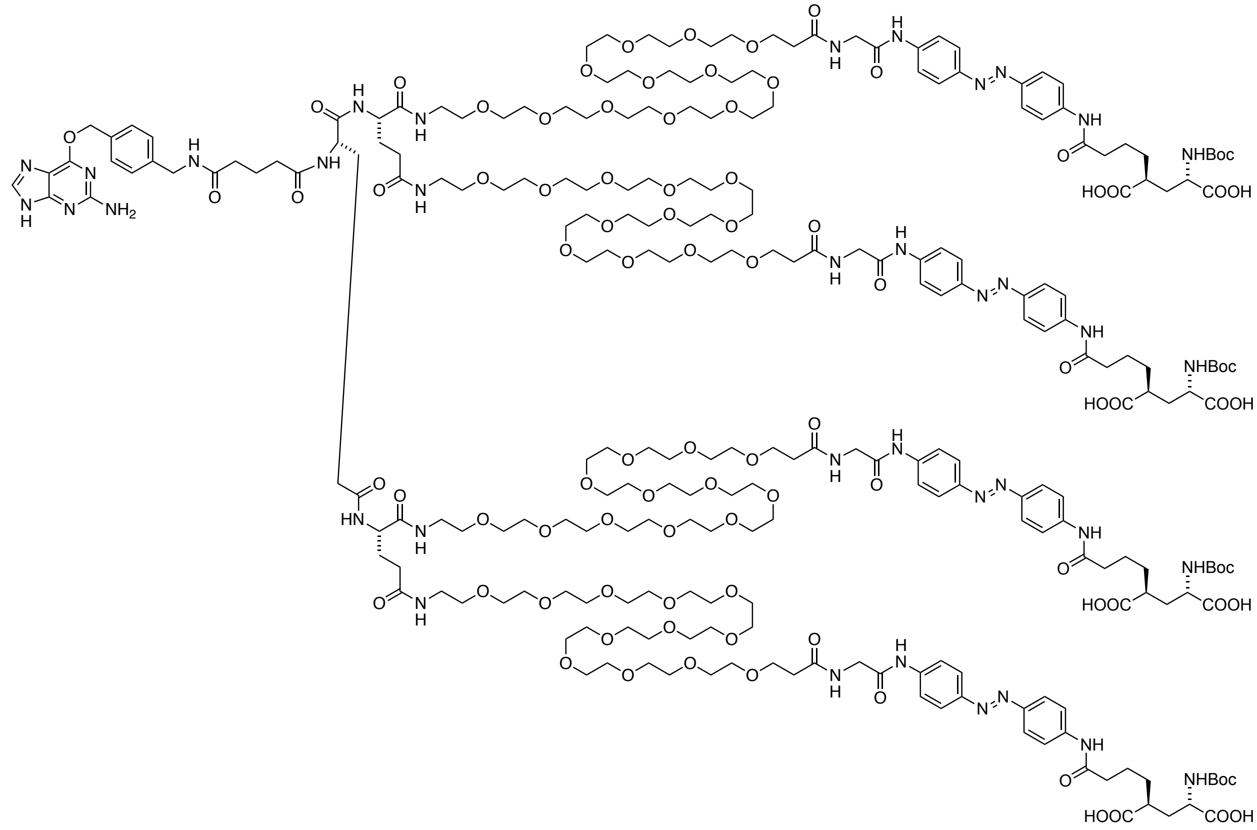

**14** was prepared according to general procedure 1.4, with **BG-COOSu** prepared according to general procedure 1.3, without the addition of piperidine.

**HRMS** (ESI): calc. for $C_{253}H_{1385}N_{37}O_{93}$ $[M+4H]^{4+}$: 1360.1807, found: 1360.1804.

**1.23. (2*S*,4*S*)-2-Amino-4-[3-({4-[(1*E*)-2-[4-(2-{1-[(4*S*)-4-[(38-{[({4-[(1*E*)-2-{4-[(5*S*,7*S*)-7-amino-5,7-dicarboxyheptanamido]phenyl}diazen-1-yl]phenyl}carbamoyl)methyl]carbamoyl}-3,6,9,12,15,18,21,24,27,30,33,36-dodecaoxaoctatriacontan-1-yl)carbamoyl]-4-[(2*S*)-2-(4-{[(4-{[(2-amino-9*H*-purin-6-yl)oxy]methyl}phenyl)methyl]carbamoyl}butanamido)-4-{[(1*S*)-1,3-bis[(38-{[({4-[(1*E*)-2-{4-[(5*S*,7*S*)-7-amino-5,7-dicarboxyheptanamido]phenyl}diazen-1-yl]phenyl}carbamoyl)methyl]carbamoyl}-3,6,9,12,15,18,21,24,27,30,33,36-dodecaoxaoctatriacontan-1-yl)carbamoyl]propyl]carbamoyl}butanamido]butanamido]-3,6,9,12,15,18,21,24,27,30,33,36-dodecaoxanonatriacontan-39-amido}acetamido)phenyl]diazen-1-yl]phenyl}carbamoyl)propyl]pentanedioic acid (qBGAG)**

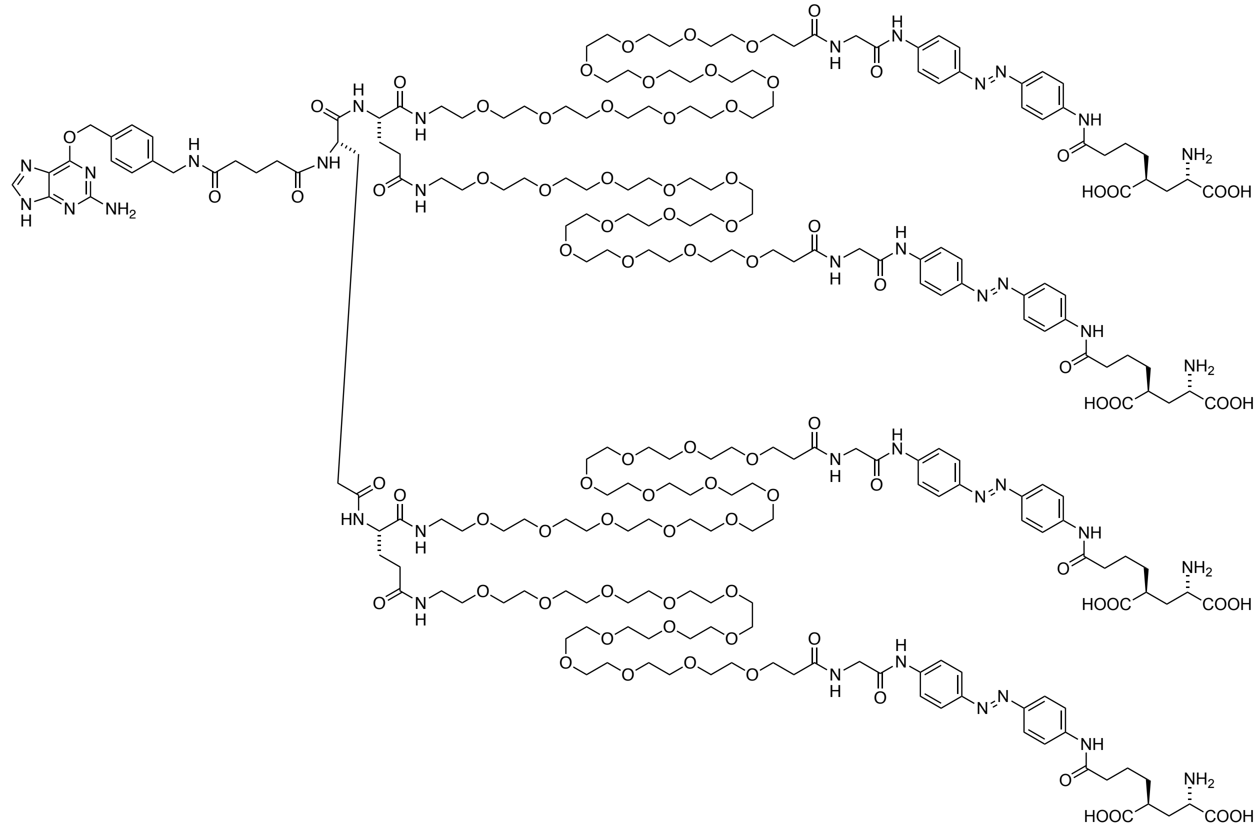

**qBGAG** was prepared according to general procedure 1.6.

**HRMS** (ESI): calc. for $C_{233}H_{363}N_{37}O_{85}$ $[M+6H]^{6+}$: 840.2541, found: 840.2533.

**1.24. (2*S*,4*S*)-2-(4-((4-((*E*)-(4-((*S*)-1-Azido-41-((2-((4-((*E*)-(4-((5*S*,7*S*)-7-((*tert*-butoxycarbonyl)amino)-5,7-dicarboxyheptanamido)phenyl)diazenyl)phenyl)amino)-2-oxoethyl)carbamoyl)-39,44-dioxo-3,6,9,12,15,18,21,24,27,30,33,36-dodecaoxa-40,45-diazaheptatetracontan-47-amido)phenyl)diazenyl)phenyl)amino)-4-oxobutyl)-4-((*tert*-butoxycarbonyl)amino)pentanedioic acid (15)**

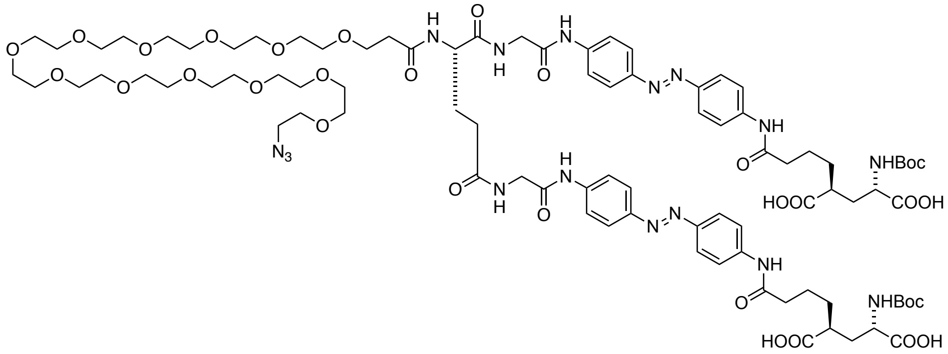

**15** was prepared according to general procedure 1.4 without adding piperidine for deprotection. The crude was subjected to RP-HPLC and azobenzene containing fractions were collected, dried and subjected to the next step without further characterization.

**1.25. (2*S*,4*S*)-2-Amino-4-(4-((4-((*E*)-(4-((*S*)-41-(3-((2-((4-((*E*)-(4-((5*S*,7*S*)-7-amino-5,7-dicarboxyheptanamido)phenyl)diazenyl)phenyl)amino)-2-oxoethyl)amino)-3-oxopropyl)-1-azido-39,42-dioxo-3,6,9,12,15,18,21,24,27,30,33,36-dodecaoxa-40,43-diazapentatetracontan-45-amido)phenyl)diazenyl)phenyl)amino)-4-oxobutyl)pentanedioic acid (16)**

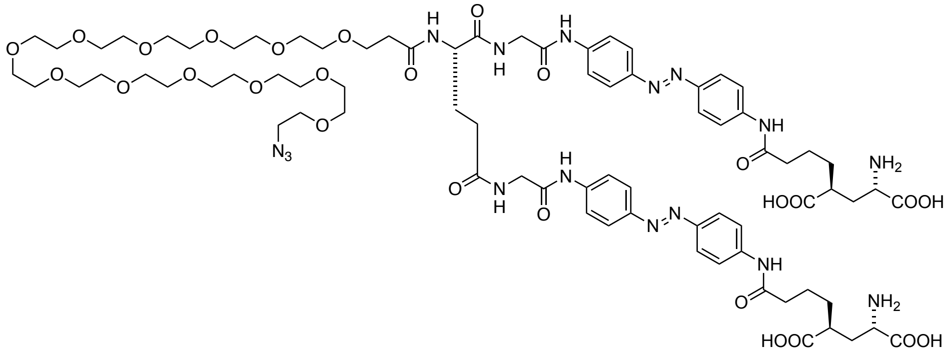

**16** was prepared according to general procedure 1.6.

**HRMS** (ESI): calc. for $C_{78}H_{115}N_{16}O_{27}$ $[M+3H]^{3+}$: 569.2700, found: 569.2705.

**1.26. (2*S*,4*S*)-2-Amino-4-(4-((4-((*E*)-(4-((*S*)-41-(3-((2-((4-((*E*)-(4-((5*S*,7*S*)-7-amino-5,7-dicarboxyheptanamido)phenyl)diazenyl)phenyl)amino)-2-oxoethyl)amino)-3-oxopropyl)-1-(8-(4-((4-(((4-aminopyrimidin-2-yl)oxy)methyl)benzyl)amino)-4-oxobutanoyl)-8,9-dihydro-3*H*-dibenzo[*b*,*f*][1,2,3]triazolo[4,5-*d*]azocin-3-yl)-39,42-dioxo-3,6,9,12,15,18,21,24,27,30,33,36-dodecaoxa-40,43-diazapentatetracontan-45-amido)phenyl)diazenyl)phenyl)amino)-4-oxobutyl)pentanedioic acid (dBCAG$_{12}$)**

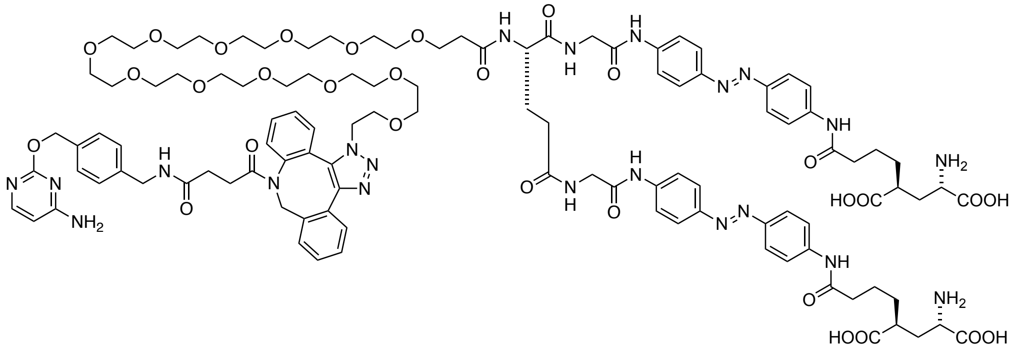

A 4 mL dram vial was charged with **16** dissolved in MeOH. BC-DBCO was added in one portion and the reaction mixture was incubated o.n. before all volatiles were removed under a gentle stream of nitrogen. The crude was taken up in DMF and water (9/1) and subjected to RP-HPLC purification to obtain **dBCAG12** as a yellow powder after lyophilization.

**HRMS** (ESI): calc. for $C_{109}H_{142}N_{21}O_{30}$ $[M+3H]^{3+}$: 742.0082, found: 742.0080.

**1.27. (2*S*,4*S*)-2-((*tert*-Butoxycarbonyl)amino)-4-(4-((4-((*E*)-(4-(61-chloro-4,44,48-trioxo-7,10,13,16,19,22,25,28,31,34,37,40,52,55-tetradecaoxa-3,43,49-triazahenhexacontanamido)phenyl)diazenyl)phenyl)amino)-4-oxobutyl)pentanedioic acid (17)**

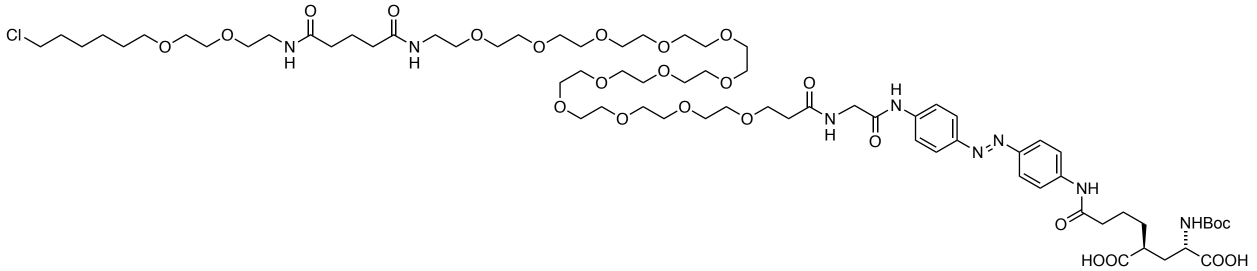

**17** was prepared according to general procedure 1.4, with **Halo-COOSu** prepared according to general procedure 1.3, without the addition of piperidine.

**HRMS** (ESI): calc. for $C_{70}H_{117}ClN_8O_{25}$ $[M+2H]^{2+}$: 753.3913, found: 753.3917.

**1.28. (2*S*,4*S*)-2-Amino-4-(4-((4-((*E*)-(4-(61-chloro-4,44,48-trioxo-7,10,13,16,19,22,25,28,31,34,37,40,52,55-tetradecaoxa-3,43,49-triazahenhexacontanamido)phenyl)diazenyl)phenyl)amino)-4-oxobutyl)pentanedioic acid (ClAG$_{12}$)**

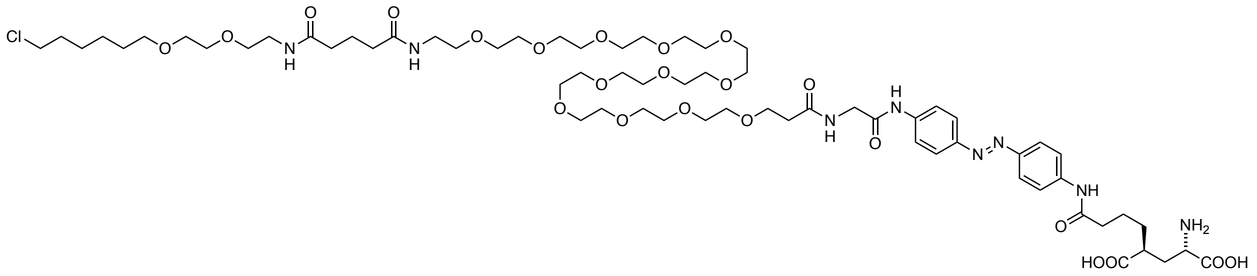

**ClAG$_{12}$** was prepared according to general procedure 1.6.

**HRMS** (ESI): calc. for $C_{65}H_{109}ClN_8O_{23}$ $[M+2H]^{2+}$: 702.3642, found: 702.3642.

**1.29. (2*S*,4*S*)-2-((*tert*-Butoxycarbonyl)amino)-4-(4-((4-((*E*)-(4-((*S*)-5-(3-((2-((4-((*E*)-(4-((5*S*,7*S*)-7-((*tert*-butoxycarbonyl)amino)-5,7-dicarboxyheptanamido)phenyl)diazenyl)phenyl)amino)-2-oxoethyl)amino)-3-oxopropyl)-64-chloro-4,7,47,51-tetraoxo-10,13,16,19,22,25,28,31,34,37,40,43,55,58-tetradecaoxa-3,6,46,52-tetraazatetrahexacontanamido)phenyl)diazenyl)phenyl)amino)-4-oxobutyl)pentanedioic acid (18)**

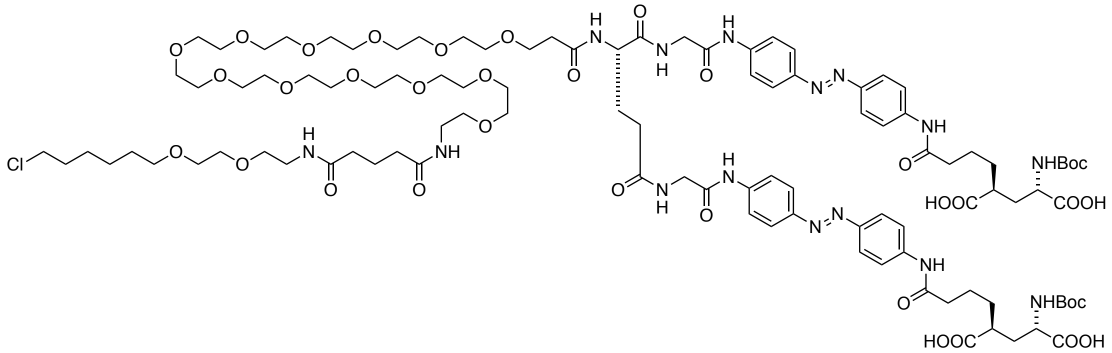

**18** was prepared according to general procedure 1.4, with **Halo-COOSu** prepared according to general procedure 1.3, without the addition of piperidine.

**HRMS** (ESI): calc. for $C_{103}H_{158}ClN_{15}O_{35}$ $[M+2H]^{2+}$: 1100.5377, found: 1100.5370.

**1.30. (2*S*,4*S*)-2-Amino-4-(4-((4-((*E*)-(4-((*S*)-5-(3-((2-((4-((*E*)-(4-((5*S*,7*S*)-7-amino-5,7-dicarboxyheptanamido)phenyl)diazenyl)phenyl)amino)-2-oxoethyl)amino)-3-oxopropyl)-64-chloro-4,7,47,51-tetraoxo-10,13,16,19,22,25,28,31,34,37,40,43,55,58-tetradecaoxa-3,6,46,52-tetraazatetrahexacontanamido)phenyl)diazenyl)phenyl)amino)-4-oxobutyl)pentanedioic acid ($dClAG_{12}$)**

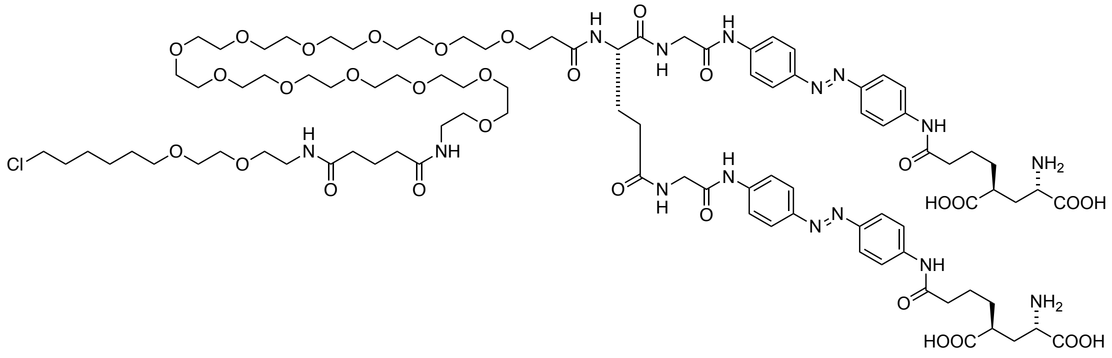

**$dClAG_{12}$** was prepared according to general procedure 1.6.

**HRMS** (ESI): calc. for $C_{93}H_{142}ClN_{15}O_{31}$ $[M+2H]^{2+}$: 1000.4852, found: 1000.4853.

**1.31. (2*S*,2'*S*,4*S*,4'*S*)-4,4'-(((((1*E*,1'*E*)-((((((*S*)-2-Aminopentanedioyl)bis(azanediyl))bis(ethane-2,1-diyl))bis(azanediyl))bis(4,1-phenylene))bis(diazene-2,1-diyl))bis(4,1-phenylene))bis(azanediyl))bis(4-oxobutane-4,1-diyl))bis(2-((*tert*-butoxycarbonyl)amino)pentanedioic acid) (19)**

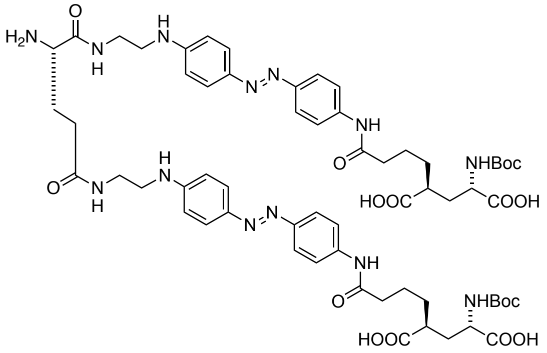

**19** was prepared according to general procedure 1.4 without the addition of piperidine. Instead, Fmoc-protected compound was obtained after RP-HPLC purification, dried and redissolved in MeCN with the addition of 5% DBU. The reaction mixture was incubated for 1 h, before it was quenched by addition of HOAc and water and subjected to RP-HPLC.

**HRMS** (ESI): calc. for $C_{61}H_{82}N_{13}O_{16}$ $[M+H]^{+}$: 1252.5997, found: 1252.5995.

**1.32. (2*S*,4*S*)-2-(4-((4-((*E*)-(4-(((*S*)-1-Amino-41-((2-((4-((*E*)-(4-((5*S*,7*S*)-7-((*tert*-butoxycarbonyl)amino)-5,7-dicarboxyheptanamido)phenyl)diazenyl)phenyl)amino)ethyl)carbamoyl)-39,44-dioxo-3,6,9,12,15,18,21,24,27,30,33,36-dodecaoxa-40,45-diazaheptatetracontan-47-yl)amino)phenyl)diazenyl)phenyl)amino)-4-oxobutyl)-4-((*tert*-butoxycarbonyl)amino)pentanedioic acid (20)**

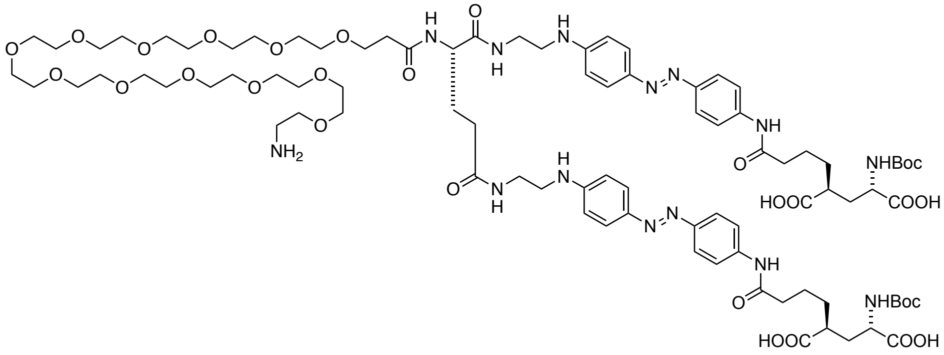

**20** was prepared according to general procedure 1.4 with the first step conducted at 50 °C and without the addition of piperidine. Instead, Fmoc-protected compound was obtained after RP-HPLC purification, dried and redissolved in MeCN with the addition of 5% DBU. The reaction mixture was incubated for 1 h, before it was quenched by addition of HOAc and water and subjected to RP-HPLC.

**HRMS** (ESI): calc. for $C_{88}H_{136}N_{14}O_{29}$ $[M+2H]^{2+}$: 926.9809, found: 926.9809.

**1.33. (2*S*,4*S*)-2-(4-((4-((*E*)-(4-(((*S*)-1-(4-(((2-Amino-9*H*-purin-6-yl)oxy)methyl)phenyl)-49-((2-((4-((*E*)-(4-((5*S*,7*S*)-7-((*tert*-butoxycarbonyl)amino)-5,7-dicarboxyheptanamido)phenyl)diazenyl)phenyl)amino)ethyl)carbamoyl)-3,7,47,52-tetraoxo-11,14,17,20,23,26,29,32,35,38,41,44-dodecaoxa-2,8,48,53-tetraazapentapentacontan-55-yl)amino)phenyl)diazenyl)phenyl)amino)-4-oxobutyl)-4-((*tert*-butoxycarbonyl)amino)pentanedioic acid (21)**

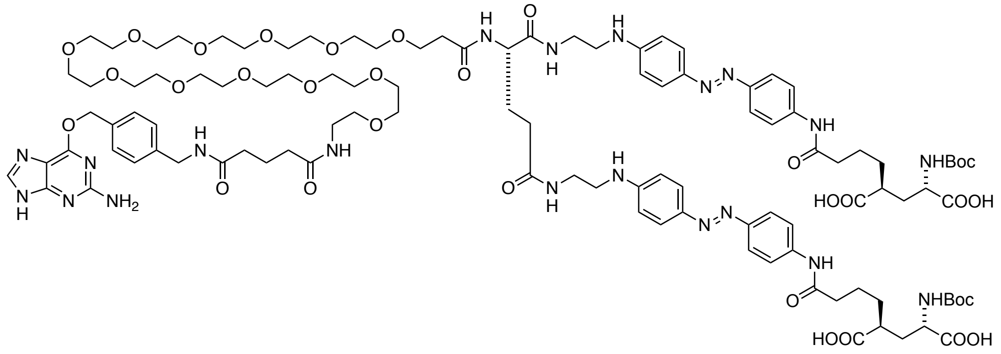

**21** was prepared according to general procedure 1.4, with **BG-COOSu** prepared according to general procedure 1.3, without the addition of piperidine.

**HRMS** (ESI): calc. for $C_{106}H_{154}N_{20}O_{32}$ $[M+2H]^{2+}$: 1110.0529, found:1110.0531.

**1.34. (2*S*,4*S*)-2-Amino-4-(4-((4-((*E*)-(4-(((*S*)-49-(3-((2-((4-((*E*)-(4-((5*S*,7*S*)-7-amino-5,7-dicarboxyheptanamido)phenyl)diazenyl)phenyl)amino)ethyl)amino)-3-oxopropyl)-1-(4-(((2-amino-9*H*-purin-6-yl)oxy)methyl)phenyl)-3,7,47,50-tetraoxo-11,14,17,20,23,26,29,32,35,38,41,44-dodecaoxa-2,8,48,51-tetraazatripentacontan-53-yl)amino)phenyl)diazenyl)phenyl)amino)-4-oxobutyl)pentanedioic acid (dBGAG$_{12,460}$)**

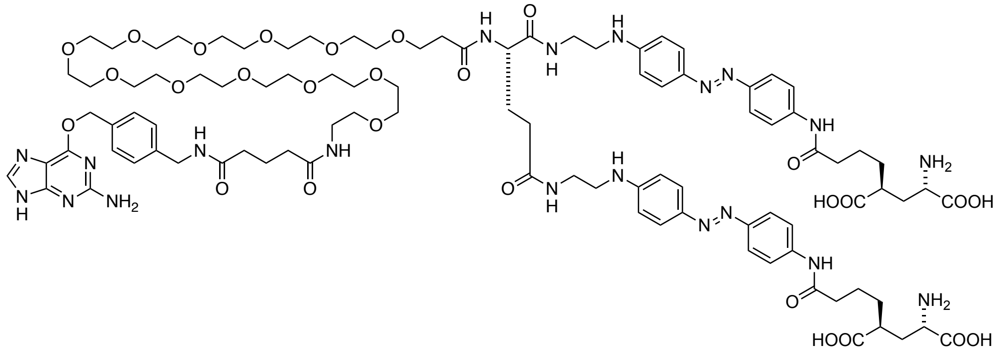

**dBGAG$_{12,460}$** was prepared according to general procedure 1.6.

**HRMS** (ESI): calc. for $C_{96}H_{140}N_{20}O_{28}$ $[M+4H]^{4+}$: 505.5038, found:505.5041.

**1.35. (2*S*,4*S*)-2-(4-((4-((*E*)-(4-((1-Amino-39-oxo-3,6,9,12,15,18,21,24,27,30,33,36-dodecaoxa-40-azadotetracontan-42-yl)amino)phenyl)diazenyl)phenyl)amino)-4-oxobutyl)-4-((*tert*-butoxycarbonyl)amino)pentanedioic acid (22)**

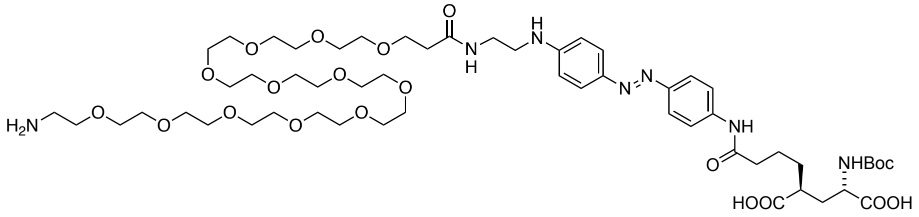

**22** was prepared according to general procedure 1.4 without the addition of piperidine. Instead, Fmoc-protected compound was obtained after RP-HPLC purification, dried and redissolved in MeCN with the addition of 5% DBU. The reaction mixture was incubated for 1 h, before it was quenched by addition of HOAc and water and subjected to RP-HPLC.

**HRMS** (ESI): calc. for $C_{55}H_{93}N_7O_{20}$ $[M+2H]^{2+}$: 585.8232, found: 585.8232.

**1.36. (2*S*,2'*S*,4*S*,4'*S*)-4,4'-(((((1*E*,1'*E*)-((((*S*)-45-Amino-4,44,48,88-tetraoxo-7,10,13,16,19,22,25,28,31,34,37,40,52,55,58,61,64,67,70,73,76,79,82,85-tetracosaoxa-3,43,49,89-tetraazahennonacontane-1,91-diyl)bis(azanediyl))bis(4,1-phenylene))bis(diazene-2,1-diyl))bis(4,1-phenylene))bis(azanediyl))bis(4-oxobutane-4,1-diyl))bis(2-((*tert*-butoxycarbonyl)amino)pentanedioic acid) (23)**

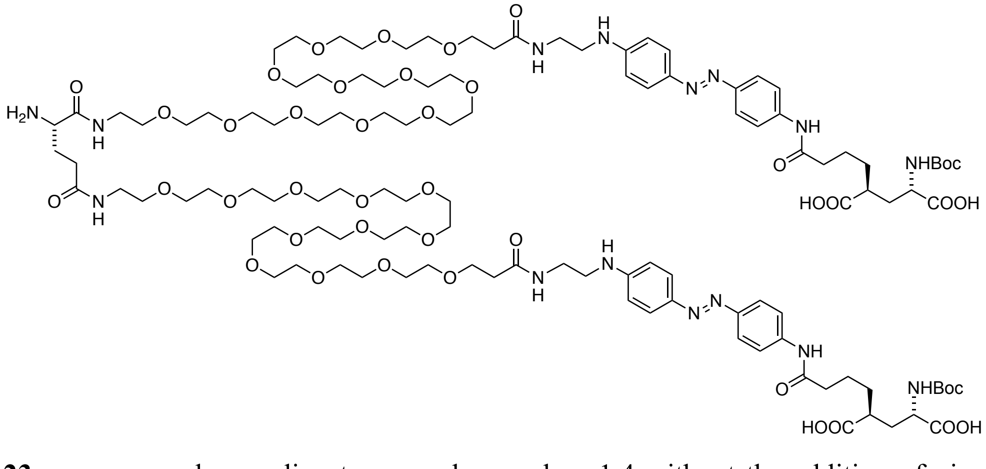

**23** was prepared according to general procedure 1.4 without the addition of piperidine. Instead, Fmoc-protected compound was obtained after RP-HPLC purification, dried and redissolved in MeCN with the addition of 5% DBU. The reaction mixture was incubated for 1 h, before it was quenched by addition of

**HRMS** (ESI): calc. for $C_{115}H_{190}N_{15}O_{42}$ $[M+3H]^{3+}$: 818.1069, found: 818.1075.

**1.37. (2*S*,2'*S*,4*S*,4'*S*)-4,4'-(((((1*E*,1'*E*)-((((*S*)-45-(5-((4-(((2-Amino-9*H*-purin-6-yl)oxy)methyl)benzyl)amino)-5-oxopentanamido)-4,44,48,88-tetraoxo-7,10,13,16,19,22,25,28,31,34,37,40,52,55,58,61,64,67,70,73,76,79,82,85-tetracosaoxa-3,43,49,89-tetraazahennonacontane-1,91-diyl)bis(azanediyl))bis(4,1-phenylene))bis(diazene-2,1-diyl))bis(4,1-phenylene))bis(azanediyl))bis(4-oxobutane-4,1-diyl))bis(2-((*tert*-butoxycarbonyl)amino)pentanedioic acid) (24)**

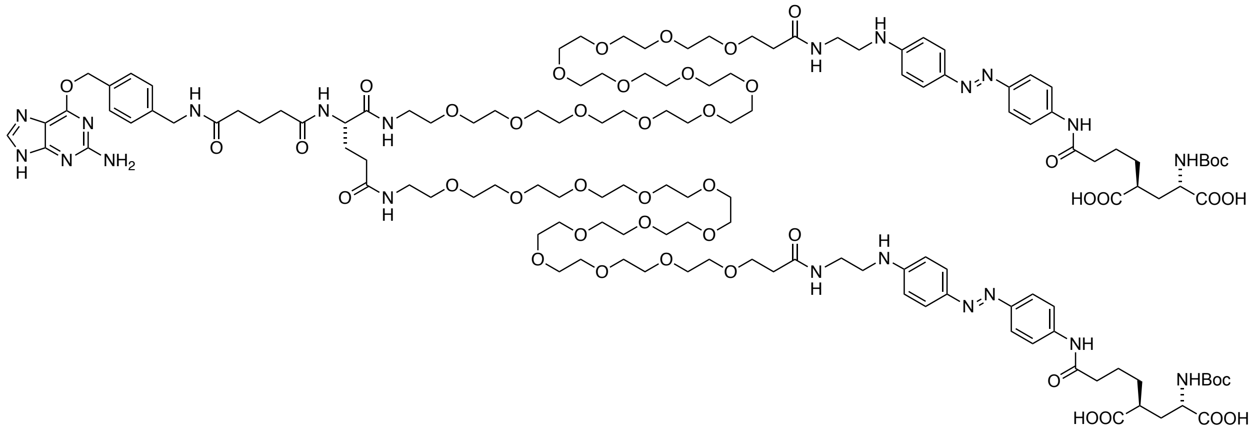

**24** was prepared according to general procedure 1.4, with **BG-COOSu** prepared according to general procedure 1.3, with the first step conducted at 50 °C and without the addition of piperidine.

**HRMS** (ESI): calc. for $C_{133}H_{208}N_{21}O_{45}$ $[M+3H]^{3+}$: 940.1549, found: 940.1546.

**1.38. (2*S*,2'*S*,4*S*,4'*S*)-4,4'-(((((1*E*,1'*E*)-((((*S*)-45-(5-((4-(((2-Amino-9*H*-purin-6-yl)oxy)methyl)benzyl)amino)-5-oxopentanamido)-4,44,48,88-tetraoxo-7,10,13,16,19,22,25,28,31,34,37,40,52,55,58,61,64,67,70,73,76,79,82,85-tetracosaoxa-3,43,49,89-tetraazahennonacontane-1,91-diyl)bis(azanediyl))bis(4,1-phenylene))bis(diazene-2,1-diyl))bis(4,1-phenylene))bis(azanediyl))bis(4-oxobutane-4,1-diyl))bis(2-aminopentanedioic acid) (dBGAG$_{12,460}$v2)**

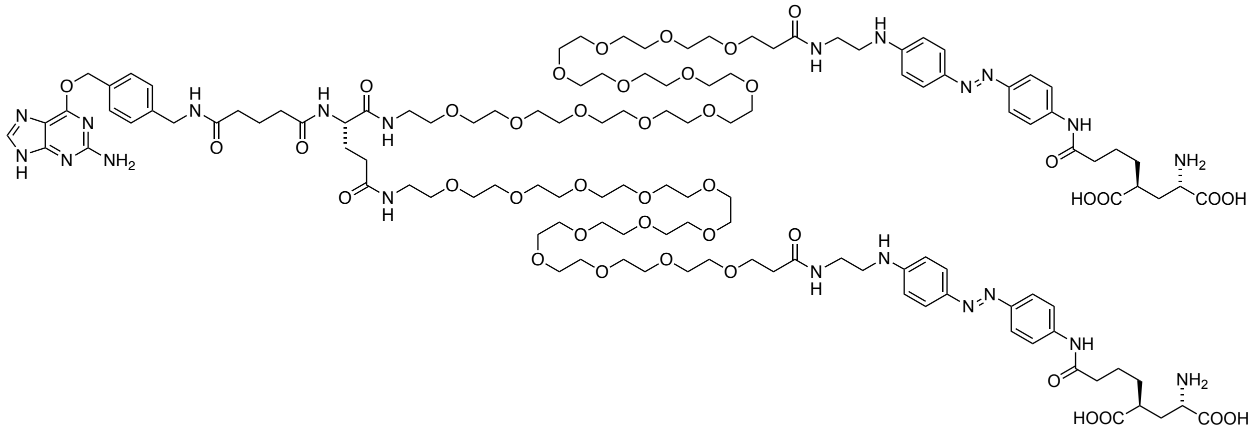

**dBGAG$_{12,460}$v2** was prepared according to general procedure 1.6.

**HRMS** (ESI): calc. for $C_{123}H_{193}N_{21}O_{41}$ $[M+4H]^{4+}$: 655.3418, found: 655.3416.

**1.39. 1-(6-(((*S*)-5-((((9*H*-Fluoren-9-yl)methoxy)carbonyl)amino)-5-carboxypentyl)amino)-6-oxohexyl)-3,3-dimethyl-2-((1*E*,3*E*)-5-((*Z*)-1,3,3-trimethylindolin-2-ylidene)penta-1,3-dien-1-yl)-3*H*-indol-1-ium (27)**

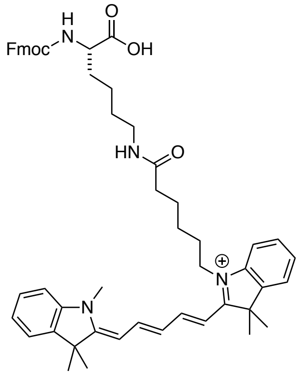

A round bottom flask was charged with 35.0 mg (72.4 µmol, 1.0 equiv.) of 1-(5-carboxypentyl)-3,3-dimethyl-2-((1*E*,3*E*)-5-((*Z*)-1,3,3-trimethylindolin-2-ylidene)penta-1,3-dien-1-yl)-3*H*-indol-1-ium (**26**)[1] which was dissolved in 1.5 mL DMSO and 50 µL DIPEA. TSTU (21.8 mg, 72.4 µmol, 1.0 equiv.) was added in one portion and the mixture was incubated for 30 min before 32.0 mg (86.8 µmol, 1.2 equiv.) of Fmoc-Lys-OH (**25**) was added in one portion. The reaction mixture was incubated for another hour before it was quenched by addition of 50 µL HOAc and subjected to RP-HPLC to obtain 26 mg (32.2 µmol) of the desired product as a blue powder after lyophilization in 45% yield.

**HRMS** (ESI): calc. for $C_{53}H_{61}N_4O_5$ $[M]^+$: 833.4636, found: 833.4639.

**1.40. 1-(6-(((*S*)-5-Amino-6-((2-((4-((*E*)-(4-((5*S*,7*S*)-7-((*tert*-butoxycarbonyl)amino)-5,7-dicarboxyheptanamido)phenyl)diazenyl)phenyl)amino)-2-oxoethyl)amino)-6-oxohexyl)amino)-6-oxohexyl)-3,3-dimethyl-2-((1*E*,3*E*)-5-((*Z*)-1,3,3-trimethylindolin-2-ylidene)penta-1,3-dien-1-yl)-3*H*-indol-1-ium (28)**

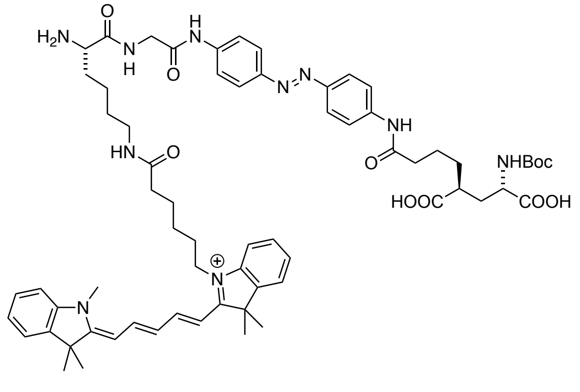

**28** was prepared according to general procedure 1.4.

**HRMS** (ESI): calc. for $C_{66}H_{85}N_{10}O_{10}$ $[M]^+$: 1177.6445, found: 1177.6452.

**1.41. 1-((*S*)-1-Amino-41-((2-((4-((*E*)-(4-((5*S*,7*S*)-7-((*tert*-butoxycarbonyl)amino)-5,7-dicarboxyheptanamido)phenyl)diazenyl)phenyl)amino)-2-oxoethyl)carbamoyl)-39,47-dioxo-3,6,9,12,15,18,21,24,27,30,33,36-dodecaoxa-40,46-diazadopentacontan-52-yl)-3,3-dimethyl-2-((1*E*,3*E*)-5-((*Z*)-1,3,3-trimethylindolin-2-ylidene)penta-1,3-dien-1-yl)-3*H*-indol-1-ium (29)**

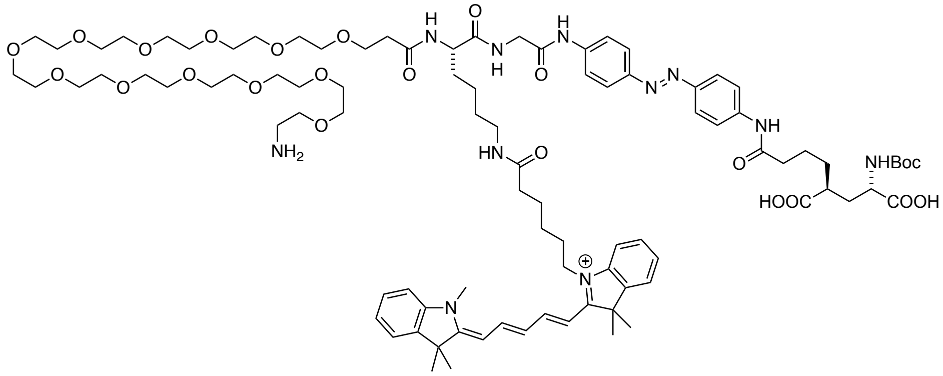

**29** was prepared according to general procedure 1.4 with the first step conducted at 50 °C.

**HRMS** (ESI): calc. for $C_{93}H_{139}N_{11}O_{23}$ $[M+H]^{2+}$: 889.5033, found: 889.5037.

**1.42. 1-((*S*)-1-(4-(((2-Amino-9*H*-purin-6-yl)oxy)methyl)phenyl)-49-((2-((4-((*E*)-(4-((5*S*,7*S*)-7-((*tert*-butoxycarbonyl)amino)-5,7-dicarboxyheptanamido)phenyl)diazenyl)phenyl)amino)-2-oxoethyl)carbamoyl)-3,7,47,55-tetraoxo-11,14,17,20,23,26,29,32,35,38,41,44-dodecaoxa-2,8,48,54-tetraazahexacontan-60-yl)-3,3-dimethyl-2-((1*E*,3*E*)-5-((*Z*)-1,3,3-trimethylindolin-2-ylidene)penta-1,3-dien-1-yl)-3*H*-indol-1-ium (30)**

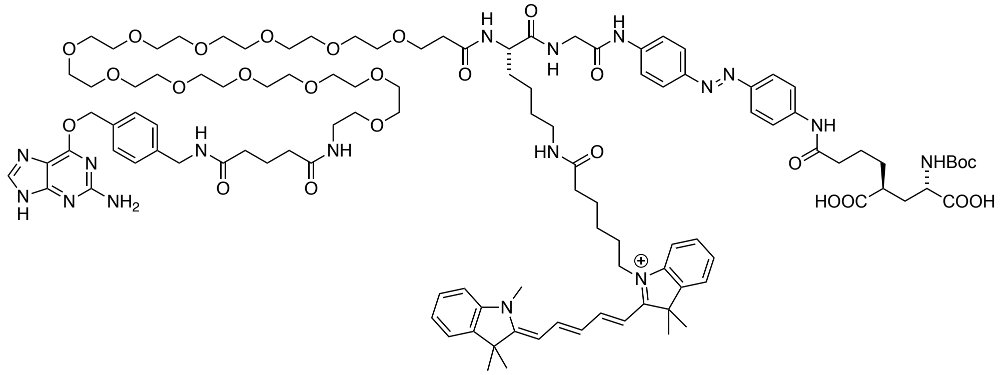

**30** was prepared according to general procedure 1.4, with **BG-COOSu** prepared according to general procedure 1.3, without the addition of piperidine.

**HRMS** (ESI): calc. for $C_{111}H_{158}N_{17}O_{24}$ $[M+2H]^{3+}$: 715.3859, found: 715.3859.

**1.43. 1-((*S*)-49-((2-((4-((*E*)-(4-((5*S*,7*S*)-7-Amino-5,7-dicarboxyheptanamido)phenyl)diazenyl)phenyl)amino)-2-oxoethyl)carbamoyl)-1-(4-(((2-amino-9*H*-purin-6-yl)oxy)methyl)phenyl)-3,7,47,55-tetraoxo-11,14,17,20,23,26,29,32,35,38,41,44-dodecaoxa-2,8,48,54-tetraazahexacontan-60-yl)-3,3-dimethyl-2-((1*E*,3*E*)-5-((*Z*)-1,3,3-trimethylindolin-2-ylidene)penta-1,3-dien-1-yl)-3*H*-indol-1-ium (BG-Cy5-AG$_{12}$)**

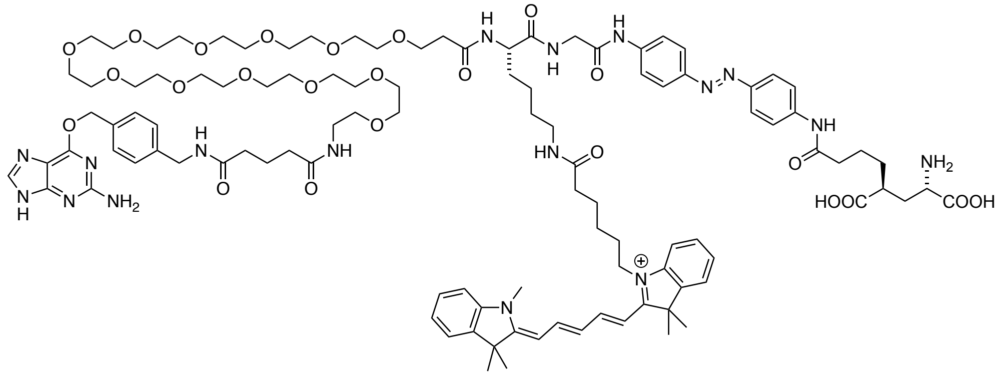

**BG-Cy5-AG$_{12}$** was prepared according to general procedure 1.6.

**HRMS** (ESI): calc. for $C_{106}H_{150}N_{17}O_{24}$ $[M+2H]^{3+}$: 682.0351, found: 682.0354.

#### 2. *References*

1. Ueno, Y.; Jose, J.; Loudet, A.; Perez-Bolivar, C.; Anzenbacher, P., Jr.; Burgess, K., Encapsulated energy-transfer cassettes with extremely well resolved fluorescent outputs. *J Am Chem Soc* **2011,** *133*, 51-5.
